# Identification and Evaluation of dibasic piperidines as novel cell wall inhibitors against *Mycobacterium tuberculosis*

**DOI:** 10.64898/2026.01.30.702510

**Authors:** Claire Naylor, Gareth A Prosser, Tracy Bayliss, Lila Berle, Joshua B Wallach, Heather Kim, Rodrigo Aguilera Olvera, Stephen Thompson, Thomas R Ioerger, Laura Simpson, Ruth Casanueva, Laura Guijarro-Lopez, Kevin D. Read, Paul G Wyatt, Dirk Schnappinger, Clifton E Barry, Simon R Green, Helena I Boshoff, Laura A T Cleghorn

## Abstract

Globally, *Mycobacterium tuberculosis* remains a significant disease burden. Although effective treatment regimens exist, drug resistance continues to emerge. This clinical resistance, combined with side effects and protracted treatment times from the current front-line therapies, means there is a need to identify novel agents to combat this disease. Here we report on a new chemical series, identified by whole-cell phenotypic growth inhibition screening that demonstrates significant activity across multiple media. Mode of action studies indicate that this series targets the same biological pathway as Ethambutol (EMB), a drug used in the current frontline treatment of tuberculosis. Screening selected analogues against clinical isolates, resistant to EMB, demonstrated differential sensitivity both across the molecules and against the different specific resistant mutations. The data obtained suggests that this series has potential to be developed into a viable, alternative to EMB.

**TOC figure:** 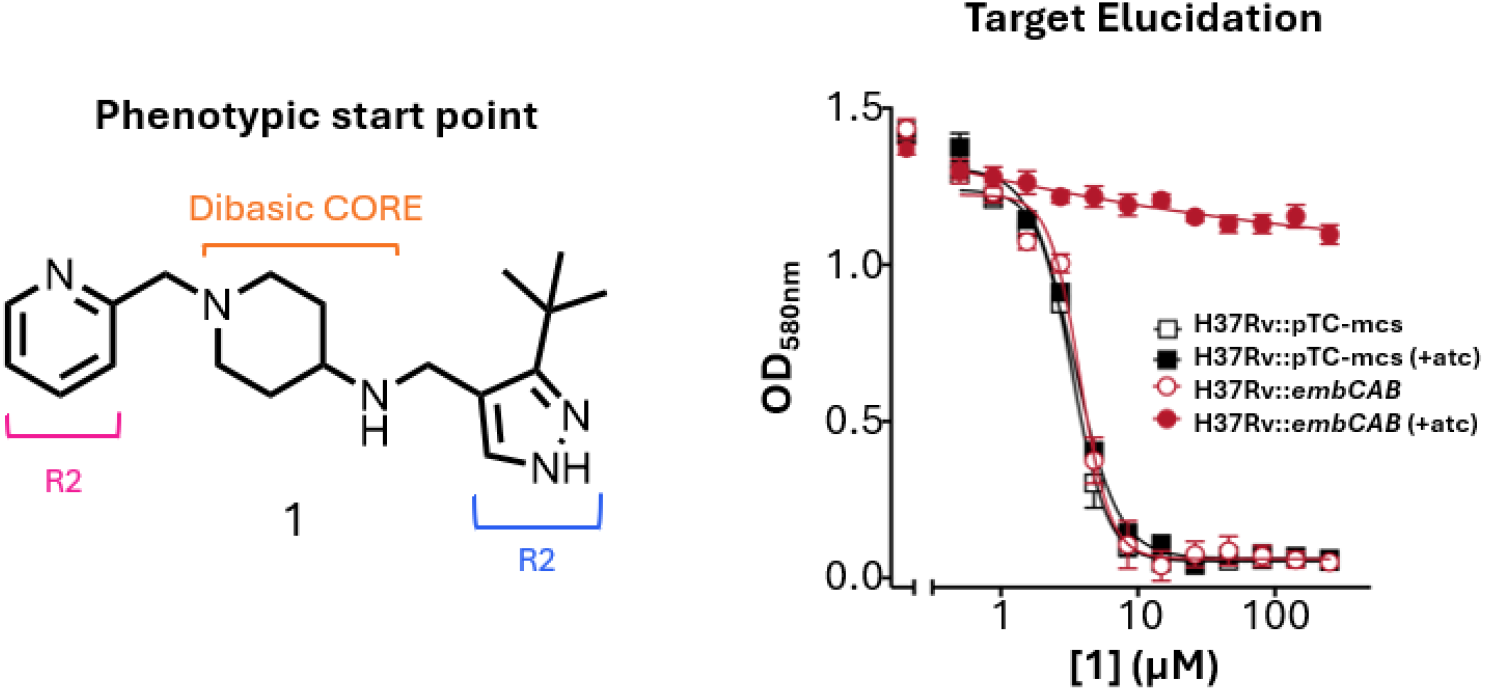

## Introduction

Tuberculosis (TB) is an infectious disease caused by the bacterium *Mycobacterium tuberculosis*. Treating TB during its pulmonary stage is crucial to reduce its transmission. Despite being preventable and curable, it’s estimated that TB infected 10.7 million people in 2023, with approximately 1.23 million deaths attributed to the disease^1^. The standard regimen for drug-susceptible TB necessitates a minimum of 6 months of treatment, initially for two months using a four-drug regimen isoniazid (INH), rifampicin, pyrazinamide and ethambutol (EMB) and then a further 4 months treatment with isoniazid and rifampicin^2^. The extended duration of treatment is necessary to target the diverse physiological states of *M. tuberculosis* present during infection, including actively replicating, slowly replicating, and dormant bacilli^3^. The length and complexity of the treatment can be challenging for both individuals and public health sector resources; when treatment is under less-than-ideal conditions drug resistant forms of the disease can emerge that are even more challenging to treat^4^. The growing resistance to frontline drugs underscores the critical need for new TB medications with a shorter treatment duration^3^. To tackle this challenge, a whole-cell phenotypic growth inhibition screening campaign targeting *M. tuberculosis* was performed utilising a chemically diverse set of 20,000 compounds. Here we present the design and hit assessment around a novel phenotypic hit, elucidation of the mechanism of action and lessons learnt for future studies with di-basic starting points for TB drug discovery.

## Results and Discussion

*Hit Discovery:* The MMV Diversity Library (40,000 compounds) is a diverse screening library of lead-like properties co-created by the Medicine for Malaria Venture and the Dundee Drug Discovery Unit selected from the Enamine library. The molecules were selected from the 1.5 million compounds that were commercially available from the Enamine library. The strategy for this collection was to provide a diverse lead-like library that would be available for screening against ‘new malaria’ assays, as well as other neglected disease assays. High-throughput phenotypic growth inhibition screening with *M. tuberculosis* was performed in two standard media as previously described^5^. From this screen, a hit compound (**1**) emerged as an attractive chemical starting point for further exploration (**Figure 1**). For an initial hit it had reasonable potency and good drug-like properties, such as low intrinsic clearance, excellent solubility and a reasonable cytotoxicity selectivity window. In addition, **1** had an acceptable *in vivo* pharmacokinetic (PK) profile with high volume of distribution, moderate oral bioavailability and unexpectedly high *in vivo* clearance (**Figure 1**).

**Figure 1.**
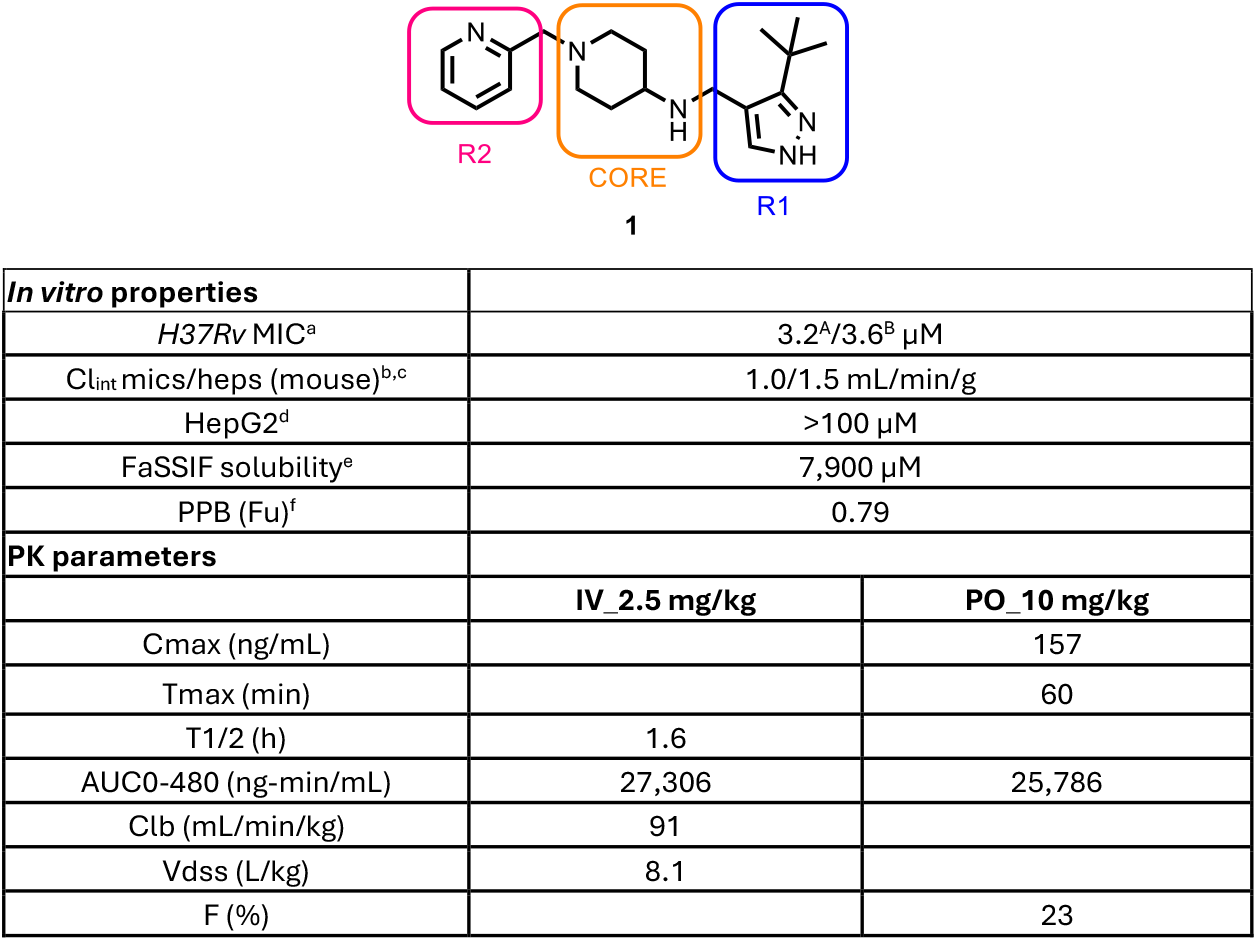
Profile of initial hit compound 1. ^a^MIC is the minimum concentration required to inhibit the growth of *M. tuberculosis* (H37Rv) in liquid culture by 90% compared to untreated control (MIC^A^ 7H9/DPPC/CAS/Tx media, MIC^B^ 7H9/GLU/CAS/Tx media); ^b^Intrinsic clearance (Cli) using CD1 mouse liver microsomes(female); ^c^Intrinsic clearance (Cli) using mouse hepatocytes. ^d^HepG2 inhibitory concentration (IC_50_) is the concentration required to inhibit growth of HepG2 cells by 50%; ^e^Solubility in simulated fasted intestinal fluid (FaSSIF); ^f^Plasma protein binding (Fraction unbound); ^g^Pharmacokinetic parameters for **1**, dosed at 2.5mg/kg IV and 10 mg/kg PO in mice.

Many potent hits identified via *M. tuberculosis* phenotypic screening target membrane-bound proteins, involved in cell wall biosynthesis^6^. The most frequently used TB treatment includes isoniazid (INH) and ethambutol (EMB), two front-line drugs which target cell wall biosynthesis, so this pathway is a clinically validated area for anti-tubercular drugs. To assess whether **1** was inhibiting cell wall biosynthesis, it was evaluated against a P*iniB*-LUX strain, in which the *luxCDABE* operon, encoding a bacterial luciferase is present downstream of the *iniBAC* promoter. This strain was established as a bioluminescent reporter for compounds that disrupt cell wall biosynthesis^7-8^. A clear bioluminescence signal was detected when **1** was evaluated in the P*iniB*-LUX assay, as was also seen for known cell wall inhibitor controls (**Figure S1**). There are several compounds currently in development that target some of the most “frequent hitter” cell wall biosynthesis enzymes, such as DprE1, MmpL3 and Pks13^9-10^. As such, a decision was made not to pursue more compounds against these enzymes, until the clinical studies of agents already in development had progressed to the extent where it could be seen if they were clinically effective.

To evaluate whether compound **1** interacted with any of these promiscuous cell wall targets, it was assessed using hypomorph strains with regulated expression of specific cell wall biosynthetic genes^5^. In these strains the expression of the targeted gene(s), e.g., *dprE1* and *dprE2* in *dprE12*-TetON1 and *mmpL3* in *mmpL3*-TetON, is (are) downregulated in the absence of anhydrotetracycline (atc). As a result, if a compound inhibited the target in question, it was expected that there would be a clear differential in sensitivity to the compound ±atc. In contrast to the control compounds targeting DprE1, MmpL3, Pks13 or InhA, **1** had no differential effect on growth in any of the hypomorph strains (**Figure 2**), implying that **1** was not targeting one of the cell wall biosynthesis proteins for which compounds are already in clinical use or under clinical evaluation and was therefore worthy of further investigation.

**Figure 2:**
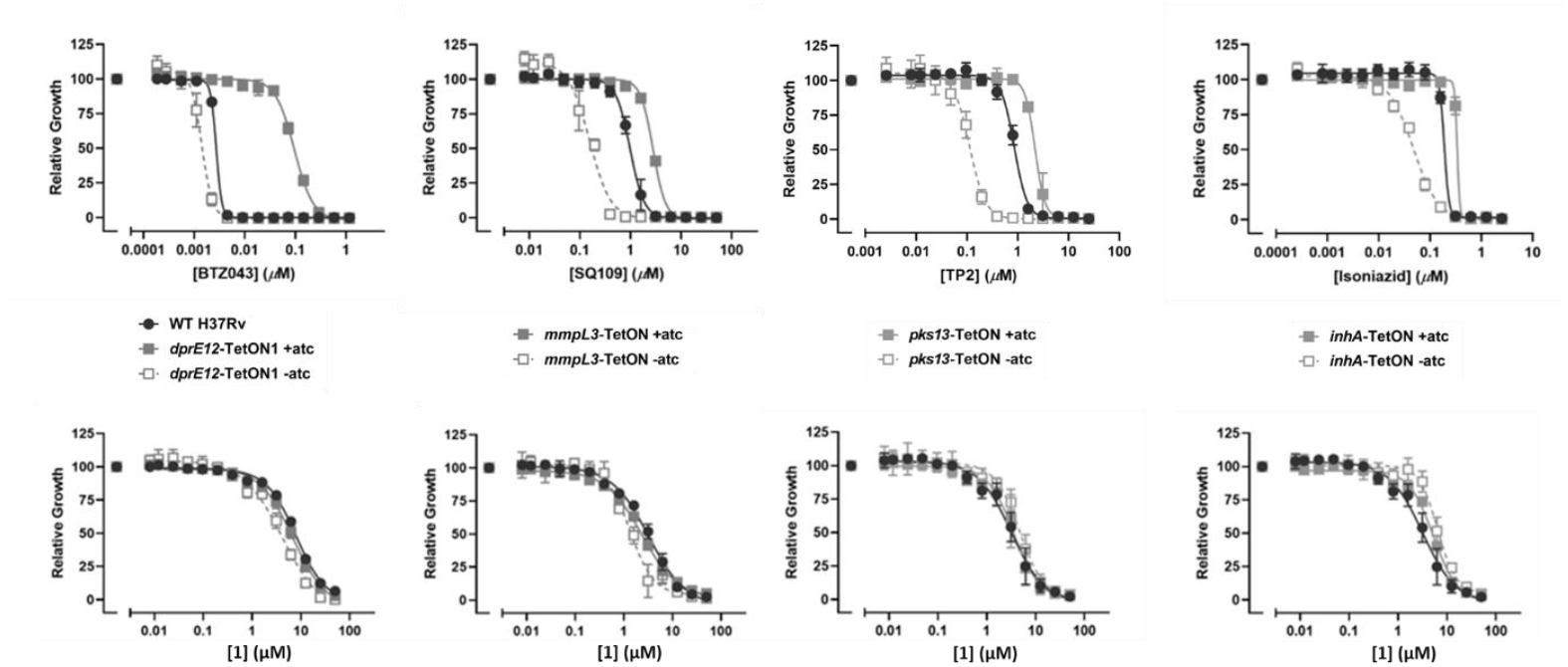
Studies to explore the cellular mechanism of action. Compound **1** was profiled against hypomorph strains of specific cell wall biosynthetic genes looking at the impact of TetON overexpression on sensitivity to **1** ± anhydrotetracycline (ATc), BTZ043, SQ109, TP2 and Isoniazid were used as the control compounds for DprE1, Mmpl3, Pks13 and InhA respectively, WT H37Rv is included in each plot for comparison. All graphs are representative data from one of two independent experiments each run in triplicate presented as mean values ± standard deviation.

**Figure 3:**
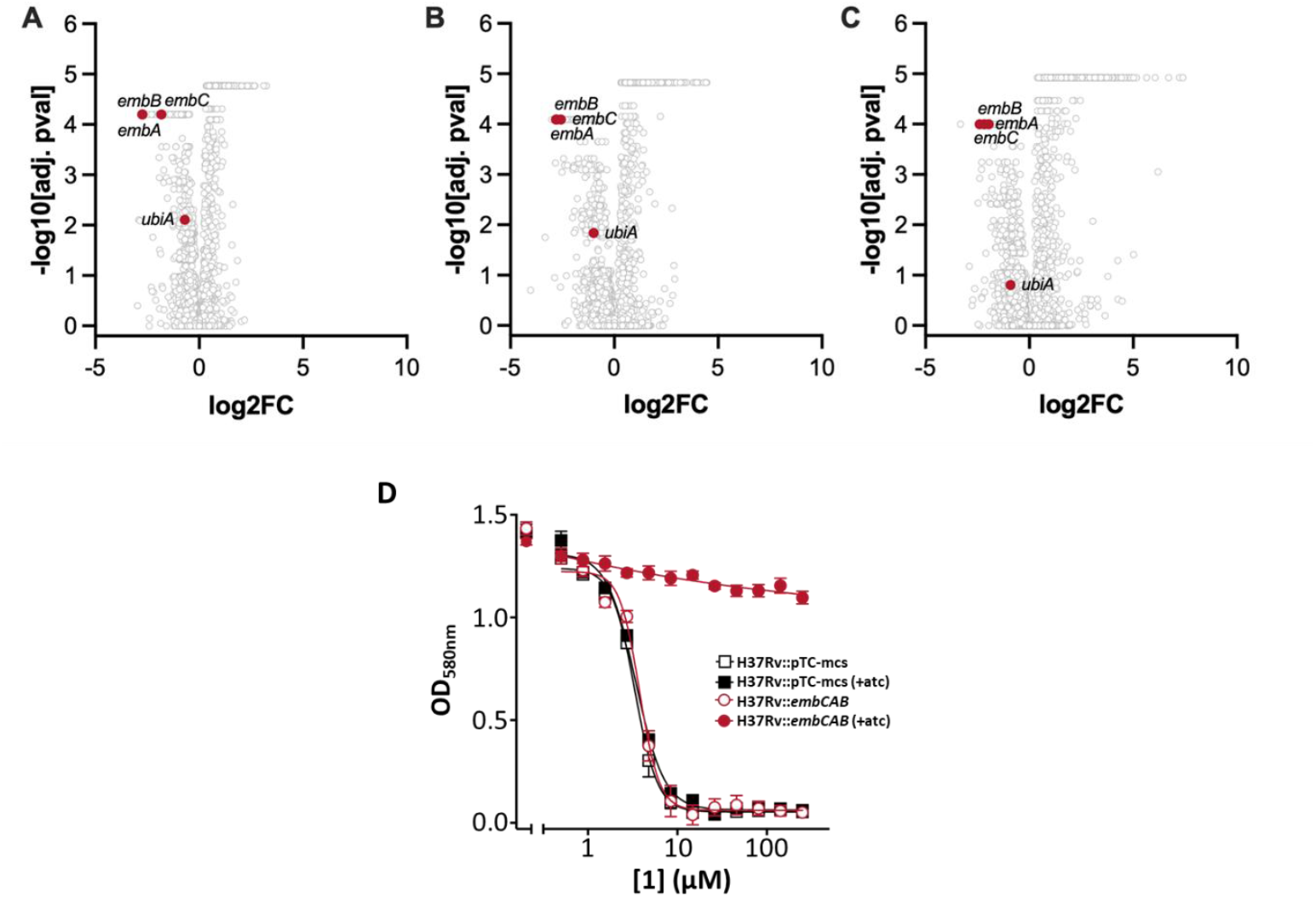
Compound 1 targets the arabinogalactan biosynthesis pathway. Pooled CRISPRi drug profiling(**A-C**), volcano plots showing log_2_ fold change (log2FC) values vs the log_10_ probability at different concentrations of **1**. Experiments were done in duplicate at three different concentrations (0.625x, 0.125x and 0.25x MIC A-C respectively), (**D**) Overexpression of the *embCAB* operon eliminates growth inhibition of **1**. Strains were grown ±atc, data are averages of two or three cultures and representative of two independent experiments.

### Structure activity relationship (SAR)

Initial exploration of the SAR of **1** was divided into three areas, R_1_, the core, and R_2_ (**Figure 1**). The initial SAR investigation focussed on examining the scope of changes that were possible at R_1_. Unfortunately, all R_1_ changes strongly reduced potency and therefore are only described in the supplementary information (**Table S1:** compounds **2 - 11**). As a result of the extremely tight SAR around the R_1_ group, it was held constant in all further molecules, and the SAR focus was shifted to exam modifications to the central amino piperidine core region to identify opportunities that could enhance MIC potency (**Table 1**).

**Table 1.**
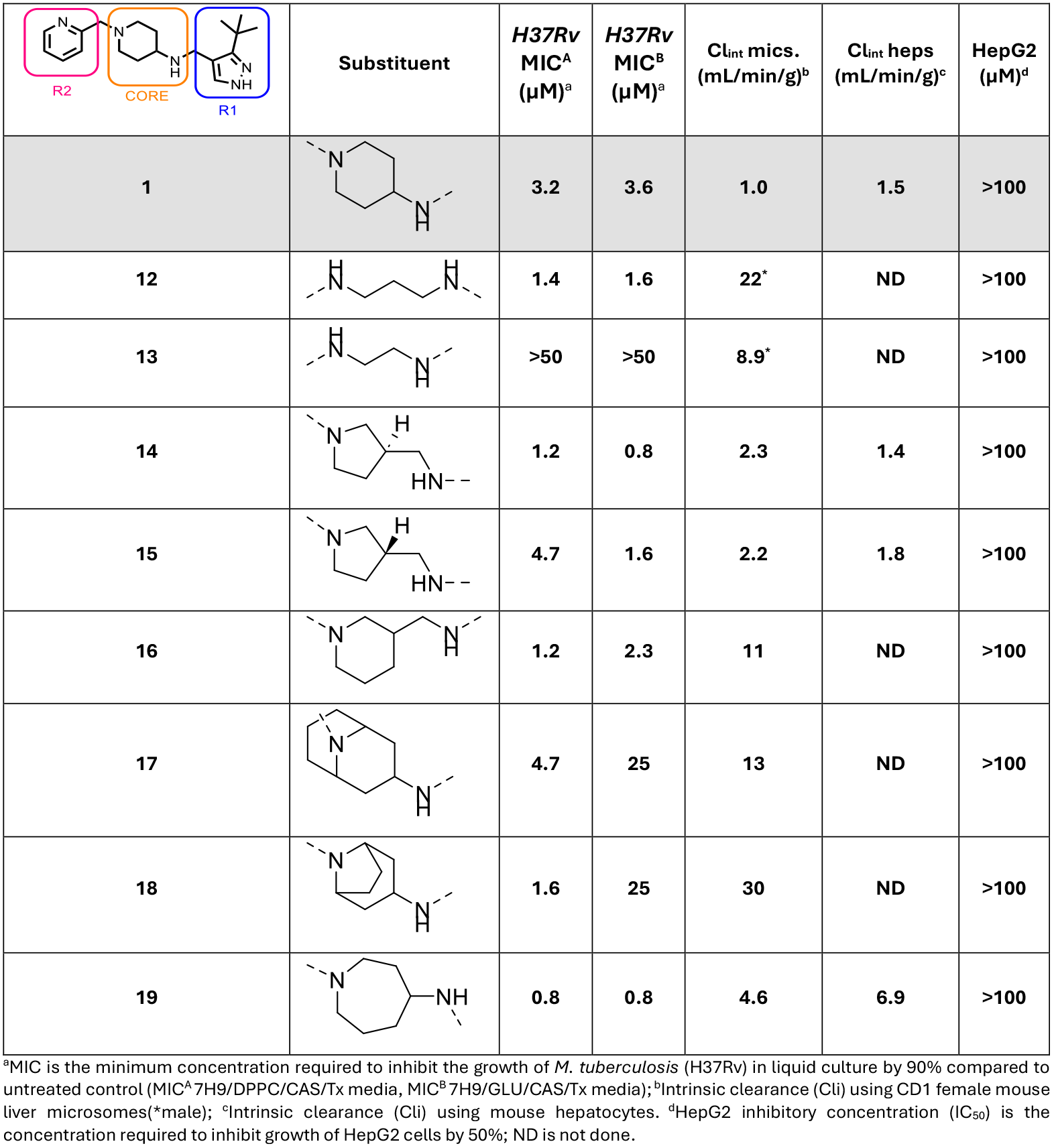
Core Modifications.

| | Substituent | H37Rv<br>MIC<sup>A</sup><br>(μM)<sup>a</sup> | H37Rv<br>MIC<sup>B</sup><br>(μM)<sup>a</sup> | Cl<sub>int</sub> mics.<br>(mL/min/g)<sup>b</sup> | Cl<sub>int</sub> heps<br>(mL/min/g)<sup>c</sup> | HepG2<br>(μM)<sup>d</sup> |
| --- | --- | --- | --- | --- | --- | --- |
| **1** |  | 3.2 | 3.6 | 1.0 | 1.5 | >100 |
| **12** |  | 1.4 | 1.6 | 22* | ND | >100 |
| **13** |  | >50 | >50 | 8.9* | ND | >100 |
| **14** |  | 1.2 | 0.8 | 2.3 | 1.4 | >100 |
| **15** |  | 4.7 | 1.6 | 2.2 | 1.8 | >100 |
| **16** |  | 1.2 | 2.3 | 11 | ND | >100 |
| **17** |  | 4.7 | 25 | 13 | ND | >100 |
| **18** |  | 1.6 | 25 | 30 | ND | >100 |
| **19** |  | 0.8 | 0.8 | 4.6 | 6.9 | >100 |

<sup>a</sup>MIC is the minimum concentration required to inhibit the growth of *M. tuberculosis* (H37Rv) in liquid culture by 90% compared to untreated control (MIC<sup>A</sup> 7H9/DPPC/CAS/Tx media, MIC<sup>B</sup> 7H9/GLU/CAS/Tx media); <sup>b</sup>Intrinsic clearance (Cl<sub>i</sub>) using CD1 female mouse liver microsomes(\*male); <sup>c</sup>Intrinsic clearance (Cl<sub>i</sub>) using mouse hepatocytes. <sup>d</sup>HepG2 inhibitory concentration (IC<sub>50</sub>) is the concentration required to inhibit growth of HepG2 cells by 50%; ND is not done.

The aminopiperidine could be replaced with diaminopropyl (**12**) with modest gains in MIC potency, however in stark contrast, diaminoethyl (**13**) resulted in a complete loss of activity, highlighting the distance between the two nitrogen’s was key to the observed potency. Consequently, further core modification focused on cores with at least three carbon spacers between the diamine functionalities. Substituted pyrrolidine (**14, 15**) and piperidine methanamine (**16**) were equipotent and showed a similar profile to the original hit (**1**). However, the 3-substituted piperidine was found to have a higher microsomal turnover than the original hit (**1**). Interestingly, rigidifying the piperidine of **1** with 3- or 2**-** carbon bridges (**17, 18**) retained MIC activity in media supporting beta-oxidation with dipalmitoyl phosphatidylcholine (DPPC) as carbon source (media A) but lost potency in glucose media supporting glycolytic metabolism (media B). Ring expansion of the piperidine to azepane (**19**) improved potency but was detrimental to the intrinsic clearance most likely due to increased lipophilicity.

For the R_2_ region, a range of substitution patterns and deletion of the pyridyl nitrogen were investigated (**Table 2**) and compared to **1**. Removal of the pyridyl nitrogen to the unsubstituted phenyl (**20**) was tolerated in DPPC media but had a 3-fold potency loss in glucose media. To explore the effect of adding substituents to the pyridine ring, a mono methoxy group was systematically substituted at each position. A 3-methoxy (**21**) showed a 10-fold improvement in MIC activity with comparable clearance in mouse hepatocytes (**Table 2**). In contrast, 4-,5- and 6-methoxy substituents (**22 - 24**) had a detrimental effect on MIC and were not pursued further. As the 3-position of the pyridyl ring was identified as a promising vector to improve the antibacterial activity (**21**) the 3-methyl, fluoro and hydroxy substituents (**25 - 27**) were investigated but showed no gain in potency compared to **21**. Extending the 3-methoxy to 3-isopropoxy (**28**) or 3-ethoxy (**29**) showed no improvement in either potency or microsomal turnover. Replacement of the pyridine with benzimidazole (**30**) was equipotent to **1** and displayed a similar *in vitro* clearance profile. Truncation to imidazole (**31**) yielded excellent MIC activity and exhibited very low microsomal clearance. However, **31** was subsequently confirmed as a potent CYP inhibitor (CYP2C19 & CYP2D6 IC_50_ = 0.7 µM & 0.6 µM, respectively) and therefore not pursued further. Compound **32** was prepared to combine the optimal core (azepan-4-amine, **19**) with the best R_2_ substituent (3-methoxypyridine, **21**). Unfortunately, no further gain in MIC potency was observed and the molecule had high microsomal turnover (**Table 2**).

**Table 2.**
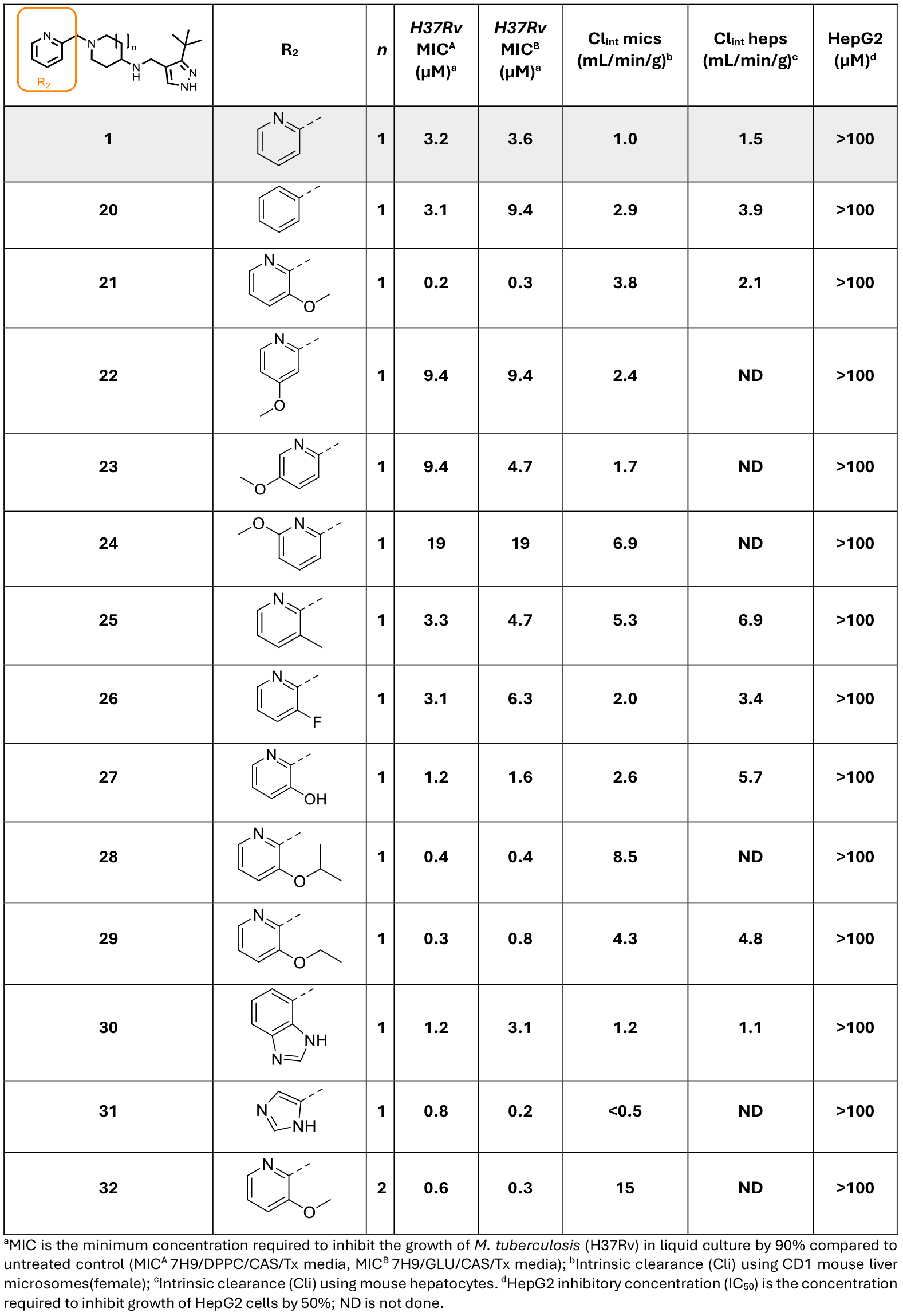
SAR of R_2_ substituent.

|  | R<sub>2</sub> | n | H37Rv<br>MIC<sup>A</sup><br>(μM)<sup>a</sup> | H37Rv<br>MIC<sup>B</sup><br>(μM)<sup>a</sup> | Cl<sub>int</sub> mics<br>(mL/min/g)<sup>b</sup> | Cl<sub>int</sub> heps<br>(mL/min/g)<sup>c</sup> | HepG2<br>(μM)<sup>d</sup> |
| --- | --- | --- | --- | --- | --- | --- | --- |
| **1** |  | 1 | 3.2 | 3.6 | 1.0 | 1.5 | >100 |
| **20** |  | 1 | 3.1 | 9.4 | 2.9 | 3.9 | >100 |
| **21** |  | 1 | 0.2 | 0.3 | 3.8 | 2.1 | >100 |
| **22** |  | 1 | 9.4 | 9.4 | 2.4 | ND | >100 |
| **23** |  | 1 | 9.4 | 4.7 | 1.7 | ND | >100 |
| **24** |  | 1 | 19 | 19 | 6.9 | ND | >100 |
| **25** |  | 1 | 3.3 | 4.7 | 5.3 | 6.9 | >100 |
| **26** |  | 1 | 3.1 | 6.3 | 2.0 | 3.4 | >100 |
| **27** |  | 1 | 1.2 | 1.6 | 2.6 | 5.7 | >100 |
| **28** |  | 1 | 0.4 | 0.4 | 8.5 | ND | >100 |
| **29** |  | 1 | 0.3 | 0.8 | 4.3 | 4.8 | >100 |
| **30** |  | 1 | 1.2 | 3.1 | 1.2 | 1.1 | >100 |
| **31** |  | 1 | 0.8 | 0.2 | <0.5 | ND | >100 |
| **32** |  | 2 | 0.6 | 0.3 | 15 | ND | >100 |

<sup>a</sup>MIC is the minimum concentration required to inhibit the growth of *M. tuberculosis* (H37Rv) in liquid culture by 90% compared to untreated control (MIC<sup>A</sup> 7H9/DPPC/CAS/Tx media, MIC<sup>B</sup> 7H9/GLU/CAS/Tx media); <sup>b</sup>Intrinsic clearance (Cl<sub>i</sub>) using CD1 mouse liver microsomes(female); <sup>c</sup>Intrinsic clearance (Cl<sub>i</sub>) using mouse hepatocytes. <sup>d</sup>HepG2 inhibitory concentration (IC<sub>50</sub>) is the concentration required to inhibit growth of HepG2 cells by 50%; ND is not done.

### MOA evaluation

In parallel with the SAR exploration of the series, further investigations were conducted to elucidate the precise cell wall mechanism of action. Spontaneous resistant mutants were generated through growth on solid media containing **1**. The frequency of resistance (∼5 times MIC) was high, 2.5 x10^-6^. Four individual resistant isolates were obtained and whole genome sequencing revealed three distinct single nucleotide polymorphisms (SNPs) in the genes *ubiA, embB* and *embC* (**Table S2**). Mutations in these genes have been linked to phenotypic resistance to the front-line antitubercular drug EMB^11-13^, which inhibits essential arabinosyltransferases (EmbA, EmbB, and EmbC) involved in the biosynthesis of the arabinogalactan layer of the mycobacterial cell wall, as well as the glycolipid lipoarabinomannan^14^. UbiA catalyses the formation of decaprenylphosphoryl-5-phosphoribose^15^, a precursor for the final arabinose donor molecule used by EmbB and EmbC, decaprenylphosphoryl-D-arabinose.

As a second approach for target identification, we applied a genome-wide CRISPRi approach^16-17^. Following treatment with **1**, strains with reduced expression of *embA, embB* or *embC* were consistently among the topmost sensitized strains (**Figure 4A-C**). Silencing of *ubiA* also sensitized Mtb to growth inhibition by **1**, but the extent of sensitization was less pronounced than for e*mbCAB*. These results corroborated the resistant mutant data and supported the hypothesis that **1** was targeting the arabinogalactan biosynthesis pathway.

**Figure 4:**
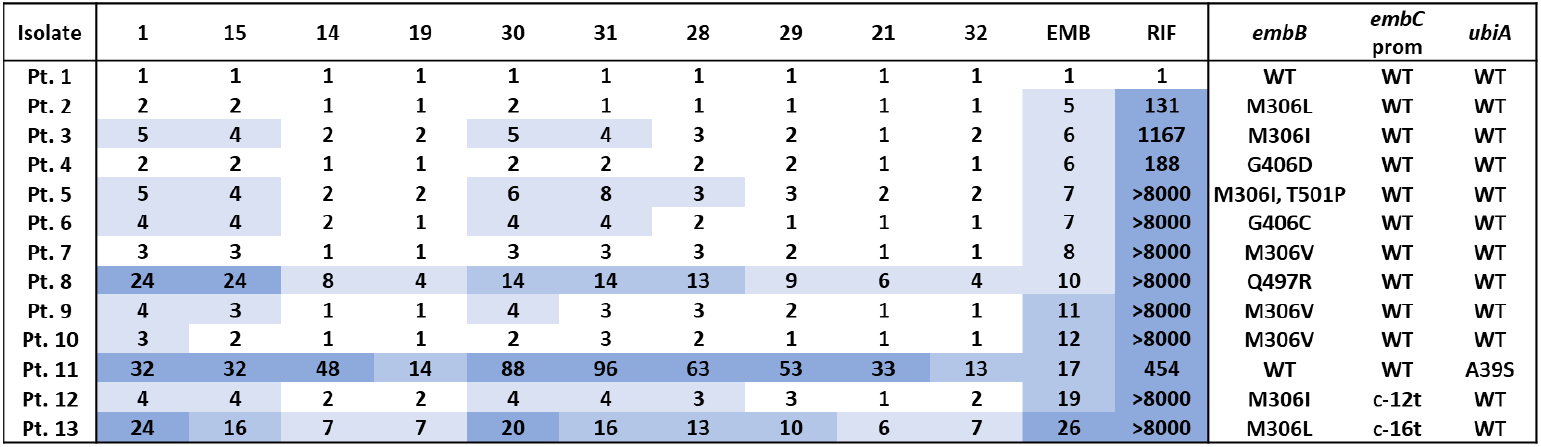
Profiling of selected inhibitors against clinical samples from EMB resistant patients. The values shown for each strain are the fold increase in MIC relative to the MIC of H37Rv.

To explore the involvement of the *embCAB* gene cluster in resistance to this series, a strain was constructed that overexpressed *embCAB* under the control of a TetON expression system; thus,in the presence of atc, expression of the *embCAB* genes was increased. When this strain (embCAB OE) was treated with **1**, in the absence of atc, the IC_50_ for growth was the same as H37Rv (**Figure 4D**). In the presence of atc, there was no impact on the growth inhibition of **1** on H37Rv; in contrast, no growth inhibition was observed for the embCAB OE strain after addition of atc (**Figure 4D**). Taken together, there are three lines of evidence demonstrating that this series, like EMB, is targeting the arabinogalactan biosynthesis pathway. Given that EMB is a frontline therapy currently used in the clinic, there is extensive information available about resistance mutations that are already circulating in the TB patient population. None of the 3 resistant mutants identified following treatment with **1** are listed in the WHO clinical mutation catalogue^18^, suggesting that they are not common mutations. However, the mutations in *embB* and *ubiA* are certainly present in regions of the molecule that contain clinically relevant resistant mutations^11-12, 18^. Since mutations in genes that might cause resistance to the series are already circulating in the patient population, this raised concerns about the viability of pursuing the series further. While there are some chemical similarities between this series and EMB, in that both compounds contain a di-basic centre, the SAR of the series also has some very clear differences, in that: a 2-carbon linker was not tolerated (**13**) and the *t*-butyl on R1 was key, yet a similar moiety is not present in EMB. Based on these chemical differences, it was possible that strains clinically resistant to EMB might not be impacted to the same extent for their sensitivity to this series. If this were the case, then there was still potential for further development. To explore this question, a representative set of ten compounds were tested against 13 clinical isolates, 12 of which contained known EMB resistance mutations (**Figure 4**).

As would be expected, the different mutations had differing effects on compound sensitivity. So, isolate 11 that contained an *ubiA* A39S point mutation was significantly resistant to all 10 molecules. Likewise, isolate 8 with a single Q497R point mutation in *embB*, and isolate 13 with a point mutation in *embB* (M306L) and an *embC* promoter mutation (c-16t) both showed increased resistance across the series. However, isolates containing single mutations in *embB* such as M306V or G406D, although causing resistance to EMB, had minimal impact on sensitivity to this series. Unexpectedly, some of the molecules appeared to be significantly less affected by the mutations than others. Notably, if the size of the core was expanded to a homopiperidine the extent of resistance was reduced (**19** vs **1**). Likewise, substituents at the 3-position on the pyridine ring also decreased resistance (**21/29** vs **1**). Moreover, when these two changes were combined (**32**) then the increase in resistance was significantly reduced for all the clinical isolates compared to both **1** and EMB. Thus, while there was some overlap in resistance, there were clear differences in sensitivity to some of the representatives of this series when compared to EMB.

*In vivo efficacy in a murine model of TB infection*. Efficacy studies in mice evaluate activity of inhibitors against the pathogen growing in macrophages since standard mouse models consist of cellular lesions, where most of the bacilli reside in macrophages^19^. Prior to progressing to *in vivo* studies, the potency of the compounds against *M. tuberculosis* growing in a macrophage cell line were evaluated. Compounds **1, 19** and **21** at 10 µM (3-, 13- and 40-fold MIC values) were superior to 1 μM Rifampicin in bacterial killing in the macrophage assay (**Figure 5A**). The intramacrophage activity of the cell wall inhibitors aligns with the known activity of cell wall inhibitors such as EMB against intracellular *M. tuberculosis*^*20*^; although the relative fold kill between these analogues was not correlated with *in vitro* MIC, likely reflecting differences in intraphagosomal penetration of the compounds. These results prompted the progression of the series to *in vivo* evaluation. It had been established previously that the initial hit **1** had an acceptable *in vivo* PK profile (**Figure 1**). Reviewing the properties of the analogues synthesised, although **19** appeared the most potent in the intramacrophage assay, it had unacceptable metabolic stability in hepatocytes (**Table 2**). Compound **21** demonstrated a >10-fold enhancement in MIC potency relative to **1**, while maintaining comparable *in vitro* clearance. Additionally, **21** maintained a high free fraction (FU = 0.87) and excellent FaSSIF solubility (6,800 µM) suggesting it would exhibit at least an equivalent PK profile compared to compound **1**. As such, **21** was selected for evaluation in a murine model of acute TB infection^21^. For an early-stage molecule, the efficacy obtained was very promising (**Figure 5B**). The bacterial load in the lungs was measured by qPCR and a 2.8 log_10_ reduction in DNA was achieved, approaching the maximum reduction possible in this model format (3.0 log_10_ reduction). Compound **21** demonstrated efficacy superior to the reference compound moxifloxacin, providing encouraging *in vivo* proof-of-concept for the chemical series.

**Figure 5:**
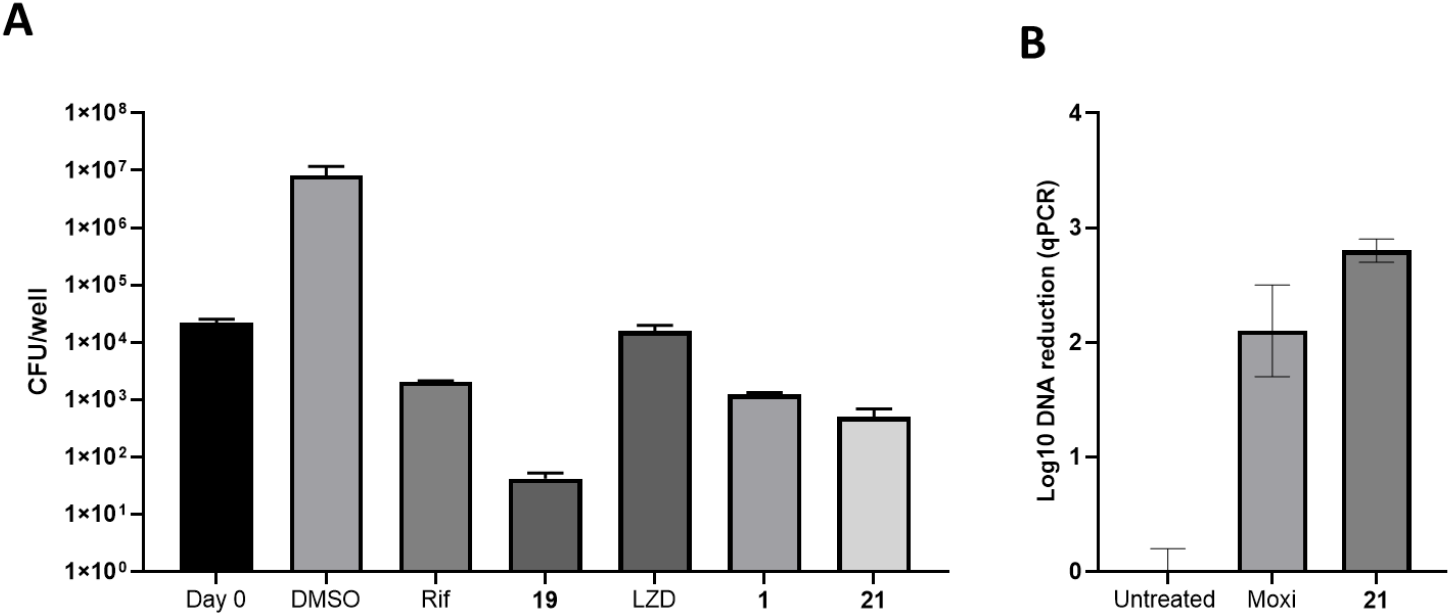
Intramacrophage and *in vivo* efficacy. **A**. Activity of compounds **1, 19** and **21** against *M. tuberculosis* grown in J774 macrophages. J774 macrophages were infected with *M. tuberculosis* and treatment of triplicate wells initiated 24 hours after infection. Cells were treated with **1, 19** and **21** (all 10 µM) along with controls Rifampicin (1 µM) and linezolid (25 µM). Cells were lysed after 7 days treatment and bacterial burdens enumerated by plating of dilutions of the homogenates on Middlebrook 7H11/OADC agar. **B**. *In vivo* efficacy of compound **21** (200 mg/kg) and control compound moxifloxacin (30 mg/kg) tested in an acute murine model of *M. tuberculosis* infection in C57BL/6 mice. Oral dosing started 1 day after infection and lasted for 8 days; each control group consisted of 4 mice and the **21** treatment group was 2 mice. The effect on bacterial load in mouse lungs was assessed using qPCR to monitor bacterial DNA in lung homogenates compared to the untreated group.

## Conclusion

Screening directly for the inhibition of growth of *M. tuberculosis* ensures that hits are cellularly active and represent potential start points for drug discovery programs. Here, the SAR of an amino piperidine series has been examined. The series displayed excellent potency against *M. tuberculosis*, and a key analogue was advanced for assessment in a proof-of-concept acute model of infection. Remarkably, especially for an early hit molecule, a profound reduction in bacterial load was observed within the lungs. In parallel, mode of action studies found the series to be targeting the same pathway as EMB, a key drug used in first-line TB treatment. Interestingly, when screened against a range of clinical isolates resistant to EMB, not all analogues from the series displayed the same resistance profile. This suggests that there may be scope for further SAR to identify molecules that limit potential preexisting clinical resistance. The main EMB mutational hotspots are in the *embB* gene, which appear to be associated with direct binding of EMB to the EmbB protein e.g. via residue M306^22-23^. These mutations had limited impact on the sensitivity to the current series. Other EMB resistance mutations arise in UbiA, which is involved in the synthesis of DPA (decaprenylphosphoryl-β-D-arabinose) a substrate for the Emb family of arabinosyltransferases. Mutations in UbiA that lead to EMB resistance are believed to be due to increased production of DPA directly competing with EMB^24-25^. Although the one UbiA mutation evaluated was still resistant to this series, there was dramatic variability in the impact on sensitivity from a 13 to 96-fold increase. Additional SAR rounds could reduce the EMB cross resistance further. Given this is a clinically validated pathway, developing an inhibitor with enhanced selectivity against EMB resistant isolates would be highly valuable.

## Methods

**Scheme 1:**
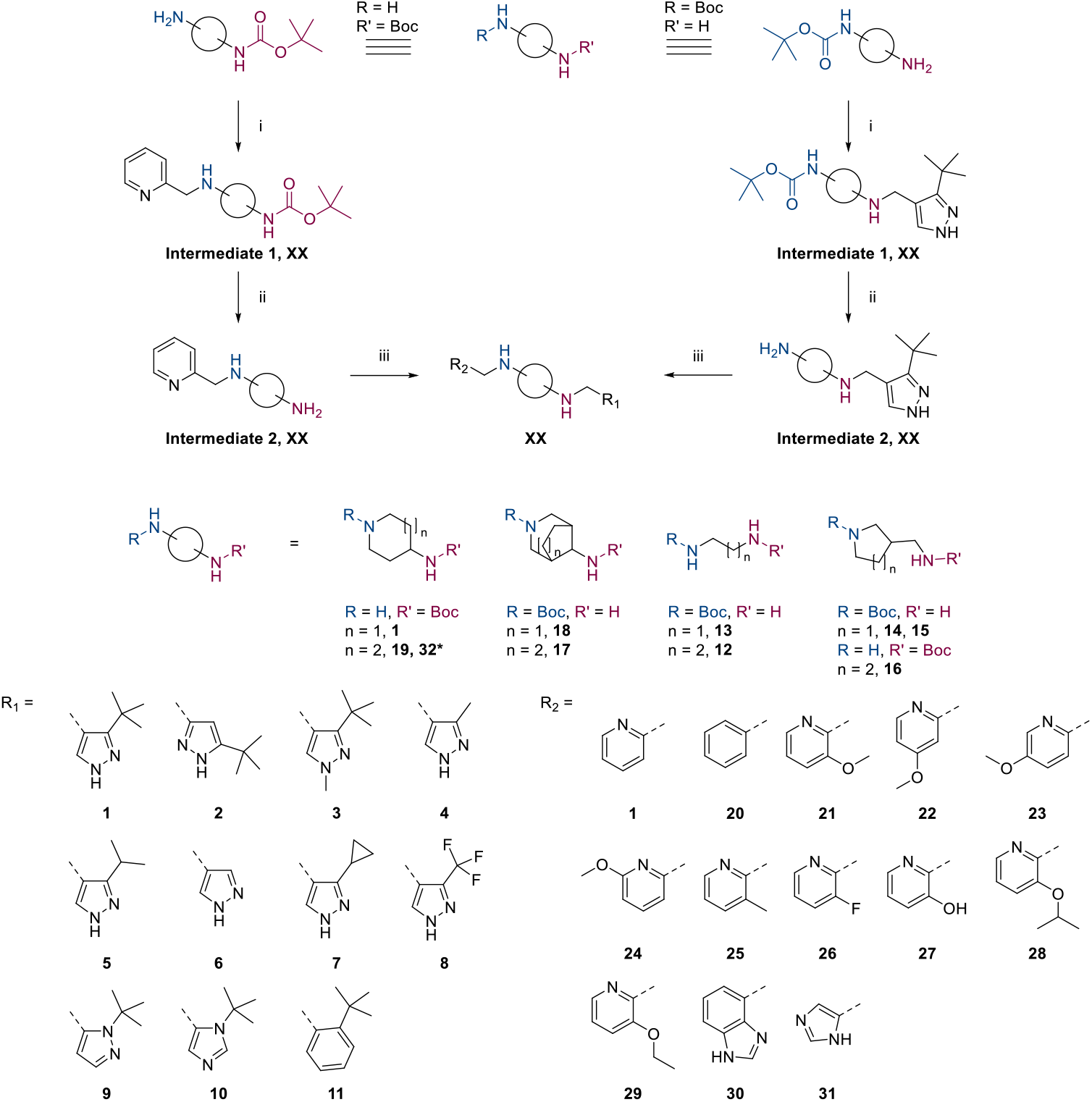
General synthetic scheme, 1-31. *Made on 3-methoxypyridine (i) Aldehyde, AcOH, MeOH, NaBH_3_CN/AcONa or NaBH(OAc)_3_, rt (ii) HCl in dioxane, ACN or MeOH, rt or 0°C (iii) Aldehyde, AcOH, NaBH(OAc)_3_ or NaBH_3_CN/AcONa, THF or EtOH or DCM or MeOH, mol. sieves, rt

### Synthetic Routes

Exploring the SAR of the hit **1** utilized a common reaction route varying the di-basic central core (**Scheme 1**) starting with either commercially available R or R’ amine Boc protected. The corresponding R_1_ or R_2_ substituent was added via reductive amination, the Boc group removed to afford the key amine building block for final step modification via a second reductive amination adding the corresponding R_1_ or R_2_ substituent to synthesize **1**-**31**. While most of the pyrazole building blocks were sourced from commercial suppliers, those used in the synthesis of **2, 3, 5**, and **8** were prepared following established literature procedures. The synthesis of **28** was achieved from alkylation of **27** as shown in **Scheme 2**.

**Scheme 1:**
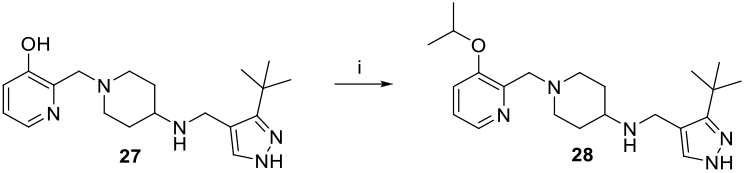
Synthetic schedule for 3-isopropoxy pyridine, 28. (i) K_2_CO_3_, isopropyl bromide, DMF, rt.

### Compound synthesis

Full experimental procedures can be found in the Supplementary information section

### Non-Chemistry methods

#### Screening

*M. tuberculosis* H37Rv or derivatives thereof were used for all experiments. Single point (20 µM) high throughput screening was performed on an ∼40,000 compound library provided by Medicines for Malaria Venture. HTS was performed in two liquid media 7H9/Glu/BSA/Tx (4.7 g/L Middlebrook 7H9 broth base, 4 g/L glucose, 0.8 g/L NaCl, 5 g/L BSA fraction V, and 0.05% Tyloxapol) and 7H9/DPPC/Chol/BSA (4.7 g/L Middlebrook 7H9, 6 µM DPPC, 62.5 µM cholesterol, 0.8 g/L NaCl, 5 g/L BSA fraction V, and 0.05% Tyloxapol) as previously described^26^.

#### MIC

MIC measurements were performed as previously described^27^. Multiple media were used, with this report highlighting the results from 7H9/DPPC/Cas/Tx (4.7 g/L Middlebrook 7H9, 6 µM DPPC,0.8 g/L NaCl, 0.03% casitone and 0.05% Tyloxapol) and 7H9/Glu/Cas/Tx (4.7 g/L Middlebrook 7H9 broth base, 4 g/L glucose, 0.8 g/L NaCl, 0.03% casitone and 0.05% Tyloxapol). The MICs were determined by visual inspection of the microtitre plates after 1- or 2-weeks growth at 37 °C using an enlarging inverted mirror.

#### Mutant generation and analysis

To raise resistant mutants against **1**, H37Rv cells (10^7^, 10^8^, and 10^9^) were plated on 7H11/OADC plates containing 5 times or 10 times the *in vitro* MIC of **1**. Drug-free plates were used to enumerate bacterial load. The plates were incubated at 37 °C for 4-6 weeks until colonies grew to an appreciable size. To confirm resistance against **1**, colonies were established on drug-free medium and the MIC was determined. Genomic DNA of the mutants was isolated using a CTAB method and sequenced as previously described^28^. Sequencing reads were mapped to the *M. tuberculosis* H37Rv reference genome using BWA^29^. A custom script was used to extract genetic variants (SNPs and indels) by comparative analysis with the parental genome sequence, applying standard filters to exclude sites with low coverage (<10x) or heterogeneous base-calls (<70% purity).

#### Intramacrophage efficacy

J774A.1 mouse macrophages were grown in J774 medium consisting of DMEM supplemented with 4mM L-glutamine, 4.5g/L glucose, 0.5mM sodium pyruvate, 15mM HEPES and 10% FBS. Cells were plated at 1 × 10^4^ cells/mL (1mL/well) in tissue-culture treated 24-well plates and incubated overnight at 37 °C under 5% CO_2_. The macrophages were infected at an MOI of 1 with H37Rv diluted in J774 medium (100 µL/well of 1×10^5^ cells/mL) for 24 hours at 37 °C under 5% CO_2_ after which cells were washed 3 times with warm Dulbecco’s PBS. Cells were fed with J774 medium (1mL/well) containing vehicle control (DMSO) or the compounds. All treatments were done in triplicate wells. Macrophages in five wells were lysed by addition of SDS to 0.1%, homogenization of lysate by repeated pipetting and colony enumeration by plating of appropriate dilutions in 7H9/Glu/BSA on 7H11/OADC plates in duplicate for every dilution. Remaining infected cells were fed every 2 days by removal of spent medium and replenishment with 1mL of the J774 (with added compounds as needed per well). After 7 days of treatment, cells were lysed as before and bacterial counts enumerated by plating of appropriate dilutions in duplicate on 7H11/OADC plates. The 7H11/OADC plates were incubated for 4 weeks at 37 °C before colony enumeration.

#### MIC Comparison in Cell Wall Hypomorphs

MIC measurements for the hypomorph strains were performed as described previously^5^. The cell wall hypomorph strains all grew similarly ±atc, indicating that the different expression levels of the cell wall genes had no impact on cell growth directly.

#### CRISPRi analysis and individual overexpressing strains

CRISPRi profiling was performed as previously described^30^ (Deschner et al. submitted). In brief, triplicate cultures of a CRISPR library were supplemented with atc (0.1 μg/mL), kanamycin (10 μg/mL) and grown for 24h before addition of either DMSO (vehicle control) or **1** dissolved in DMSO. Cultures were then incubated for another 14 days at 37°C in 5% CO_2_. On day 7, atc was replenished. Next, genomic DNA was extracted from bacterial pellets using the Omega Bio-Tek Mag-Bind Universal Pathogen DNA Kit (M4029), with cell lysis performed using a Geno/Grinder and partial automation via a Hamilton Starlet robot. Genomic DNA concentration was quantified using a Thermo Scientific NanoDrop ND-8000 spectrophotometer. The sgRNA-encoding region was subsequently amplified, purified, and pooled for Illumina sequencing as previously described^30^. Construction and characterization of the embCAB overexpression strains was performed as described previously^31^.

#### ADME/PK analysis

Assays to establish HepG2 cytotoxicity, *in vitro* microsomal/hepatocyte metabolic stability, plasma protein binding and *in vivo* pharmacokinetic profiles were all performed as previously described^32^. Likewise, assays for CYP450 inhibition and FaSSIF solubility were performed as described previously^33-34^.

#### Efficacy

Acute efficacy studies involving murine models of TB infection were performed as described^21^.

### Ethical Statements

#### Mouse Pharmacokinetics

All regulated procedures, at the University of Dundee, on living animals was carried out under the authority of a project licence (PP5016780) issued by the Home Office under the Animals (Scientific Procedures) Act 1986, as amended in 2012 (and in compliance with EU Directive EU/2010/63). Licence applications will have been approved by the University’s Ethical Review Committee (ERC) before submission to the Home Office. The ERC has a general remit to develop and oversee policy on all aspects of the use of animals on university premises and is a subcommittee of the University Court, its highest governing body.

#### Acute efficacy studies

All procedures were performed in accordance with protocols (AP32489) approved by the GSK Institutional Animal Care and Use Committee and met or exceeded the standards of the American Association for the Accreditation of Laboratory Animal Care (AAALAC). All animal studies were ethically reviewed and carried out in accordance with European Directive 2010/63/EEC and the GSK Policy on the Care, Welfare and Treatment of Animals.

## Supporting information

ENA04 supplementary information

## Abbreviations

SCX: Strong cation exchange
FA: Formic acid
Pet ether: petroleum ether
br: broad
NPLC: Normal phase liquid chromatography
Cl_int_ mics/heps: Intrinsic microsomal/hepatic clearance
DprE1: Decaprenylphosphoryl-β-d-ribose 2′-epimerase
MmpL3: Mycobacterial membrane protein Large 3
Pks13: Polyketide synthase 13
atc: anhydrotetracycline
DPPC: 1,2-dipalmitoyl-*sn*-glycero-3-phosphocholine
Cas: Casitone
Glu: Glucose
Mol. Sieves: Molecular sieves
EMB: Ethambutol
DPPC: Dipalmitoylphosphatidylcholine
OADC: Oleic Albumin Dextrose Catalase
TB: tuberculosis
ND: not done

## ANCILLARY INFORMATION

### SUPPORTING INFORMATION

The supporting information (PDF) contains

- Table S1: Modifications on R_1_;
- Table S2: Compound **1** resistant mutants
- Figure S1: Compound **1** targets cell wall biosynthesis
- General Chemistry Methods
- Chemistry Experimental Procedures
- Analytical data compounds 1H NMR, 13 C NMR and HRMS **1**-**32**.

Key *in vitro* data CSV file contains

- *In vitro* data, PAINS alert analysis
- Molecular formula strings

## Notes

The authors declare no competing financial interest.

## Acknowledgements

We thank Laura Frame, Fred Simeons, Jennifer Riley, Nicole Mutter, Yoko Shishikura, Karolina Wrobel, Alex Cookson, Kirsty Cookson, Fraser Hughes and Lynsey Swann for technical assistance. In addition, we thank Curtis Engelhart for help with characterization of over expression strains. The work carried out by Dundee was funded by the Gates Foundation (INV-002563). The work at NIH was funded in part by the Gates Foundation (INV001727). The work carried out by WCMC was funded by the Gates Foundation (INV-055896). The work carried out by Texas A&M was funded by the Gates Foundation (INV-002178). The conclusions and opinions expressed in this work are those of the author(s) alone and shall not be attributed to the Foundation. Under the grant conditions of the Foundation, a Creative Commons Attribution 4.0 License has already been assigned to the Author Accepted Manuscript version that might arise from this submission. The NIH work was additionally supported by the Intramural Research Program of NIAID. The contributions of the NIH authors were made as part of their official duties as NIH federal employees, are in compliance with agency policy requirements, and are considered Works of the United States Government. However, the findings and conclusions presented in this paper are those of the author(s) and do not necessarily reflect the views of the NIH or the U.S. Department of Health and Human Services.

## Notes

### Competing Interest Statement

The authors have declared no competing interest.

