## Supplementary material for "Identification and Evaluation of dibasic piperidines as novel cell wall inhibitors against *Mycobacterium tuberculosis*": ENA04 supplementary information

**ORCHID iD** 0000-0001-6218-0092



| Table of Contents | Page |
|---|---|
| Table S1: Modifications on $R_1$ | S2 |
| Table S2: Compound **1** resistant mutants | S2 |
| Figure S1: Compound **1** targets cell wall biosynthesis | S3 |
| General Chemistry Methods | S3 |
| Chemistry Experimental Procedures | S4 |
| 1H and 13 C NMR Analytical data compounds 1-32 | S19 |

**Table S1**: $R_1$Modifications

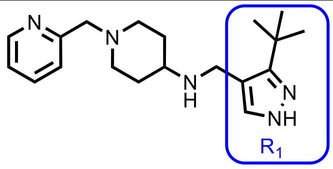


| Compound | $R_1$ | *H37Rv* MIC[A] (µM) | *H37Rv* MIC[B] (µM) | Compound | $R_1$ | *H37Rv* MIC[A] (µM) | *H37Rv* MIC[B] (µM) |
|---|---|---|---|---|---|---|---|
| **1** | 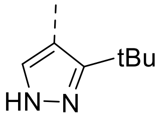 | **3.2** | **3.6** | **7** | 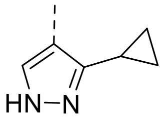 | **37** | **>50** |
| **2** | 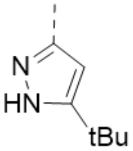 | **>50** | **>50** | **8** | 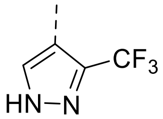 | **>50** | **>50** |
| **3** | 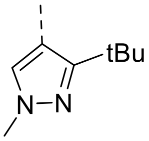 | **>50** | **>50** | **9** | 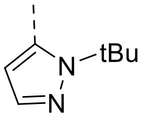 | **>50** | **>50** |
| **4** | 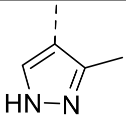 | **>50** | **>50** | **10** | 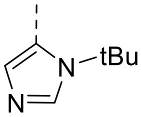 | **>50** | **>50** |
| **5** | 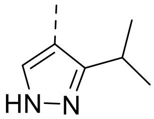 | **25** | **25** | **11** | 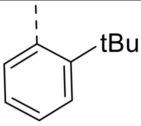 | **>50** | **>50** |
| **6** | 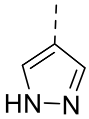 | **>50** | **>50** | | | | |

[a]MIC is the minimum concentration required to inhibit the growth of *M. tuberculosis* (H37Rv) in liquid culture by 90% compared to untreated control (MIC[A] 7H9/DPPC/CAS/Tx media, MIC[B] 7H9/GLU/CAS/Tx media);

**Table S2:** Compound **1** resistant mutants

| Strain | 2 wk MIC (7H9 ADC Tw) | Fold-resistance | Mutations |
|---|---|---|---|
| **WT** | **2.3** | **-** | **-** |
| **A1** | **37** | **16** | **Rv3806c/ubiA:S183P** |
| **B5** | **9.4** | **4** | **embB:D300G** |
| **C5** | **37** | **16** | **embC:T348A** |

MIC required to inhibit the growth of *M. tuberculosis* in liquid culture (7H9/GLU/GLY/Tween). The MIC is shown for wild type H37Rv and each of the three strains resistant to **1**. The fold resistance is also shown along with the SNP identified through whole genome sequencing.

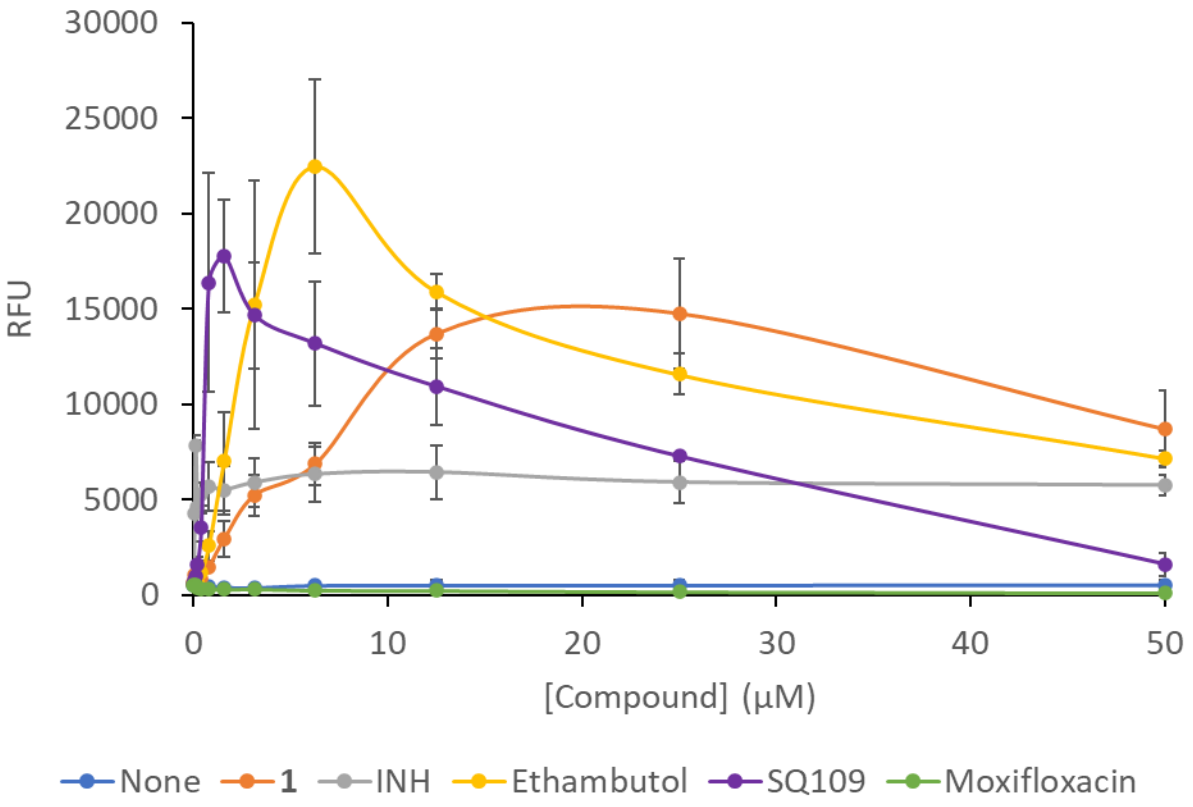


**Figure S1. Compound 1 targets cell wall biosynthesis** H37Rv strain transfected with the Pini-LUX reporter vector was treated with compound as previously reported{Naran, 2016 #265}. Luminescence was monitored at 4 different timepoints, data shown for the 48h timepoint. The relative fluorescence units are shown for the strain treated with DMSO, **1**, INH, Ethambutol, SQ109 (positive controls) and moxifloxacin (negative control). Experiments were performed in duplicate.

**General Chemistry methods**: Normal phase TLCs were carried out on pre-coated silica plates (Kieselgel 60 F254, BDH) with visualization via U.V. light (UV254/365 nm) and/or potassium permanganate solution. Normal phase flash chromatography was performed using Combiflash Companion Rf (commercially available from Teledyne ISCO) and prepacked silica gel columns purchased from Teledyne ISCO. Mass-directed preparative HPLC separations and preparative HPLC chromatographic separations were conducted with either Waters XBridge C18 columns, 100 * 19 mm, 5 µm particle size or Waters X bridge 150 * 25 mm * 5 µm .$^{1}$H NMR spectra were recorded on a Bruker Avance DPX 500 spectrometer ($^{1}$H at 500.1 MHz), or a Bruker Avance DPX 400 ($^{1}$H at 400 MHz) or 500MHz Cryo NMR (Bruker, Avance 3). Chemical shifts (δ) are expressed in ppm recorded using the residual solvent as the internal reference in all cases. Signal splitting patterns are described as singlet (s), doublet (d), triplet (t), quartet (q), multiplet (m), broad (br), or a combination thereof. Coupling constants (J) are quoted to the nearest 0.1 Hz. Due to the tautomeric forms of the NH-pyrazoles within this series of compounds the signals for these cannot always be observed within the $^{13}$C spectra and have therefore not been reported in these cases. High-resolution electrospray measurements were performed on a Bruker Daltonics MicroTOF mass spectrometer or on an Orbitrap Exploris 120 Mass spectrometer. Low resolution electrospray (ES) mass spectra were recorded on an Advion Compact Mass Spectrometer (CMS; model ExpressIon CMS) connected to Dionex Ultimate 3000 UPLC system with diode array detector. HPLC chromatographic separations were conducted using a Waters XBridge C18 column, 2.1 x 50 mm, 3.5 µm particle size or Waters XSelect 2.1 x 30 mm, 2.5 µm particle size. The compounds were eluted with a gradient of 5 to 95 % acetonitrile/water + 0.1 % Ammonia or + 0.1 % formic acid. Unless otherwise stated herein reactions have not been optimized. Solvents and reagents were purchased from commercial suppliers and used without further purification. Dry solvents were purchased in sure sealed bottles stored over molecular sieves. All final compounds showed chemical purity of ≥ 95 % as determined from the UV chromatogram (190−450 nm) obtained by LC-MS analysis.

**Experimental Procedures:**

**General procedures**

**General procedure 1**: NaOAc (2.0 equiv.) was added to a mixture of amine (1.0 equiv. HCl) in EtOH and the reaction stirred at room temperature for 10 min, the desired aldehyde (1.2 equiv.), AcOH (0.13 equiv.) and $NaBH_3CN$ (2.6 equiv.) was added and the reaction stirred for 16 h, quenched with water and concentrated *in vacuo*. The residue was purified by preparative HPLC (10-40 % $CH_3CN$ in water (+1 % $NH_4HCO_3$)) to afford **compounds 1-3.**

**General procedure 2**: To a solution of amine (1.0 equiv.) in DCM and AcOH (0.1 equiv.) was added the desired aldehyde (1.0 equiv.) followed by $NaBH(OAc)_3$ (1.2 equiv.) and the reaction stirred at room temperature for 16 h, diluted with MeOH and transferred to SCX SPE cartridge, washed with MeOH and eluted with 3M $NH_3$ in MeOH. The residue was purified by reverse phase flash column chromatography (5-95 % $CH_3CN$ in water (+0.1 % $NH_4OH$) collecting at 220 nm) to afford **compounds 6-11** and **27-30.**

**General procedure 3**: To a solution of desired aldehyde (1.0 equiv.) and amine (1.0 equiv.) in MeOH was added AcOH (0.1 equiv.) and $NaBH_3CN$ (2 equiv.) and the reaction stirred at room temperature for 12 h, quenched with water, the pH adjusted to ~8 with saturated, aqueous $NaHCO_3$ solution and extracted with EtOAc. The combined organic layers were washed with brine, dried over anhydrous $Na_2SO_4$, filtered and concentrated in vacuo to afford **compounds 4, 5, 12-20** and **31-32.**

**General procedure 4:** To a solution of Boc intermediate (1.0 equiv.) in $CH_3CN$ was added HCl in dioxane (4 M, 14.0 *eq*uiv.). The reaction mixture stirred at room temperature for 0.5 h, concentrated in vacuo to afford intermediates on route to **compounds 12-19.**

**General procedure 5:** To a solution of amine (1.0 equiv.) in THF/EtOH was added the desired aldehyde (1.0 equiv.) 3Å molecular sieves and AcOH (0.1 equiv.). After 1 h $NaBH(OAc)_3$ (2 equiv.) was added and the reaction stirred at room temperature for 16 h. the pH was adjusted to ~7 with aqueous $NaHCO_3$, filtered and the filtrate concentrated in vacuo. The crude material was purified by preparative HPLC conditions to afford **compounds 21-26.**

***tert*-butyl (1-(pyridin-2-ylmethyl)piperidin-4-yl) carbamate (Intermediate 1, R1)**

To a solution of *tert*-butyl piperidin-4-ylcarbamate (31 g, 154 mmol, 1.0 equiv.) and pyridine-2-carbaldehyde (17.4 g, 163 mmol, 1.1 equiv.) and AcOH in MeOH (886 µl, 15.5 mmol, 0.1 equiv., 0.5 M) at room temperature under $N_2$ was added $NaBH(OAc)_3$ (65.6 g, 310 mmol, 2.0 equiv.) in one portion and the reaction stirred for 12 h, concentrated, diluted with water and extracted with DCM. The combined organic extracts were washed with brine, dried over $Na_2SO_4$, filtered and concentrated *in vacuo*. The residue was purified by reverse phase flash column chromatography (5-95 % $CH_3CN$ in water (+0.1 % FA)) to afford *tert*-butyl (1-(pyridin-2-ylmethyl)piperidin-4-yl) carbamate (31.6 g, 107.8 mmol, 70 %) as white solid. **MS** (ES+): *m/z* (%) 292.2 (100) $[M+H]^+$.

**1-(pyridin-2-ylmethyl)piperidin-4-amine hydrochloride ( R1.HCl)**

To a solution of *tert*-butyl (1-(pyridin-2-ylmethyl)piperidin-4-yl)carbamate (31.6 g, 108 mmol, 1.0 equiv.) in $CH_3CN$ (0.36 M) was added 4M HCl in dioxane (11 equiv.) dropwise at 0 °C under $N_2$. The reaction mixture stirred at 10 °C for 1 h, concentrated *in vacuo* and the residue triturated with $CH_3CN$ and filtered to afford 1-(pyridin-2-ylmethyl)piperidin-4-amine hydrochloride (30.5 g, 98.3 mmol, 91 %, 2.8 HCl salt) as an off-white solid. **$^1$H NMR** (400 MHz, DMSO-$d_6$) δ 8.93 - 8.81 (m, 1H), 8.56 - 8.46 (m, 1H), 8.12 (d, *J* = 7.6 Hz, 1H), 8.05 - 7.97 (m, 1H), 4.72 (s, 2H), 3.72 (m, 2H), 3.65

- 3.54 (m, 1H), 3.36 (td, *J* = 2.8, 12.8 Hz, 2H), 2.34 (m, 2H), 2.07 - 1.91 (m, 2H). **MS** (ES+): *m/z* (%) 192.1 (100) $[M+H]^+$.

**1-(pyridin-2-ylmethyl)piperidin-4-amine (R1)**

1-(2-pyridylmethyl)piperidin-4-amine hydrochloride (3.73 g, 16.4 mmol) was dissolved in DCM loaded onto a preconditioned SCX cartridge, washed with DCM, DCM/MeOH and MeOH then eluted with 3M $NH_3$ in MeOH to afford 1-(pyridin-2-ylmethyl)piperidin-4-amine (2.14 g, 11.2 mmol, 65 %) as a light yellow oil. **$^1$H NMR** (500 MHz, MeOD) δ 8.49 (d, *J* = 4.3 Hz, 1H), 7.85 - 7.81 (m ,1H), 7.56 - 7.54 (m, 1H), 7.35 - 7.32 (m, 1H), 3.66 (s, 2H), 2.91 - 2.86 (m, 2H), 2.73 - 2.65 (m, 1H), 2.21 - 2.14 (m, 2H), 1.87 - 1.81 (m, 2H), 1.53 - 1.43 (m, 2H). **MS** (ES+): *m/z* (%) 191.9 (100) $[M+H]^+$.

***tert*-butyl 4-(((3-(tert-butyl)-1*H*-pyrazol-4-yl)methyl)amino)piperidine-1-carboxylate hydroformate(Intermediate 1, R2)**

To a solution of *tert*-butyl 4-aminopiperidine-1-carboxylate (5.5 g, 27.46 mmol, 1.0 equiv.), and 3-(*tert*-butyl)-1*H*-pyrazole-4-carbaldehyde (3.76 g, 24.72 mmol, 0.9 equiv.) in MeOH (0.5 M) was added AcOH (157 µL, 2.75 mmol, 0.1 equiv.) in one portion at rt under $N_2$. The reaction mixture was stirred for 0.5 h, $NaBH_3CN$ (3.45 g, 54.9 mmol, 2.0 equiv.) added and stirred for a further 12 h. The reaction mixture was quenched with water and extracted with EtOAc. The combined organic layers were washed with brine, dried over $Na_2SO_4$, filtered and concentrated *in vacuo* and purified by reverse phase flash column chromatography (5-95 % $CH_3CN$ in water (+0.1 % FA)) to afford *tert*-butyl 4-(((3-(*tert*-butyl)-1*H*-pyrazol-4-yl)methyl)amino)piperidine-1-carboxylate (9.2 g, 24.0 mmol, 98 % , FA salt) as a white solid. **MS** (ES+): *m/z* 337.3 (85) $[M+H]^+$.

***N*-((3-(*tert*-butyl)-1*H*-pyrazol-4-yl)methyl)piperidin-4-amine hydrochloride, (R2.HCl)**

To a solution of tert-butyl 4-(((3-(*tert*-butyl)-1*H*-pyrazol-4-yl)methyl)amino)piperidine-1-carboxylate (9.2 g, 24.0 mmol, 1.0 equiv., FA salt) in $CH_3CN$ (0.3 M) was added HCl in dioxane (4 M, 13.2 equiv.) in one portion at 0 °C under $N_2$. The mixture was stirred at 0 °C for 1 h. The reaction mixture was concentrated *in vacuo* and the residue triturated with $CH_3CN$ to afford *N*-((3-(*tert*-butyl)-1*H*-pyrazol-4-yl)methyl)piperidin-4-amine hydrochloride as a white solid (6.41 g, 17.7 mmol, 65 %, 3.1 HCl salt). **$^1$H NMR (**400 MHz, $D_2O$) δ 8.16 (s, 1H), 4.44 (s, 2H), 3.74 - 3.65 (m, 1H), 3.64 - 3.54 (m, 2H), 3.18 - 3.06 (m, 2H), 2.51 – 5.43 (m, 2H), 2.04 - 1.93 (m, 2H), 1.38 (s, 9H). **MS** (ES+): m/z 237.2 (96) $[M+H]^+$.

***N*-((3-(*tert*-butyl)-1*H*-pyrazol-4-yl)methyl)piperidin-4-amine (R2)**

Synthesized following the same procedure as **R1** from *N*-((3-(*tert*-butyl)-1*H*-pyrazol-4-yl)methyl)piperidin-4-amine (1.23 g, 3.54 mmol, 1.0 equiv. HCl salt) to afford *N*-((3-(*tert*-butyl)-1*H*-pyrazol-4-yl)methyl)piperidin-4-amine (0.83 g, 3.49 mmol, 98 %) as a pale yellow oil. **$^1$H NMR** (500 MHz, MeOD) δ 7.50 (s, 1H), 3.82 (s, 2H), 3.13 - 3.06 (m, 2H), 2.73 - 2.68 (m, 1H), 2.66 - 2.59 (m, 2H), 1.98 - 1.94 (m, 2H), 1.41 - 1.39 (m, 11H). **MS** (ES+): m/z 237 (95) $[M+H]^+$.

***N*-((3-(*tert*-butyl)-1*H*-pyrazol-4-yl)methyl)-1-(pyridin-2-ylmethyl)piperidin-4-amine, (1)**

Synthesized according to **general procedure 1** from 1-(pyridin-2-ylmethyl)piperidin-4-amine hydrochloride (449 mg, 1.97 mmol, 1.0 equiv., HCl salt)and 3-*tert*-butyl-1*H*-pyrazole-4-carbaldehyde (300 mg, 1.97 mmol, 1 equiv.) in DCM (0.3 M) using NaOAc (1.5 equiv.) and $NaBH_3CN$ (2.0 equiv.) to afford *N*-((3-(*tert*-butyl)-1*H*-pyrazol-4-yl)methyl)-1-(pyridin-2-ylmethyl)piperidin-4-amine (230 mg, 0.69 mmol, 35 %) as a white gum. **$^1$H NMR** (500 MHz, MeOD) δ 8.54 – 8.51 (m, 1H), 7.88 – 7.84 (m, 1H), 7.7 (br s, 1H), 7.55 (d, *J* = 7.8 Hz, 1H), 7.38 – 7.34 (m, 1H), 4.29 (s, 2H), 3.75 (s, 2H), 3.30 – 3.22 (m, 1H), 3.10 – 3.04 (m, 2H), 2.34 – 2.26 (m, 2H), 2.22 – 2.16 (m, 2H), 1.83 – 1.73 (m, 2H), 1.43 (s, 9H). **$^{13}$C NMR** (126 MHz, MeOD) δ 157.4, 148.4, 140.6,

137.4, 123.8, 122.7, 107.7, 62.6, 55.4, 51.3, 39.7, 32.0, 29.3, 28.3. **HRMS** (ES+): calcd. for $C_{19}H_{29}N_5$ [M+H]$^+$ 328.2496, found 328.2505 (2.7 ppm).

**(5-(*tert*-butyl)-1*H*-pyrazol-3-yl)methanol (2-S1)**

To a solution of ethyl 5-(*tert*-butyl)-1*H*-pyrazole-3-carboxylate (500 mg, 2.6 mmol, 1.0 equiv.) in THF (0.3 M) was added $LiAlH_4$ (290 mg, 7.64 mmol, 2.9 equiv.) at 0 °C and the mixture was stirred at room temperature for 2.5 h. The reaction quenched with 15 % aqueous NaOH solution slowly at 0 °C and the solution extracted with EtOAc. The combined organic extracts were dried over $Na_2SO_4$, filtered and concentrated *in vacuo* to afford (5-(*tert*-butyl)-1*H*-pyrazol-3-yl)methanol (0.4 g, crude) as a white solid.

**5-(*tert*-butyl)-1*H*-pyrazole-3-carbaldehyde (2-S2)**

A solution of (5-(*tert*-butyl)-1*H*-pyrazol-3-yl)methanol (400 mg, 2.6 mmol, 1.0 equiv.) and $MnO_2$ (2.26 g, 25.9 mmol, 10 equiv.) in THF (0.43 M) was stirred at room temperature for 1 h. The reaction mixture was filtered and concentrated *in vacuo* to afford 5-(*tert*-butyl)-1*H*-pyrazole-3-carbaldehyde (378 mg, crude) as a yellow oil. **MS** (ES+): *m/z* (%) 153.2 (100) [M+H]$^+$.

***N*-((5-(*tert*-butyl)-1*H*-pyrazol-3-yl)methyl)-1-(pyridin-2-ylmethyl)piperidin-4-amine, (2)**

Synthesized according to **general procedure 1** from 1-(pyridin-2-ylmethyl)piperidin-4-amine hydrochloride (570 mg, 2.50 mmol, 1.0 equiv., HCl salt) and 5-(*tert*-butyl)-1*H*-pyrazole-3-carbaldehyde (343 mg, 2.25 mmol, 0.9 equiv.) in EtOH (0.5 M) to afford *N*-((5-(*tert*-butyl)-1*H*-pyrazol-3-yl)methyl)-1-(pyridin-2-ylmethyl)piperidin-4-amine (141 mg, 0.41 mmol, 18 %) as a colorless gum. **$^1$H NMR** (500 MHz, MeOD) δ 8.48 (d, *J* = 4.2 Hz, 1H), 7.85 - 7.81 (m, 1H), 7.57 - 7.54 (m, 1H), 7.34 - 7.30 (m, 1H), 6.12 (s, 1H), 3.78 (s, 2H), 3.66 - 3.65 (m, 2H), 2.91 (d, *J* = 12.1 Hz, 2H), 2.59 - 2.50 (m, 1H), 2.17 - 2.10 (m, 2H), 1.96 - 1.90 (m, 2H), 1.54 - 1.44 (m, 2H), 1.32 (s, 9H). **$^{13}$C NMR** (126 MHz, MeOD) δ 158.1, 148.1, 137.3, 123.8, 122.4, 99.9, 63.4, 53.8, 52.2, 31.2, 29.3. **HRMS** (ES+): calcd. for $C_{19}H_{29}N_5$ [M+H]$^+$ 328.2496, found 328.2496 (0.0 ppm).

***3.1.* Preparation of 3-(*tert*-butyl)-1-methyl-1*H*-pyrazole-4-carbaldehyde (3-S1)**

A mixture of 3-(*tert*-butyl)-1*H*-pyrazole-4-carbaldehyde (300 mg, 1.97 mmol, 1.0 equiv.), $CH_3I$ (560 mg, 3.94 mmol, 245 µL, 2.0 equiv.) and $K_2CO_3$ (1.09 g, 7.88 mmol, 4.0 equiv.) in DMF (0.5 M) was stirred at 85 °C for 1.5 h, quenched with saturated, aqueous$NH_4Cl$ solution, extracted with EtOAc and the combined organic extracts were dried over $Na_2SO_4$, filtered and concentrated *in vacuo* and the residue was purified by reverse phase flash column chromatography (5-95 % $CH_3CN$ in water (+0.1 % FA)) to afford 3-(*tert*-butyl)-1-methyl-1*H*-pyrazole-4-carbaldehyde (255 mg, 1.46 mmol, 74 %) as yellow oil. **$^1$H NMR** (400 MHz, $CDCl_3$) δ 10.00 (s, 1H), 7.87 (s, 1H), 3.87 (s, 3H), 1.41 (s, 9H). **MS** (ES+): *m/z* (%) 167.2 (100) [M+H]$^+$.

***N*-((3-(*tert*-butyl)-1-methyl-1*H*-pyrazol-4-yl)methyl)-1-(pyridin-2-ylmethyl)piperidin-4-amine (3)**

Synthesized according to **general procedure 1** from 1-(pyridin-2-ylmethyl)piperidin-4-amine hydrochloride (300 mg, 1.01 mmol, 1.0 equiv., HCl salt) and 3-(*tert*-butyl)-1-methyl-1*H*-pyrazole-4-carbaldehyde (197 mg, 1.19 mmol, 1.2 equiv.) in EtOH (0.3 M) to afford *N*-((3-(*tert*-butyl)-1-methyl-1*H*-pyrazol-4-yl)methyl)-1-(pyridin-2-ylmethyl)piperidin-4-amine (88 mg, 0.24 mmol, 19 %) as a yellow gum. **$^1$H NMR** (500 MHz, MeOD) δ 8.50 - 8.47 (m, 1H), 7.86 - 7.82 (m, 1H), 7.56 (d, *J* = 7.9 Hz, 1H), 7.47 (s, 1H), 7.35 - 7.31 (m, 1H), 3.79 - 3.78 (m, 5H), 3.67 - 3.66 (m, 2H), 2.92 (d, *J* = 11.9 Hz, 2H), 2.64 - 2.55 (m, 1H), 2.21 - 2.15 (m, 2H), 1.97 - 1.91 (m, 2H), 1.56 - 1.46 (m, 2H), 1.35 - 1.34 (m, 9H). **$^{13}$C NMR** (126 MHz, MeOD) δ 158.1, 157.1, 148.2, 137.3, 131.6, 124.8, 122.5, 115.9, 63.5, 54.4, 52.2, 41.1, 37.6, 32.6, 31.5, 29.3. **HRMS** (ES+): calcd. for $C_{20}H_{31}N_5$ [M+H]$^+$ 342.2652, found 342.2653 (0.3 ppm). MS (ES+): *m/z* (%) 342.4 (100) [M+H]$^+$.

***N*-((3-methyl-1*H*-pyrazol-4-yl)methyl)-1-(pyridin-2-ylmethyl)piperidin-4-amine (4)**

Synthesized according to **general procedure 3** from 1-(pyridin-2-ylmethyl)piperidin-4-amine hydrochloride (300 mg, 1.32 mmol, 1.0 equiv., HCl salt) and 3-methyl-1*H*-pyrazole-4-carbaldehyde (145 mg, 1.32 mmol, 1.0 equiv.) in EtOH (0.2 M) using AcOH (1.5 equiv.) in addition to NaOAc (0.1 equiv.).The reaction mixture was extracted with DCM and the combined organic extracts were concentrated *in vacuo* and purified by preparative TLC ($SiO_2$, 10:1 DCM/MeOH, $R_f$ = 0.15) and preparative HPLC (6-36 % $CH_3CN$ in water (+1 % $NH_4HCO_3$)) to afford *N*-((3-methyl-1*H*-pyrazol-4-yl)methyl)-1-(pyridin-2-ylmethyl)piperidin-4-amine (16 mg, 0.05 mmol, 4 %) as a colorless gum. **$^1$H NMR** (500 MHz, MeOD) δ 8.50 – 8.47 (m, 1H), 7.85 – 7.80 (m, 1H), 7.56 (d, *J* = 7.4 Hz, 1H), 7.51 (s, 1H), 7.34 – 7.30 (m, 1H), 3.66 (s, 2H), 3.65 (s, 2H), 2.95 – 2.89 (m, 2H), 2.59 – 2.51 (m, 1H), 2.27 (s, 3H), 2.19 – 2.10 (m, 2H), 1.98 – 1.90 (m, 2H), 1.55 – 1.44 (m, 2H). **$^{13}$C NMR** (126 MHz, MeOD) δ 158.0, 148.1, 137.2, 123.8, 122.5, 115.5, 63.5, 53.9, 52.3, 38.8, 31.3, 8.7. **HRMS** (ES+): calcd. for $C_{16}H_{24}N_5$ $[M+H]^+$ 286.2026, found 286.2024 (0.6 ppm).

**(3-isopropyl-1*H*-pyrazol-4-yl)methanol (5-S1)**

To a solution of ethyl 3-isopropyl-1*H*-pyrazole-4-carboxylate (500 mg, 2.74 mmol, 1.0 equiv.) in THF (0.14 M) was added $LiAlH_4$ (416 mg, 10.9 mmol, 4.0 equiv.) at 0 °C. The reaction mixture warmed to room temperature then heated at 60 °C.After 2 h, the mixture was cooled to 0 °C, brine was added dropwise and the reaction mixture filtered. The filtrate was lyophilized to give a residue which was purified by normal phase flash chromatography eluting with 2:1 Pet ether:EtOAc to 10:1 DCM:MeOH to afford (3-isopropyl-1*H*-pyrazol-4-yl)methanol (260 mg, 1.84 mmol, 67 %) as a white solid. **$^1$H NMR** (400 MHz, MeOD) δ 7.66 - 7.20 (m, 1H), 4.50 (br s, 2H), 3.25 - 3.05 (m, 1H), 1.30 (br d, *J* = 7.2 Hz, 6H). **MS** (ES+): *m/z* (%) 141.2 $[M+H]^+$.

**3-isopropyl-1*H*-pyrazole-4-carbaldehyde (5-S2)**

To a solution of (3-isopropyl-1*H*-pyrazol-4-yl)methanol (200 mg, 1.43 mmol, 1.0 equiv.) in THF (0.04 M) was added Dess-Martin periodinane (907 mg, 2.14 mmol, 1.5 equiv.). The reaction mixture stirred at 25 °C for 1 h, filtered and concentrated *in vacuo,* saturated, aqeous $NaHCO_3$ solution and saturated, aqueous $Na_2SO_3$ solution were added to the residue and the mixture was extracted with 10:1 DCM:MeOH. The combined organic extracts were washed with brine, dried over anhydrous $Na_2SO_4$, filtered and concentrated *in vacuo* to afford 3-isopropyl-1*H*-pyrazole-4-carbaldehyde (120 mg, 0.87 mmol, 61 %) as a yellow oil. **$^1$H NMR** (400 MHz, MeOD) δ 9.89 (s, 1H), 8.02 (br s, 1H), 3.54 (br s, 1H), 1.33 (br d, *J* = 6.8 Hz, 6H). **MS** (ES+): *m/z* (%) 139.2 (98) $[M+H]^+$.

***N*-((3-isopropyl-1*H*-pyrazol-4-yl)methyl)-1-(pyridin-2-ylmethyl)piperidin-4-amine (5)**

Synthesized according to **general procedure 3** from 1-(pyridin-2-ylmethyl)piperidin-4-amine hydrochloride (217 mg, 0.96 mmol, 1.2 equiv., HCl salt) and 3-isopropyl-1*H*-pyrazole-4-carbaldehyde (110 mg, 0.79 mmol, 1.0 *eq*uiv.) in MeOH (0.16 M) using AcOH (1.0 equiv.) in addition to NaOAc (2.0 equiv.).The product extracted with 10:1 DCM:MeOH, the organic extracts were concentrated *in vacuo* and the residue purified by preparative HPLC (6-36 % $CH_3CN$ in water (+1 % $NH_4HCO_3$) using Phenomenex Gemini-NX C18 75 * 30 mm, 3 µm column) to afford *N*-((3-isopropyl-1*H*-pyrazol-4-yl)methyl)-1-(pyridin-2-ylmethyl)piperidin-4-amine (34.3 mg, 0.11 mmol, 13 %) as a yellow gum. **$^1$H NMR** (500 MHz, MeOD) δ 8.50 – 8.47 (m, 1H), 7.86 – 7.81 (m, 1H), 7.56 (d, *J* = 7.8 Hz, 1H), 7.51 (s, 1H), 7.35 – 7.30 (m, 1H), 3.68 (s, 2H), 3.66 (s, 2H), 3.15 – 3.06 (m, 1H), 2.95 – 2.88 (m, 2H), 2.60 – 2.52 (m, 1H), 2.19 – 2.11 (m, 2H), 1.97 – 1.90 (m, 2H), 1.56 – 1.45 (m, 2H), 1.30 (d, *J* = 7Hz, 6H). **$^{13}$C NMR** (126 MHz, MeOD) δ 158.0, 148.1, 137.3, 123.8, 122.5, 63.4, 53.9, 52.3, 38.9, 31.3, 21.4. **HRMS** (ES+): calcd. for $C_{18}H_{28}N_5$ $[M+H]^+$ 314.2339, found 314.2347 (2.5 ppm).

***N*-((3-(tert-butyl)-1*H*-pyrazol-4-yl)methyl)-1-(pyridin-2-ylmethyl)piperidin-4-amine (6)**

Synthesized according to **general procedure 2** from 1-(pyridin-2-ylmethyl)piperidin-4-amine (100 mg, 0.52 mmol, 1.0 equiv.) and 1*H*-pyrazole-4-carbaldehyde (50 mg, 0.52 mmol, 1.0 equiv.) in DCM (0.5 M) to afford *N*-((3-(tert-butyl)-1*H*-pyrazol-4-yl)methyl)-1-(pyridin-2-ylmethyl)piperidin-4-amine (60 mg, 0.21 mmol, 40 %) as a clear oil. **$^1$H NMR** (500 MHz, MeOD) δ 8.50 – 8.47 (m, 1H), 7.85 – 7.80 (m, 1H), 7.62 – 7.52 (m, 3H), 7.35 – 7.30 (m, 1H), 3.73 (s, 2H), 3.65 (s, 2H), 2.94 – 2.88 (m, 2H), 2.58 – 2.50 (m, 1H), 2.17 – 2.10 (m, 2H), 1.96 – 1.89 (m, 2H), 1.54 – 1.44 (m, 2H). **$^{13}$C NMR** (126 MHz, MeOD) δ 158.0, 148.1, 137.3, 123.8, 122.5, 118.4, 63.4, 53.6, 52.3, 39.4, 31.2. **HRMS** (ES+): calcd. for $C_{15}H_{22}N_5$ [M+H]$^+$ 272.1870, found 272.1870 (0.2 ppm).

***N*-((3-cyclopropyl-1*H*-pyrazol-4-yl)methyl)-1-(pyridin-2-ylmethyl)piperidin-4-amine (7)**

Synthesized according to **general procedure 2** from 1-(pyridin-2-ylmethyl)piperidin-4-amine (95 mg, 0.49 mmol, 1.0 equiv.) and 3-cyclopropyl-1*H*-pyrazole-4-carbaldehyde (68 mg, 0.49 mmol, 1.0 equiv.) in DCM (0.25 M) to afford *N*-((3-cyclopropyl-1*H*-pyrazol-4-yl)methyl)-1-(pyridin-2-ylmethyl)piperidin-4-amine (90 mg, 0.27 mmol, 55 %) as a glassy solid. **$^1$H NMR** (500 MHz, MeOD) δ 8.50 – 8.47 (m, 1H), 7.86 – 7.81 (m, 1H), 7.56 (d, *J* = 7.8 Hz, 1H), 7.50 (s, 1H), 7.35 – 7.30 (m, 1H), 3.75 (s, 2H), 3.66 (s, 2H), 2.95 – 2.88 (m, 2H), 2.60 – 2.52 (m, 1H), 2.19 – 2.11 (m, 2H), 1.98 – 1.92 (m, 2H), 1.92 – 1.86 (m, 1H), 1.56 – 1.45 (m, 2H), 0.97 – 0.90 (m, 2H), 0.82 – 0.76 (m, 2H). **$^{13}$C NMR** (126 MHz, MeOD) δ 158.0, 148.1, 137.2, 123.8, 122.5, 116.4, 63.5, 52.3, 38.8, 31.3, 5.7. **HRMS** (ES+): calcd. for $C_{18}H_{26}N_5$ [M+H]$^+$ 312.2183, found 312.2187 (1.5 ppm).

**(3-(trifluoromethyl)-1*H*-pyrazol-4-yl)methanol (8-S1)**

To a solution of ethyl 5-(trifluoromethyl)-1*H*-pyrazole-4-carboxylate (300 mg, 1.44 mmol, 1.0 equiv.) in THF (0.18 M) cooled to 0 °C under $N_2$ was added 1M $LiAlH_4$ (3.6 mL, 3.60 mmol, 2.5 equiv.) and the reaction stirred for 2 h. The mixture was quenched by the dropwise addition of water while cooling in an ice bath and the mixture was extracted with EtOAc and the solution filtered to remove precipitate. The organic phase were separated, dried over $Na_2SO_4$, filtered and concentrated *in vacuo* to yield (3-(trifluoromethyl)-1*H*-pyrazol-4-yl)methanol (173 mg, 0.83 mmol, 58 %). **$^1$H NMR** (500MHz, MeOD) δ 7.77 (s, 1H), 4.63 (s, 2H).

**3-(trifluoromethyl)-1*H*-pyrazole-4-carbaldehyde (-S2)**

To a suspension of [5-(trifluoromethyl)-1*H*-pyrazol-4-yl]methanol (173 mg, 0.83 mmol, 1.0 equiv.) in DCM (0.17 M) cooled to 0 °C was added Dess-Martin periodinane (441 mg, 1.04 mmol, 1.25 equiv.) and the reaction slowly warmed to room temperature and stirred for 16 h, diluted with DCM, filtered and concentrated *in vacuo* and purified by normal phase flash chromatography (0-100 % EtOAc in heptane) to afford 3-(trifluoromethyl)-1*H*-pyrazole-4-carbaldehyde (54 mg, 0.32 mmol, 39 %). **$^1$H NMR** (500 MHz, MeOD) δ 9.95 (s, 1H), 8.43 (s, 1H).

**1-(pyridin-2-ylmethyl)-*N*-((3-(trifluoromethyl)-1*H*-pyrazol-4-yl)methyl)piperidin-4-amine (8)**

Synthesized according to **general procedure 2** from 1-(pyridin-2-ylmethyl)piperidin-4-amine (63 mg, 0.33 mmol, 1.0 equiv.) and 3-(trifluoromethyl)-1*H*-pyrazole-4-carbaldehyde (54 mg, 0.33 mmol, 1.0 equiv.) in DCM (0.17 M) to afford 1-(pyridin-2-ylmethyl)-*N*-((3-(trifluoromethyl)-1*H*-pyrazol-4-yl)methyl)piperidin-4-amine (24 mg, 0.067 mmol, 20 %) as red sticky gum. **$^1$H NMR** (500 MHz, MeOD) δ 8.50 – 8.47 (m, 1H), 7.86 – 7.82 (m, 1H), 7.81 (s, 1H), 7.56 (d, *J* = 7.8 Hz, 1H), 7.35 – 7.30 (m, 1H), 3.81 (s, 2H), 3.66 (s, 2H), 2.95 – 2.88 (m, 2H), 2.59 – 2.51 (m, 1H), 2.19 – 2.11 (m, 2H), 1.96 – 1.89 (m, 2H), 1.55 – 1.44 (m, 2H). **$^{13}$C NMR** (126 MHz, MeOD) δ 157.9, 148.1, 137.2, 129.9, 123.8, 123.3, 122.5, 121.1, 117.7, 63.4, 53.9, 52.2, 38.5, 31.3. **HRMS** (ES+): calcd. for $C_{16}H_{21}F_3N_5$ [M+H]$^+$ 340.1744, found 340.1752 (2.5 ppm).

***N*-((1-(*tert*-butyl)-1*H*-pyrazol-5-yl)methyl)-1-(pyridin-2-ylmethyl)piperidin-4-amine (9)**

Synthesized according to **general procedure 2** from 1-(pyridin-2-ylmethyl)piperidin-4-amine (40 mg, 0.21 mmol, 1.0 equiv.) and 1-*tert*-butyl-1*H*-pyrazole-5-carbaldehyde (32 mg, 0.21 mmol, 1.0 equiv.) in DCM (0.2 M) to afford *N*-((1-(*tert*-butyl)-1*H*-pyrazol-5-yl)methyl)-1-(pyridin-2-ylmethyl)piperidin-4-amine (25 mg, 0.07 mmol, 35 %) as a colorless gum. **$^1$H NMR** (500 MHz, MeOD) δ 8.49 - 8.48 (m, 1H), 7.86 - 7.81 (m, 1H), 7.58 - 7.55 (m, 1H), 7.35 - 7.31 (m, 2H), 6.34 (d, *J* = 1.7 Hz, 1H), 4.03 (s, 2H), 3.67 (s, 2H), 2.95 - 2.90 (m, 2H), 2.65 - 2.58 (m, 1H), 2.21 - 2.15 (m, 2H), 1.98 - 1.92 (m, 2H), 1.65 (s, 9H), 1.57 - 1.48 (m, 2H). **$^{13}$C NMR** (126 MHz, MeOD) δ 157.9, 148.1, 141.7, 137.3, 136.2, 123.8, 122.5, 106.8, 63.5, 60.1, 54.4, 52.3, 43.0, 31.5, 29.2. **HRMS** (ES+): calcd. for $C_{19}H_{29}N_5$ $[M+H]^+$ 328.2496, found 328.2489 (2.1 ppm).

***N*-((1-(*tert*-butyl)-1*H*-imidazol-5-yl)methyl)-1-(pyridin-2-ylmethyl)piperidin-4-amine (10)**

Synthesized according to **general procedure 2** from 1-(pyridin-2-ylmethyl)piperidin-4-amine (40 mg, 0.21 mmol, 1.0 equiv.)and 1-(*tert*-butyl)-1*H*-imidazole-5-carbaldehyde (32 mg, 0.21 mmol, 1.0 equiv.) in DCM (0.2 M) to afford *N*-((1-(*tert*-butyl)-1*H*-imidazol-5-yl)methyl)-1-(pyridin-2-ylmethyl)piperidin-4-amine (28 mg, 0.08 mmol, 38 %) as a gum. **$^1$H NMR** (500 MHz, MeOD) δ 8.49 - 8.47 (m, 1H), 7.85 - 7.81 (m, 1H), 7.73 (d, *J* = 1.4 Hz, 1H), 7.58 - 7.55 (m, 1H), 7.34 - 7.31 (m, 1H), 7.22 - 7.21 (m, 1H), 3.72 (s, 2H), 3.66 - 3.65 (m, 2H), 2.94 - 2.89 (m, 2H), 2.59 - 2.51 (m, 1H), 2.18 - 2.11 (m, 2H), 1.95 - 1.90 (m, 2H), 1.59 (s, 9H), 1.54 - 1.46 (m, 2H). **$^{13}$C NMR** (126 MHz, MeOD) δ 158.1, 148.1, 138.9, 137.3, 134.0, 123.8, 122.5, 114.5, 63.5, 55.0, 53.8, 52.2, 42.9, 31.3, 29.3. **HRMS** (ES+): calcd. for $C_{19}H_{29}N_5$ $[M+H]^+$ 328.2496, found 328.2508 (3.8 ppm).

***N*-[(2-*tert*-butylphenyl)methyl]-1-(2-pyridylmethyl)piperidin-4-amine (11)**

Synthesized according to **general procedure 2** from 1-(pyridin-2-ylmethyl)piperidin-4-amine (40 mg, 0.21 mmol, 1.0 equiv.) and 2-*tert*-butylbenzaldehyde (34 mg, 0.21 mmol, 1.0 equiv.) in DCM (0.2 M) to afford *N*-[(2-*tert*-butylphenyl)methyl]-1-(2-pyridylmethyl)piperidin-4-amine (24 mg, 0.068 mmol, 32 %) as a gum. **$^1$H NMR** (500 MHz, MeOD) δ 8.50 - 8.48 (m, 1H), 7.86 - 7.82 (m, 1H), 7.57 (d, *J* = 7.8 Hz, 1H), 7.44 - 7.38 (m, 2H), 7.35 - 7.31 (m, 1H), 7.21 - 7.15 (m, 2H), 4.01 (s, 2H), 3.67 (s, 2H), 2.96 - 2.91 (m, 2H), 2.69 - 2.62 (m, 1H), 2.23 - 2.16 (m, 2H), 2.00 - 1.95 (m, 2H), 1.60 - 1.51 (m, 2H), 1.44 - 1.44 (m, 9H). **$^{13}$C NMR** (126 MHz, MeOD) δ 158.0, 148.1, 147.4, 138.2, 137.3, 130.9, 126.6, 125.8, 125.7, 123.8, 122.5, 63.5, 55.3, 52.3, 48.7, 35.2, 31.6, 31.0. **HRMS** (ES+): calcd. for $C_{22}H_{31}N_3$ $[M+H]^+$ 338.2591, found 338.2590 (0.2 ppm).

***tert*-butyl (3-((pyridin-2-ylmethyl)amino)propyl)carbamate (Intermediate 1, 12)**

Synthesized according to **general procedure 3** from *tert*-butyl (3-aminopropyl)carbamate (976 mg, 5.60 mmol, 978 µL, 1.2 equiv.) and pyridine-2-carbaldehyde (500 mg, 4.67 mmol, 1.0 equiv.) in EtOH (0.47 M). The reaction mixture was quenched with water, extracted with DCM and the combined organic extracts were concentrated *in vacuo* and purified by flash chromatography eluting with 5:1 pet ether:EtOAc to 3:1 EtOAc:MeOH followed by reverse phase flash column chromatography (5-95 % $CH_3CN$ in water (+0.1 % FA)) to afford *tert*-butyl (3-((pyridin-2-ylmethyl)amino)propyl)carbamate (450 mg, 1.7 mmol, 36 %) as a yellow oil. **$^1$H NMR** (400 MHz, $CDCl_3$) δ 8.56 (d, *J* = 4.4 Hz, 1H), 7.70 - 7.62 (m, 1H), 7.33 (d, *J* = 8.0 Hz, 1H), 7.21 - 7.15 (m, 1H), 5.21 (br s, 1H), 3.95 (s, 2H), 3.30-3.19 (m, 2H), 2.76 (t, *J* = 6.4 Hz, 2H), 1.81 - 1.68 (m, 2H), 1.44 (s, 9H). MS m/z 266.3 $[M+H]^+$. **MS** (ES+): *m/z* (%) 266.3 (99) $[M+H]^+$.

***N*$^1$-(pyridin-2-ylmethyl)propane-1,3-diamine hydrochloride (Intermediate 2, 12)**

Synthesized according to **general procedure 5** from *tert*-butyl (3-((pyridin-2-ylmethyl)amino)propyl)carbamate (400 mg, 1.51 mmol, 1.0 equiv.) in $CH_3CN$ (0.76 M) to afford *N*$^1$-(pyridin-2-ylmethyl)propane-1,3-diamine (300 mg, crude, HCl salt) as a yellow solid that was used without further purification.

***N*$^1$-((3-(*tert*-butyl)-1*H*-pyrazol-4-yl)methyl)-*N*$^3$-(pyridin-2-ylmethyl)propane-1,3-diamine (12)**

Synthesized according to **general procedure 3** from *N*[1]-(pyridin-2-ylmethyl)propane-1,3-diamine hydrochloride (300 mg, 1.49 mmol, 1.3 equiv., HCl salt) and 3-*tert*-butyl-1*H*-pyrazole-4-carbaldehyde (174 mg, 1.14 mmol, 1.0 equiv.) using EtOH (0.2 M) with the addition of NaOAc (1.5 equiv.). The reaction mixture was quenched with water, concentrated *in vacuo* and purified by flash chromatography (20-35% MeOH in EtOAc)and preparative HPLC (10-40 % $CH_3CN$ in water (+1 % $NH_4HCO_3$)) to afford product which was further purified by preparative HPLC (0-18 % $CH_3CN$ in water (+0.05 % HCl) using 3_Phenomenex Luna C18 75 * 30 mm * 3 µm column) to afford *N*[1]-((3-(*tert*-butyl)-1*H*-pyrazol-4-yl)methyl)-*N*[3]-(pyridin-2-ylmethyl)propane-1,3-diamine (33 mg, 0.10 mmol, 9 %) as a white solid. **$^1$H NMR** (500 MHz, MeOD) δ 8.90 - 8.87 (m, 1H), 8.54 (s, 1H), 8.46 - 8.42 (m, 1H), 8.11 (d, *J* = 7.9 Hz, 1H), 7.93 - 7.89 (m, 1H), 4.68 (s, 2H), 4.49 (s, 2H), 3.47 - 3.39 (m, 4H), 2.45 - 2.36 (m, 2H), 1.53 - 1.52 (m, 9H). **$^{13}$C NMR** (126 MHz, MeOD) δ 154.6, 147.6, 145.4, 143.5, 136.1, 126.5, 126.2, 110.4, 48.4, 45.0, 44.9, 41.5, 32.7, 28.6, 22.7. **HRMS** (ES+): calcd. for $C_{17}H_{27}N_5$ $[M+H]^+$ 302.2339, found 302.2338 (0.3 ppm).

***tert*-butyl (2-((pyridin-2-ylmethyl)amino)ethyl)carbamate (Intermediate 1, 13)**

Synthesized according to **general procedure 3** from *tert*-butyl 2-aminoethyl)carbamate (897 mg, 5.60 mmol, 880 µL, 1.2 equiv.) and picolinaldehyde (500 mg, 4.67 mmol, 1.0 equiv.) in EtOH (0.47 M) to afford *tert*-butyl (2-((pyridin-2-ylmethyl)amino)ethyl)carbamate (450 mg, 1.76 mmol, 38 %) as a yellow oil.. **$^1$H NMR** (400 MHz, $CDCl_3$) δ 8.57 (d, *J* = 4.8 Hz, 1H), 7.70 - 7.63 (m, 1H), 7.30 (d, *J* = 7.6 Hz, 1H), 7.22 - 7.16 (m, 1H), 5.18 (br s, 1H), 3.96 (s, 2H), 3.35 - 3.25 (m, 2H), 2.85 - 2.82 (m, 2H), 1.45 (s, 9H). **MS** (ES+): *m/z* (%) 252.3 (100) $[M+H]^+$.

***N*[1]-(pyridin-2-ylmethyl)ethane-1,2-diamine hydrochloride (Intermediate 2, compound 13)**

Synthesized according to **general procedure 5** from *tert*-butyl (2-((pyridin-2-ylmethyl)amino)ethyl)carbamate (450 mg, 1.76 mmol, 1.0 equiv.) in $CH_3CN$ (0.76 M) to afford *N*[1]-(pyridin-2-ylmethyl)ethane-1,2-diamine (290 mg, crude, HCl salt) as a yellow solid that was used without further purification.

***N*[1]-((3-(*tert*-butyl)-1*H*-pyrazol-4-yl)methyl)-*N*[2]-(pyridin-2-ylmethyl)ethane-1,2-diamine (13)**

Synthesized according to **general procedure 3** from *N*[1]-(pyridin-2-ylmethyl)ethane-1,2-diamine hydrochloride (290 mg, 1.55 mmol, 1.3 equiv., HCl salt) and 3-*tert*-butyl-1*H*-pyrazole-4-carbaldehyde (182 mg, 1.19 mmol, 1.0 equiv.) using EtOH (0.2 M) with the addition of NaOAc (1.5 equiv.). The reaction mixture was quenched with water, concentrated *in vacuo* and the residue was purified by preparative HPLC (10-40 % $CH_3CN$ in water (+1 % $NH_4HCO_3$) to afford product which was further purified by preparative HPLC (0-15 % $CH_3CN$ in water (+0.05 % HCl) using Phenomenex Synergi C18 150 * 25 mm * 10 µm column) followed by additional preparative HPLC (0-5 % $CH_3CN$ in water (+0.05 % HCl) using 3_Phenomenex Luna C18 75 * 30 mm, 3 µm column) to afford *N*[1]-((3-(*tert*-butyl)-1*H*-pyrazol-4-yl)methyl)-*N*[2]-(pyridin-2-ylmethyl)ethane-1,2-diamine (25 mg, 0.08 mmol, 7 %) as a yellow solid. **$^1$H NMR** (500 MHz, MeOD) δ 8.87 (d, *J* = 4.1 Hz, 1H), 8.56 (s, 1H), 8.42 (t, *J* = 7.2 Hz, 1H), 8.13 (d, *J* = 7.5 Hz, 1H), 7.93 - 7.87 (m, 1H), 4.75 (s, 2H), 4.57 (s, 2H), 4.56 (s, 2H), 3.80 (s, 4H), 1.53 (s, 9H). **$^{13}$C NMR** (126 MHz, MeOD) δ 154.4, 147.6, 145.4, 143.5, 136.2, 126.4, 126.2, 110.1, 48.6, 43.8, 41.8, 32.7, 28.7. **HRMS** (ES+): calcd. for $C_{16}H_{25}N_5$ $[M+H]^+$ 288.2183, found 288.2180 (0.8 ppm).

***tert*-butyl (*R*)-3-((((3-(*tert*-butyl)-1*H*-pyrazol-4-yl)methyl)amino)methyl) pyrrolidine-1-carboxylate (Intermediate 1, 14)**

Synthesized according to **general procedure 3** from *tert*-butyl (3*R*)-3-(aminomethyl)pyrrolidine-1-carboxylate (500 mg, 2.50 mmol, 1.0 equiv.) and 3-*tert*-butyl-1*H*-pyrazole-4-carbaldehyde (303 mg, 2.00 mmol, 0.8 equiv.) in MeOH (0.25 M) using AcOH (1.0 equiv.). The crude residue was purified by flash column chromatography eluting with 2:1 Pet. Ether:EtOAc to 20:1 EtOAc:MeOH to afford *tert*-butyl (3*R*)-3-[[(3-*tert*-butyl-1*H*-pyrazol-4-yl)methylamino]methyl]pyrrolidine-1-

carboxylate (390 mg, 1.07 mmol, 53 %) as a yellow solid. **$^1$H NMR** (400 MHz, $CDCl_3$) δ 7.49 (s, 1H), 3.80 (br s, 2H), 3.58 - 3.42 (m, 2H), 3.36 - 3.22 (m, 1H), 3.08 - 2.96 (m, 1H), 2.79 - 2.61 (m, 2H), 2.45 - 2.28 (m, 1H), 2.01 (br s, 1H), 1.64 - 1.55 (m, 1H), 1.46 (s, 9H), 1.39 (s, 9H). **MS** (ES+): *m/z* (%) 337.2 (100) $[M+H]^+$.

**(*S*)-1-(3-(*tert*-butyl)-1*H*-pyrazol-4-yl)-*N*-(pyrrolidin-3-ylmethyl) methanamine hydrochloride (Intermediate 2, 14)**

Synthesized according to **general procedure 4** from *tert*-butyl (3*R*)-3-[[(3-*tert*-butyl-1*H*-pyrazol-4-yl)methylamino]methyl]pyrrolidine-1-carboxylate (370 mg, 1.10 mmol, 1.0 equiv.) in $CH_3CN$ (0.28 M) to afford (*S*)-1-(3-(*tert*-butyl)-1*H*-pyrazol-4-yl)-*N*-(pyrrolidin-3-ylmethyl) methanamine hydrochloride (400 mg, crude, HCl salt)as a white solid. **$^1$H NMR** (400 MHz, MeOD) δ 8.54 (s, 1H), 4.47 (s, 2H), 3.65 - 3.60 (m, 1H), 3.52 - 3.45 (m, 1H), 3.40 (br d, *J* = 7.2 Hz, 2H), 3.40 - 3.30 (m, 1H), 3.23 - 3.14 (m, 1H), 2.93 (td, *J* = 8.0, 16.0 Hz, 1H), 2.41 (s, 2H), 2.41 - 2.34 (m, 1H), 1.97 - 1.86 (m, 1H), 1.50 (s, 9H). **MS** (ES+): *m/z* (%) 237.2 (100) $[M+H]^+$.

**(*R*)-1-(3-(*tert*-butyl)-1*H*-pyrazol-4-yl)-*N*-((1-(pyridin-2-ylmethyl)pyrrolidin-3-yl)methyl)methanamine (14)**

Synthesized according to **general procedure 3** from 1-(3-*tert*-butyl-1*H*-pyrazol-4-yl)-*N*-[[(3*R*)-pyrrolidin-3-yl]methyl]methanamine hydrochloride (350 mg, 1.28 mmol, 1.0 equiv., HCl salt) and pyridine-2-carbaldehyde (109 mg, 1.03 mmol, 0.8 equiv.) in MeOH (0.1 M) using AcOH (1.0 equiv.) in addition to NaOAc (2.0 equiv.) and extracted with 10:1 DCM:MeOH. The crude residue was purified by preparative HPLC (21-51 % $CH_3CN$ in water (+1 % $NH_4HCO_3$)) to afford (*R*)-1-(3-(*tert*-butyl)-1*H*-pyrazol-4-yl)-*N*-((1-(pyridin-2-ylmethyl)pyrrolidin-3-yl)methyl)methanamine (49.09 mg, 0.15 mmol, 15 %, 99.9 %ee) as a yellow gum. **$^1$H NMR** (500 MHz, MeOD) δ 8.48 – 8.51 (m, 1H), 7.81 – 7.86 (m, 1H), 7.49 – 7.53 (m, 2H), 7.31 – 7.36 (m, 1H), 3.73 – 3.83 (m, 4H), 2.83 – 2.88 (m, 1H), 2.60 – 2.72 (m, 4H), 2.40 – 2.47 (m, 1H), 2.33 – 2.38 (m, 1H), 2.02 – 2.11 (m, 1H), 1.50 – 1.57 (m, 1H), 1.38 (s, 9H). **$^{13}$C NMR** (126 MHz, MeOD) δ 158.2, 148.2, 137.3, 123.6, 122.5, 61.0, 58.4, 53.9, 53.5, 43.9, 37.2, 29.1, 28.7. **HRMS** (ES+): calcd. for $C_{19}H_{29}N_5$ $[M+H]^+$ 328.2501, found 328.2507 (1.8 ppm).

***tert*-butyl (3*S*)-3-[[(3-*tert*-butyl-1*H*-pyrazol-4-yl)methylamino]methyl]pyrrolidine-1-carboxylate (Intermediate 1,15)**

Synthesized according to **general procedure 3** from *tert*-butyl (3*S*)-3-(aminomethyl)pyrrolidine-1-carboxylate (500 mg, 2.50 mmol, 1.0 equiv.) and 3-*tert*-butyl-1*H*-pyrazole-4-carbaldehyde (303 mg, 2.00 mmol, 0.8 equiv.) to afford *tert*-butyl (3*S*)-3-[[(3-*tert*-butyl-1*H*-pyrazol-4-yl)methylamino]methyl]pyrrolidine-1-carboxylate (380 mg, 1.11 mmol, 56 %) as a white solid. **$^1$H NMR** (400 MHz, $CDCl_3$) δ 7.48 (s, 1H), 3.77 (s, 2H), 3.61 - 3.38 (m, 2H), 3.36 - 3.22 (m, 1H), 3.09 - 2.93 (m, 1H), 2.77 - 2.59 (m, 2H), 2.34 (td, *J* = 7.2, 15.2 Hz, 1H), 2.01 - 1.90 (m , 1H), 1.64 - 1.55 (m, 1H), 1.46 (s, 9H), 1.39 (s, 9H). **MS** (ES+): *m/z* (%) 337.2 (98) $[M+H]^+$.

**(*S*)-1-(3-(*tert*-butyl)-1*H*-pyrazol-4-yl)-*N*-((1-(pyridin-2-ylmethyl)pyrrolidin-3-yl)methyl)methanamine hydrochloride (intermediate 2,15)**

Synthesized according to **general procedure 4** from *tert*-butyl (3*S*)-3-[[(3-*tert*-butyl-1*H*-pyrazol-4-yl)methylamino]methyl]pyrrolidine-1-carboxylate (380 mg, 1.11 mmol, 1.0 equiv.) to afford (*S*)-1-(3-(*tert*-butyl)-1*H*-pyrazol-4-yl)-*N*-((1-(pyridin-2-ylmethyl)pyrrolidin-3-yl)methyl)methanamine hydrochloride (380 mg, crude, HCl salt) as a white solid. **$^1$H NMR** (400 MHz, MeOD) δ 8.62 (s, 1H), 4.49 (s, 2H), 3.66 - 3.60 (m, 1H), 3.52 - 3.45 (m, 1H), 3.41 (br d, *J* = 7.2 Hz, 2H), 3.38 - 3.35 (m, 1H), 3.21 (dd, *J* = 8.4, 11.6 Hz, 1H), 2.95 (td, *J* = 7.6, 15.2 Hz, 1H), 2.42 - 2.35 (m, 1H), 1.98 - 1.90 (m, 1H), 1.51 (s, 9H). **MS** (ES+): *m/z* (%) 237.2 (100) $[M+H]^+$.

**(*S*)-1-(3-(*tert*-butyl)-1*H*-pyrazol-4-yl)-*N*-((1-(pyridin-2-ylmethyl)pyrrolidin-3-yl)methyl)methanamine (15)**

Synthesized according to **general procedure 3** from (*S*)-1-(3-(*tert*-butyl)-1*H*-pyrazol-4-yl)-*N*-((1-(pyridin-2-ylmethyl)pyrrolidin-3-yl)methyl)methanamine hydrochloride (380 mg, 1.67 mmol, 1.0 equiv.) and pyridine-2-carbaldehyde (144 mg, 1.34 mmol, 0.8 equiv.) in MeOH (0.1 M) using AcOH (1.0 equiv.) in addition to NaOAc (2.0 equiv.) and extracted with 10:1 DCM:MeOH. The crude residue was purified by preparative HPLC (21-51 % $CH_3CN$ in water (+1 % $NH_4HCO_3$)) to afford product which was further purified by preparative HPLC (23-53 % $CH_3CN$ in water (+0.05 % $NH_4OH$) to afford (*S*)-1-(3-(*tert*-butyl)-1*H*-pyrazol-4-yl)-*N*-((1-(pyridin-2-ylmethyl)pyrrolidin-3-yl)methyl)methanamine (33 mg, 0.101 mmol, 8 %, 99.9 %ee) as a yellow gum. **$^1$H NMR** (400 MHz, MeOD) δ 8.47 (td, *J* = 0.8, 4.0 Hz, 1H), 7.82 (dt, *J* = 2.0, 8.0 Hz, 1H), 7.50 (br d, *J* = 7.6 Hz, 2H), 7.33 - 7.30 (m, 1H), 3.84 - 3.68 (m, 4H), 2.83 (dd, *J* = 7.6, 9.2 Hz, 1H), 2.71 - 2.56 (m, 4H), 2.47 - 2.36 (m, 1H), 2.36 - 2.28 (m, 1H), 2.10 - 1.98 (m, 1H), 1.54 - 1.49 (m, 1H), 1.36 (s, 9H). **$^{13}$C NMR** (101 MHz, MeOD) δ 158.2, 148.2, 137.3, 123.6, 122.5, 61.0, 58.4, 53.9, 53.5, 43.9, 37.2, 29.1, 28.7. **HRMS** (ES+): calcd. for $C_{19}H_{30}N_5$ $[M+H]^+$ 328.2496, found 328.2499 (1.14 ppm).

***tert*-butyl ((1-(pyridin-2-ylmethyl)piperidin-3-yl)methyl)carbamate hydrochloride (Intermediate 1, 16)**

Synthesized according to **general procedure 3** from *tert*-butyl *N*-(3-piperidylmethyl)carbamate (500 mg, 2.33 mmol, 1.0 equiv.) and pyridine-2-carbaldehyde (249 mg, 2.33 mmol, 1.0 equiv.) in MeOH (0.23 M) using AcOH (1.0 equiv.). The reaction was quenched with water, concentrated *in vacuo* and the residue was purified by reverse phase flash column chromatography (5-95 % $CH_3CN$ in water (+0.1 % FA)) to afford *tert*-butyl *N*-[[1-(2-pyridylmethyl)-3-piperidyl]methyl]carbamate (200 mg, 0.65 mmol, 25 %) as a yellow oil. **$^1$H NMR** (400 MHz, $CDCl_3$) δ 8.60 - 8.53 (m, 1H), 7.65 (dt, *J* = 1.6, 7.6 Hz, 1H), 7.42 (br d, *J* = 5.6 Hz, 1H), 7.21 - 7.13 (m, 1H), 4.75 - 4.57 (m, 1H), 3.65 (br s, 2H), 3.16 - 2.94 (m, 2H), 2.89 - 2.64 (m, 2H), 1.97 - 1.53 (m, 6H), 1.43 (s, 9H), 1.17 - 0.92 (m, 1H). **MS** (ES+): *m/z* (%) 306.3 (100) $[M+H]^+$.

***tert*-butyl ((1-(pyridin-2-ylmethyl)piperidin-3-yl)methyl)carbamate hydrochloride (Intermediate 2, 16)**

Synthesized according to **general procedure 4** from *tert*-butyl *N*-[[1-(2-pyridylmethyl)-3-piperidyl]methyl]carbamate (200 mg, 0.65 mmol 1.0 equiv.) in $CH_3CN$ (0.33 M) to afford *tert*-butyl ((1-(pyridin-2-ylmethyl)piperidin-3-yl)methyl)carbamate hydrochloride (158 mg, 0.65 mmol, >99 %, HCl salt ) as a white solid which was used without further purification. **MS** (ES+): *m/z* (%) 206.2 (100) $[M+H]^+$.

**1-(3-(*tert*-butyl)-1*H*-pyrazol-4-yl)-*N*-((1-(pyridin-2-ylmethyl)piperidin-3-yl)methyl)methanamine (16)**

Synthesized according to **general procedure 3** from [1-(2-pyridylmethyl)-3-piperidyl]methanamine hydrochloride (330 mg, 1.36 mmol, 1.0 equiv.) and 3-*tert*-butyl-1*H*-pyrazole-4-carbaldehyde (186 mg, 1.23 mmol, 0.9 equiv.) in MeOH (0.2 M) using AcOH (1.0 equiv.) in addition to NaOAc (2.0 equiv.).The reaction was quenched with water, concentrated *in vacuo* and purified by preparative HPLC (15-45 % $CH_3CN$ in water (+1 % $NH_4HCO_3$) using Phenomenex Gemini-NX C18 75 * 30 mm, 3 μm column) to afford 1-(3-(*tert*-butyl)-1*H*-pyrazol-4-yl)-*N*-((1-(pyridin-2-ylmethyl)piperidin-3-yl)methyl)methanamine (87 mg, 0.25 mmol, 18 %) as a white solid. **$^1$H NMR** (500 MHz, MeOD) δ 8.47 (d, *J* = 4.8 Hz, 1H), 7.85 - 7.80 (m, 1H), 7.56 - 7.52 (m, 2H), 7.32 (dd, *J* = 5.6, 6.9 Hz, 1H), 3.91 - 3.89 (m, 2H), 3.68 - 3.67 (m, 2H), 2.94 (d, *J* = 7.2 Hz, 1H), 2.81 (d, *J* = 11.1 Hz, 1H), 2.69 (d, *J* = 5.8 Hz, 2H), 2.21 - 2.13 (m, 1H), 1.94 (t, *J* = 6.9 Hz, 2H), 1.86 - 1.80 (m, 1H), 1.76 - 1.60 (m, 2H), 1.37 (s, 9H). **$^{13}$C NMR** (126 MHz, MeOD) δ 157.8, 148.2,

137.2, 123.8, 122.6, 112.5, 63.8, 57.9, 53.9, 52.4, 43.7, 35.0, 31.9, 29.2, 28.3, 24.1. **HRMS** (ES+): calcd. for $C_{20}H_{31}N_5$ $[M+H]^+$ 342.2652, found 342.2661 (2.6 ppm).

***tert*-butyl 3-(((3-(*tert*-butyl)-1*H*-pyrazol-4-yl)methyl)amino)-9-azabicyclo[3.3.1]nonane-9-carboxylate (Intermediate 1, 17)**

Synthesized according to **general procedure 3** from *tert*-butyl 3-amino-9-azabicyclo[3.3.1]nonane-9-carboxylate (500 mg, 2.08 mmol, 1.0 equiv.) and 3-*tert*-butyl-1*H*-pyrazole-4-carbaldehyde (253 mg, 1.66 mmol, 0.8 equiv.) in MeOH (0.21 M) using AcOH (1.0 equiv.). The crude residue was purified by reverse phase flash column chromatography (5-95 % $CH_3CN$ in water (+0.1 % FA)) to afford *tert*-butyl 3-(((3-(*tert*-butyl)-1*H*-pyrazol-4-yl)methyl)amino)-9-azabicyclo[3.3.1]nonane-9-carboxylate (350 mg, 0.93 mmol, 56 %) as a yellow oil. **$^1$H NMR** (400 MHz, MeOD) δ 7.50 (br s, 1H), 4.41 (br d, *J* = 10.4 Hz, 2H), 3.79 (d, *J* = 2.0 Hz, 2H), 2.52 - 2.41 (m, 1H), 2.41 - 2.27 (m, 2H), 2.12 - 2.02 (m, 1H), 1.66 - 1.46 (m, 5H), 1.43 (s, 9H), 1.37 (s, 9H), 1.30 - 1.25 (m, 2H). **MS** (ES+): *m/z* (%) 377.4 (100) $[M+H]^+$.

***N*-((3-(*tert*-butyl)-1*H*-pyrazol-4-yl)methyl)-9-azabicyclo[3.3.1]nonan-3-amine hydrochloride (Intermediate 2, 17)**

Synthesized according to **general procedure 4** from *tert*-butyl 3-(((3-(*tert*-butyl)-1*H*-pyrazol-4-yl)methyl)amino)-9-azabicyclo[3.3.1]nonane-9-carboxylate (340 mg, 0.90 mmol, 1.0 equiv.)*)* in MeOH (0.23 M) to afford *N*-((3-(*tert*-butyl)-1*H*-pyrazol-4-yl)methyl)-9-azabicyclo[3.3.1]nonan-3-amine hydrochloride (300 mg, crude, HCl salt)which was used without further purification. **$^1$H NMR** (400 MHz, MeOD) δ 8.27 (s, 1H), 4.45 (s, 2H), 4.03 - 3.93 (m, 2H), 3.90 - 3.76 (m, 1H), 2.82 (td, *J* = 6.0, 12.0 Hz, 2H), 2.23 - 2.08 (m, 1H), 2.01 - 1.89 (m, 4H), 1.83 - 1.69 (m, 3H), 1.49 (s, 9H). **MS** (ES+): *m/z* (%) 277.2 (100) $[M+H]^+$.

***N*-((3-(*tert*-butyl)-1*H*-pyrazol-4-yl)methyl)-9-(pyridin-2-ylmethyl)-9-azabicyclo[3.3.1]nonan-3-amine (17)**

Synthesized according to **general procedure 3** from *N*-((3-(*tert*-butyl)-1*H*-pyrazol-4-yl)methyl)-9-azabicyclo[3.3.1]nonan-3-amine hydrochloride (280 mg, 0.89 mmol, 1.0 equiv., HCl salt) and pyridine-2-carbaldehyde (96 mg, 0.89 mmol, 1.0 equiv.) in MeOH (0.1 M) using AcOH (1.0 equiv.) in addition to NaOAc (2.0 equiv.). . The crude residue was purified by preparative HPLC (1-40 % EtOH in hexane (+0.1 % $NH_4OH$) using Welch Ultimate XB-CN 250 * 50 mm, 10 µm column) followed by preparative HPLC (32-62 % $CH_3CN$ in water (+1 % $NH_4HCO_3$)) to afford *N*-((3-(*tert*-butyl)-1*H*-pyrazol-4-yl)methyl)-9-(pyridin-2-ylmethyl)-9-azabicyclo[3.3.1]nonan-3-amine (74 mg, 0.19 mmol, 22 %) as a yellow solid. **$^1$H NMR** (500 MHz, MeOD) δ 8.45 - 8.42 (m, 1H), 7.85 - 7.80 (m, 1H), 7.60 - 7.57 (m, 2H), 7.31 - 7.27 (m, 1H), 3.98 (s, 2H), 3.89 (s, 2H), 3.10 (d, *J* = 10.9 Hz, 2H), 2.49 - 2.40 (m, 2H), 2.15 - 1.97 (m, 3H), 1.57 (d, *J* = 13.0 Hz, 1H), 1.49 - 1.44 (m, 1H), 1.42 - 1.40 (m, 9H), 1.37 - 1.29 (m, 2H), 1.11 (br d, *J* = 12.7 Hz, 2H). **$^{13}$C NMR** (126 MHz, MeOD) δ 160.6, 147.8, 137.4, 122.8, 122.1, 114.2, 57.0, 50.1, 49.0, 41.0, 32.2, 29.3, 24.9, 13.8. **HRMS** (ES+): calcd. for $C_{22}H_{33}N_5$ $[M+H]^+$ 368.2809, found 368.2806 (0.7 ppm).

***tert*-butyl 3-(((3-(*tert*-butyl)-1*H*-pyrazol-4-yl)methyl)amino)-8-azabicyclo[3.2.1]octane-8-carboxylate (Intermediate 1, 18)**

Synthesized according to **general procedure 3** from *tert*-butyl 3-amino-8-azabicyclo[3.2.1]octane-8-carboxylate (500 mg, 2.21 mmol, 1.0 equiv.) and 3-*tert*-butyl-1*H*-pyrazole-4-carbaldehyde (253 mg, 1.66 mmol, 0.8 equiv.) in MeOH (0.21 M) to afford *tert*-butyl 3-(((3-(*tert*-butyl)-1*H*-pyrazol-4-yl)methyl)amino)-8-azabicyclo[3.2.1]octane-8-carboxylate (360 mg, 0.99 mmol, 60 %) as a white solid. **$^1$H NMR** (400 MHz, MeOD) δ 7.61 - 7.31 (m, 1H), 4.13 (br s, 2H), 3.73 (s, 2H), 3.03 - 2.89 (m, 1H), 2.23 - 2.02 (m, 4H), 1.90 (br s, 2H), 1.64 (br d, J = 14.4 Hz, 2H), 1.46 (s, 9H), 1.38 (br s, 9H). **MS** (ES+): m/z (%) 363.3 (100) $[M+H]^+$.

***N*-((3-(*tert*-butyl)-1*H*-pyrazol-4-yl)methyl)-8-azabicyclo[3.2.1]octan-3-amine hydrochloride (Intermediate 2, 18)**

Synthesized according to **general procedure 4** from *tert*-butyl 3-(((3-(*tert*-butyl)-1*H*-pyrazol-4-yl)methyl)amino)-8-azabicyclo[3.2.1]octane-8-carboxylate (360 mg, 0.90 mmol, 1.0 equiv.) in MeOH (0.23 M) to afford the desired produce as a white solid (300 mg, crude, HCl) which was used without further purification. **$^1$H NMR** (400 MHz, MeOD) δ 8.50 (s, 1H), 4.48 (s, 2H), 4.15 (br d, *J* = 5.6 Hz, 2H), 3.87 - 3.70 (m, 1H), 2.80 (td, *J* = 7.6, 15.2 Hz, 2H), 2.31 - 2.18 (m, 6H), 1.50 (s, 9H). **MS** (ES+): *m/z* (%) 263.2 (100) $[M+H]^+$.

***N*-((3-(*tert*-butyl)-1*H*-pyrazol-4-yl)methyl)-8-(pyridin-2-ylmethyl)-8-azabicyclo[3.2.1]octan-3-amine (18)**

Synthesized according to **general procedure 3** from *N*-((3-(*tert*-butyl)-1*H*-pyrazol-4-yl)methyl)-8-azabicyclo[3.2.1]octan-3-amine hydrochloride (300 mg, 0.99 mmol, 1.0 equiv., HCl salt) and pyridine-2-carbaldehyde (96 mg, 0.89 mmol, 1.0 equiv.) in MeOH (0.1 M) to afford *N*-((3-(*tert*-butyl)-1*H*-pyrazol-4-yl)methyl)-8-(pyridin-2-ylmethyl)-8-azabicyclo[3.2.1]octan-3-amine (40 mg, 0.11 mmol, 11 %) as a light yellow solid. **$^1$H NMR** (500 MHz, MeOD) δ 8.49 - 8.46 (m, 1H), 7.86 - 7.81 (m, 1H), 7.69 - 7.65 (m, 1H), 7.46 - 7.43 (m, 1H), 7.34 - 7.30 (m, 1H), 3.75 - 3.72 (m, 4H), 3.25 - 3.21 (m, 2H), 3.00 (t, *J* = 6.4 Hz, 1H), 2.20 - 2.06 (m, 6H), 1.66 (d, *J* = 14.3 Hz, 2H), 1.40 (s, 9H). **$^{13}$C NMR** (126 MHz, MeOD) δ 159.0, 148.1, 137.3, 123.4, 122.4, 58.1, 57.4, 49.9, 43.2, 35.8, 29.1, 26.1. **HRMS** (ES+): calcd. for $C_{21}H_{31}N_5$ $[M+H]^+$ 354.2652, found 354.2658 (1.5 ppm).

***tert*-butyl (1-(pyridin-2-ylmethyl)azepan-4-yl)carbamate (Intermediate 1, 19)**

Synthesized according to **general procedure 3** from *tert*-butyl azepan-4-ylcarbamate hydrohcloride (500 mg, 1.99 mmol, 1.0 equiv., HCl salt) and pyridine-2-carbaldehyde (213 mg, 1.99 mmol, 1.0 equiv.) in MeOH (0.4 M) using AcOH (1.0 equiv.) in addition to NaOAc (2.0 equiv.) and extracted with 10:1 DCM:MeOH to afford *tert*-butyl (1-(pyridin-2-ylmethyl)azepan-4-yl)carbamate (500 mg, crude, 1.64 mmol, 78 %) as a yellow oil which was used without further purification. **MS** (ES+): *m/z* (%) 306.4 (100) $[M+H]^+$.

**1-(pyridin-2-ylmethyl)azepan-4-amine hydrochloride (Intermediate 2, 19)**

Synthesized according to **general procedure 4** from *tert*-butyl (1-(pyridin-2-ylmethyl)azepan-4-yl)carbamate (500 mg, 1.64 mmol, 1.0 equiv.) *in* $CH_3CN$ (0.33 M) to afford 1-(pyridin-2-ylmethyl)azepan-4-amine hydrochloride (400 mg, crude, HCl salt) as a yellow solid which was used without further purification. **$^1$H NMR** (400 MHz, MeOD) δ 8.77 (dd, *J* = 0.8, 5.2 Hz, 1H), 8.12 (dt, *J* = 1.6, 7.6 Hz, 1H), 7.80 (d, *J* = 8.0 Hz, 1H), 7.65 (dt, *J* = 0.8, 6.0 Hz, 1H), 4.67 (s, 2H), 3.64 - 3.60 (m, 1H), 3.59 - 3.46 (m, 4H), 2.35 - 2.20 (m, 3H), 2.15 - 2.11 (m, 1H), 2.09 - 2.01 (m, 1H), 1.83 - 1.79 (m, 1H). **MS** (ES+): *m/z* (%) 206.2 (100) $[M+H]^+$.

***N*-((3-(*tert*-butyl)-1*H*-pyrazol-4-yl)methyl)-1-(pyridin-2-ylmethyl)azepan-4-amine hydrochloride (19)**

Synthesized according to **general procedure 3** from 1-(2-pyridylmethyl)azepan-4-amine hydrochloride (400 mg, 1.65 mmol, 1.0 equiv.) and 3-*tert*-butyl-1*H*-pyrazole-4-carbaldehyde (176 mg, 1.16 mmol, 0.7 equiv.) in MeOH (0.3 M) using AcOH (1.0 equiv.) in addition to NaOAc (2.0 equiv.) and extracted with 10:1 DCM:MeOH.The crude residue was purified by preparative HPLC (1-21 % $CH_3CN$ in water (+0.05 % HCl) using 3_Phenomenex Luna C18 75 * 30 mm,3 µm column) to afford *N*-((3-(*tert*-butyl)-1*H*-pyrazol-4-yl)methyl)-1-(pyridin-2-ylmethyl)azepan-4-amine hydrochloride (127 mg, 0.37 mmol, 32 %, HCl salt) as a white solid. **$^1$H NMR** (500 MHz, MeOD) δ 8.93 (d, *J* = 5.0 Hz, 1H), 8.60 (s, 1H), 8.46 (t, *J* = 7.6 Hz, 1H), 8.20 (d, *J* = 7.6 Hz, 1H), 7.96 (t, *J* = 6.3 Hz, 1H), 4.88 - 4.85 (m, 2H), 3.89 - 3.80 (m, 2H), 3.70 - 3.57 (m, 3H), 2.74 - 2.68 (m, 1H), 2.61 - 2.50 (m, 2H), 2.32 - 2.22 (m, 1H), 2.17 - 2.05 (m, 2H), 1.54 (s, 9H). **$^{13}$C NMR** (126 MHz, MeOD) δ 154.7,

146.0, 145.9, 143.6, 136.1, 128.1, 126.7, 110.7, 57.6, 54.9, 50.5, 39.2, 32.7, 28.7, 28.6, 25.6, 19.7. **HRMS** (ES+): calcd. for $C_{20}H_{31}N_5$ $[M+H]^+$ 342.2652, found 342.2658 (1.6 ppm).

**1-benzyl-*N*-((3-(*tert*-butyl)-1*H*-pyrazol-4-yl)methyl)piperidin-4-amine (20)**

Synthesized according to **general procedure 3** from 1-benzylpiperidin-4-amine (375 mg, 1.97 mmol, 402 µL, 1.5 equiv.) and 3-*tert*-butyl-1*H*-pyrazole-4-carbaldehyde (200 mg, 1.31 mmol, 1.0 equiv.) in EtOH (0.3 M). The crude residue was purified by preparative HPLC (32-62 % $CH_3CN$ in water (+1 % $NH_4HCO_3$)) to afford 1-benzyl-*N*-((3-(*tert*-butyl)-1*H*-pyrazol-4-yl)methyl)piperidin-4-amine (14 mg, 0.041 mmol, 3 %) as colorless gum. **$^1$H NMR** (400 MHz, MeOD) δ 7.57 - 7.45 (m, 1H), 7.32 (d, *J* = 4.4 Hz, 4H), 7.29 - 7.25 (m, 1H), 3.80 (s, 2H), 3.52 (s, 2H), 2.92 (br d, *J* = 11.6 Hz, 2H), 2.65 - 2.52 (m, 1H), 2.12 - 2.03 (m, 2H), 1.99 - 1.90 (m, 2H), 1.55 - 1.42 (m, 2H), 1.37 - 1.36 (s, 9H). **HRMS** (ES+): calcd. For $C_{20}H_{31}N_4$ $[M+H]^+$ 327.2544, found 327.2547 (1.29 ppm).

***N*-((3-(*tert*-butyl)-1*H*-pyrazol-4-yl)methyl)-1-((3-methoxypyridin-2-yl)methyl)piperidin-4-amine (21)**

Synthesized according to **general procedure 5** from *N*-((3-(*tert*-butyl)-1*H*-pyrazol-4-yl)methyl)piperidin-4-amine (100 mg, 0.42 mmol, 1.0 equiv.) and 3-methoxypyridine-2-carbaldehyde (58 mg, 0.42 mmol, 1.0 equiv.) in EtOH (0.4 M) to afford *N*-((3-(*tert*-butyl)-1*H*-pyrazol-4-yl)methyl)-1-((3-methoxypyridin-2-yl)methyl)piperidin-4-amine (37 mg, 0.098 mmol, 23 %) as a clear sticky glass. **$^1$H NMR** (500 MHz, MeOD) δ 8.13 – 8.09 (m, 1H), 7.53 (br s, 1H), 7.46 (d, *J* = 8.2 Hz, 1H), 7.37 – 7.32 (m 1H), 3.9 (s, 3H), 3.85 (s, 2H), 3.74, (s 2H), 3.06 – 3.01 (m, 2H), 2.68 – 2.60 (m, 1H), 2.29 – 2.21 (m, 2H), 1.99 – 1.92 (m, 2H), 1.59 – 1.49 (m, 2H), 1.39 (s, 9H). **$^{13}$C NMR** (126 MHz, MeOD) δ 155.0, 146.2, 139.5, 123.8, 118.3, 57.0, 54.7, 54.4, 52.2, 40.6, 30.8, 29.1. **HRMS** (ES+): calcd. for $C_{20}H_{32}N_5O$ $[M+H]^+$ 358.2601, found 358.2600 (0.5 ppm).

***N*-((3-(*tert*-butyl)-1*H*-pyrazol-4-yl)methyl)-1-((4-methoxypyridin-2-yl)methyl)piperidin-4-amine (22)**

Synthesized according to **general procedure 5** from *N*-((3-(*tert*-butyl)-1*H*-pyrazol-4-yl)methyl)piperidin-4-amine (100 mg, 0.42 mmol, 1.0 equiv.) and 4-methoxypicolinaldehyde (58 mg, 0.42 mmol, 1.0 equiv.) in EtOH (0.4 M) to afford *N*-((3-(*tert*-butyl)-1*H*-pyrazol-4-yl)methyl)-1-((4-methoxypyridin-2-yl)methyl)piperidin-4-amine (9 mg, 0.023 mmol, 5.7 %) as a clear, sticky glass. **$^1$H NMR** (500 MHz, DMSO-$d_6$) δ 12.07 (s, 1H), 8.30 - 8.28 (m, 1H), 7.41 - 7.34 (m, 1H), 6.97 (d, J = 2.4 Hz, 1H), 6.84 (dd, *J* =2.6, 5.8 Hz, 1H), 3.82 (s, 3H), 3.66 (s, 2H), 3.52 - 3.50 (m, 2H), 2.81 - 2.77 (m, 2H), 2.45 (s, 1H), 2.08 - 2.03 (m, 2H), 1.84 - 1.81 (m, 2H), 1.32 - 1.30 (m, 11H). **HRMS** (ES+): calcd. For $C_{20}H_{32}ON_5$ $[M+H]^+$ 358.2602, found 358.2606 (1.36 ppm).

***N*-((3-(*tert*-butyl)-1*H*-pyrazol-4-yl)methyl)-1-((5-methoxypyridin-2-yl)methyl)piperidin-4-amine (23)**

Synthesized according to **general procedure 5** from *N*-((3-(*tert*-butyl)-1*H*-pyrazol-4-yl)methyl)piperidin-4-amine (100 mg, 0.42 mmol, 1.0 equiv.) and 5-methoxypicolinaldehyde (58 mg, 0.42 mmol, 1.0 equiv.) in EtOH (0.8 M) to afford 5-methoxypicolinaldehyde (58 mg, 0.42 mmol, 1.0 equiv.) in EtOH (0.8 M) to afford *N*-((3-(*tert*-butyl)-1*H*-pyrazol-4-yl)methyl)-1-((5-methoxypyridin-2-yl)methyl)piperidin-4-amine (19 mg, 0.050 mmol, 12 %) as a clear, sticky glass. **$^1$H NMR** (500 MHz, MeOD) δ 8.18 – 8.16 (m, 1H), 7.53 (br s, 1H), 7.47 (d, *J* = 8.4 Hz, 1H), 7.43 – 7.39 (m, 1H), 3.88 (s, 3H), 3.81 (s, 2H), 3.60, (s 2H), 2.94 – 2.89 (m, 2H), 2.64 – 2.56 (m, 1H), 2.18 – 2.11 (m, 2H), 1.98 – 1.92 (m, 2H), 1.55 – 1.46 (m, 2H), 1.38 (s, 9H). **$^{13}$C NMR** (126 MHz, MeOD) δ 155.4, 149.5, 135.6, 124.4, 121.6, 62.7, 54.9, 54.5, 52.1, 40.7, 31.3, 29.1. **HRMS** (ES+): calcd. for $C_{20}H_{32}N_5O$ $[M+H]^+$ 358.2601, found 358.2590 (3.1 ppm).

***N*-((3-(*tert*-butyl)-1*H*-pyrazol-4-yl)methyl)-1-((6-methoxypyridin-2-yl)methyl)piperidin-4-amine (24)**

Synthesized according to **general procedure 5** from *N*-((3-(*tert*-butyl)-1*H*-pyrazol-4-yl)methyl)piperidin-4-amine (100 mg, 0.42 mmol, 1.0 equiv.) and 6-methoxypyridine-2-carbaldehyde (58 mg, 0.42 mmol, 1.0 equiv.) in EtOH (0.8 M) to *N*-((3-(*tert*-butyl)-1*H*-pyrazol-4-yl)methyl)-1-((6-methoxypyridin-2-yl)methyl)piperidin-4-amine (16 mg, 0.043 mmol, 10 %) as a sticky, clear solid. **$^1$H NMR** (500 MHz, MeOD) δ 7.65 – 7.60 (m, 1H), 7.53 (br s, 1H), 7.01 (d, *J* = 7.2 Hz, 1H), 6.67 (d, J = 8.2 Hz, 1H), 3.92 (s, 3H), 3.81 (s, 2H), 3.58 (s 2H), 3.01 – 2.96 (m, 2H), 2.63 – 2.56 (m, 1H), 2.23 – 2.16 (m, 2H), 1.99 – 1.93 (m, 2H), 1.58 – 1.48 (m, 2H), 1.39 (s, 9H). **$^{13}$C NMR** (126 MHz, MeOD) δ 163.9, 155.5, 138.9, 116.0, 108.2, 63.4, 54.5, 52.5, 52.3, 40.8, 31.3, 29.1. **HRMS** (ES+): calcd. for $C_{20}H_{32}N_5O$ [M+H]$^+$ 358.2601, found 358.2600 (0.4 ppm).

***N*-((3-(*tert*-butyl)-1*H*-pyrazol-4-yl)methyl)-1-((3-methylpyridin-2-yl)methyl)piperidin-4-amine, (25)**

Synthesized according to **general procedure 5** from *N*-((3-(*tert*-butyl)-1*H*-pyrazol-4-yl)methyl)piperidin-4-amine (100 mg, 0.42 mmol, 1.0 equiv.) and 3-methylpyridine-2-carbaldehyde (51 mg, 0.42 mmol, 1.0 equiv.) in EtOH (0.8 M) to afford *N*-((3-(*tert*-butyl)-1*H*-pyrazol-4-yl)methyl)-1-((3-methylpyridin-2-yl)methyl)piperidin-4-amine (17 mg, 0.047 mmol, 11 %) as a sticky, clear oil. **$^1$H NMR** (500 MHz, MeOD) δ 8.31 – 8.27 (m, 1H), 7.66 – 7.61 (m, 1H), 7.53 (br s, 1H), 7.27 - 7.23 (m, 1H), 3.80 (s, 2H), 3.65 (s, 2H), 2.91 – 2.85 (m, 2H), 2.63 – 2.56 (m, 1H), 2.46 (s, 3H), 2.22 – 2.14 (m, 2H), 1.96 – 1.88 (m, 2H), 1.50 – 1.41 (m, 2H), 1.39 (s, 9H). **$^{13}$C NMR** (126 MHz, MeOD) δ 156.1, 144.9, 138.8, 134.2, 122.7, 61.8, 54.7, 52.4, 40.8, 31.6, 29.1, 17.1. **HRMS** (ES+): calcd. for $C_{20}H_{32}N_5$ [M+H]+ 342.2652, found 342.2641 (3.3 ppm).

***N*-((3-(*tert*-butyl)-1*H*-pyrazol-4-yl)methyl)-1-((3-fluoropyridin-2-yl)methyl)piperidin-4-amine (26)**

Synthesized according to **general procedure 5** from *N*-((3-(*tert*-butyl)-1*H*-pyrazol-4-yl)methyl)piperidin-4-amine (100mg, 0.42 mmol, 1.0 equiv.) and 3-fluoropicolinaldehyde (53 mg, 0.42 mmol, 1.0 equiv.) in EtOH (0.8 M) to afford *N*-((3-(*tert*-butyl)-1*H*-pyrazol-4-yl)methyl)-1-((3-fluoropyridin-2-yl)methyl)piperidin-4-amine (7 mg, 0.019 mmol, 4.5 %) as a sticky, clear glass**. $^1$H NMR** (500 MHz, MeOD) δ 8.40 - 8.39 (m, 1H), 7.64 - 7.59 (m, 1H), 7.52 (s, 1H), 7.44 - 7.40 (m, 1H), 3.81 (s, 2H), 3.76 (d, J = 2.6 Hz, 2H), 2.99 (d, *J* = 12.1 Hz, 2H), 2.62 - 2.56 (m, 1H), 2.27 - 2.20 (m, 2H), 1.95 - 1.91 (m, 2H), 1.55 - 1.46 (m, 2H), 1.40 - 1.37 (m, 9H). **HRMS** (ES+): calcd. for $C_{19}H_{28}N_5F$ [M+H]+ 346.2402, found 346.2405 (1.06 ppm).

**2-((4-(((3-(*tert*-butyl)-1*H*-pyrazol-4-yl)methyl)amino)piperidin-1-yl)methyl)pyridin-3-ol (27)**

Synthesized according to **general procedure 2** from from *N*-((3-(*tert*-butyl)-1*H*-pyrazol-4-yl)methyl)piperidin-4-amine (586 mg, 2.47 mmol, 1.0 equiv.) and 3-hydroxypicolinaldehyde (305 mg, 2.47 mmol, 1.0 equiv.) in DCM (0.25 M) using $NaBH(OAc)_3$ (1.5 equiv.). The residue was purified by preparative HPLC (5-95 % $CH_3CN$ in water (+0.1 % $NH_4OH$)) to afford 2-((4-(((3-(*tert*-butyl)-1*H*-pyrazol-4-yl)methyl)amino)piperidin-1-yl)methyl)pyridin-3-ol (18 mg, 0.05 mmol, 33 %). **$^1$H NMR** (500 MHz, MeOD) δ 7.95 – 7.92 (m, 1H), 7.54 (s, 1H), 7.23 – 7.19 (m, 1H), 7.18 – 7.15 (m, 1H), 3.91 (s, 2H), 3.82 (s, 2H), 3.05 – 2.99 (m, 2H), 2.73 – 2.66 (m, 1H), 2.35 – 2.28 (m, 2H), 2.07 – 2.01 (m, 2H), 1.58 – 1.49 (m, 2H), 1.39 (s, 9H). **$^{13}$C NMR** (126 MHz, MeOD) δ 154.9, 142.5, 138.7, 123.8, 123.4, 114.6, 61.9, 53.9, 51.8, 40.9, 31.4, 29.1. **HRMS** (ES+): calcd. for $C_{19}H_{30}N_5O$ [M+H]$^+$ 344.2445, found 344.2461 (4.7 ppm).

***N*-((3-(*tert*-butyl)-1*H*-pyrazol-4-yl)methyl)-1-((3-isopropoxypyridin-2-yl)methyl)piperidin-4-amine (28)**

To a solution of 2-((4-(((3-(*tert*-butyl)-1*H*-pyrazol-4-yl)methyl)amino)piperidin-1-yl)methyl)pyridin-3-ol (115 mg, 0.33 mmol, 1.0 equiv.) in DMF (0.17 M) was added $K_2CO_3$ (93 mg, 0.67 mmol, 2.0 equiv.) and 2-bromopropane (45 mg, 0.37 mmol, 1.1 equiv.). The reaction heated in a sealed tube at 80 °C for 16 h and then diluted with MeOH and transferred to SCX washing with MeOH and eluting product with $NH_3$ in MeOH. The combined organics were concentrated *in vacuo* and purified by preparative HPLC (5-95 % $CH_3CN$ in water (+0.1% $NH_4OH$)) to afford *N*-((3-(*tert*-butyl)-1*H*-pyrazol-4-yl)methyl)-1-((3-isopropoxypyridin-2-yl)methyl)piperidin-4-amine (31 mg, 0.076 mmol, 22 %). **$^1$H NMR** (500 MHz, MeOD) δ 7.90 – 7.87 (m, 1H), 7.32 (broad s, 1H), 7.27 – 7.22 (m, 2H), 7.13 – 7.08 (m, 2H), 4.53 – 4.46 (m, 1H), 3.61, (2, 2H), 3.52 (s, 2H), 2.86 – 2.80 (m, 2H), 2.41 – 2.33 (m, 1H), 2.08 – 2.01 (m, 2H), 1.76 – 1.70 (m, 2H), 1.35 – 1.26 (m, 2H), 1.20 – 1.15 (m, 15H). **$^{13}$C NMR** (126 MHz, MeOD) δ 153.2, 147.2, 139.4, 123.5, 120.5, 70.5, 56.9, 54.4, 52.2, 40.7, 31.2, 29.1, 20.8.F **HRMS** (ES+): calcd. for $C_{22}H_{36}N_5O_1$ $[M+H]^+$ 386.292, found 386.2932 (3.1 ppm).

**N-((3-(tert-butyl)-1H-pyrazol-4-yl)methyl)-1-((3-ethoxypyridin-2-yl)methyl)piperidin-4-amine (29)**

Synthesized according to **general procedure 2** from *N*-((3-(*tert*-butyl)-1*H*-pyrazol-4-yl)methyl)piperidin-4-amine (50 mg, 0.21 mmol, 1.0 equiv.) and 3-ethoxypicolinaldehyde (32 mg, 0.21 mmol, 1.0 equiv.) in DCM (0.2 M) using $NaBH(OAc)_3$ (1.5 equiv.) to afford *N*-((3-(*tert*-butyl)-1*H*-pyrazol-4-yl)methyl)-1-((3-ethoxypyridin-2-yl)methyl)piperidin-4-amine (33 mg, 0.08 mmol, 40 %). **$^1$H NMR** (500 MHz, MeOD) δ 8.11 – 8.08 (m, 1H), 7.52 (br s, 1H), 7.46 – 7.41 (m, 1H), 7.34 – 7.30 (m, 1H), 4.16 – 4.11 (m, 2H), 3.82 (br s, 2H), 3.75 (br s, 2H), 3.08 – 3.01 (m, 2H), 2.63 – 2.56 (m, 1H), 2.29 – 2.21 (m, 2H), 1.98 – 1.90 (m, 2H), 1.57 – 1.49 (m, 2H), 1.49 – 1.43 (m, 3H), 1.39 (br s, 9H). **$^{13}$C NMR** (126 MHz, MeOD) δ 152.9, 144.9, 137.9, 122.2, 117.7, 62.3, 55.4, 52.9, 50.7, 39.3, 29.5, 27.6, 12.1. **HRMS** (ES+): calcd. for $C_{22}H_{36}N_5O_1$ $[M+H]^+$ 372.2763, found 372.2763 (3.2 ppm).

**1-((1H-benzo[d]imidazol-7-yl)methyl)-N-((3-(tert-butyl)-1H-pyrazol-4-yl)methyl)piperidin-4-amine (30)**

Synthesized according to **general procedure 2** from *N*-((3-(*tert*-butyl)-1*H*-pyrazol-4-yl)methyl)piperidin-4-amine (50 mg, 0.21 mmol, 1.0 equiv.) and 3*H*-benzimidazole-4-carbaldehyde (31 mg, 0.21 mmol, 1.0 equiv.) in DCM (0.2 M) using $NaBH(OAc)_3$ (1.5 equiv.) to afford 1-((1*H*-benzo[d]imidazol-7-yl)methyl)-*N*-((3-(*tert*-butyl)-1*H*-pyrazol-4-yl)methyl)piperidin-4-amine (25 mg, 0.06 mmol, 31 %). **$^1$H NMR** (500 MHz, MeOD) δ 8.18 (s, 1H), 7.61 – 7.54 (m, 2H), 7.29 – 7.21 (m, 2H), 3.98 – 3.88 (m, 4H), 3.07 – 2.97 (m, 2H), 2.85 – 2.76 (m, 1H), 2.27 – 2.16 (m, 2H), 2.05 – 1.97 (m, 2H), 1.65 – 1.55 (m 2H), 1.39, (s, 9H). **$^{13}$C NMR** (126 MHz, MeOD) δ 141.1, 123.3, 122.1, 58.2, 54.8, 51.8, 40.5, 30.4, 29.1. **HRMS** (ES+): calcd. for $C_{21}H_{31}N_6$ $[M+H]^+$ 367.261, found 367.2609 (0.3 ppm).

**1-((1H-imidazol-5-yl)methyl)-N-((3-(tert-butyl)-1H-pyrazol-4-yl)methyl)piperidin-4-amine (31)**

Synthesized according to **general procedure 3** from *N*-((3-(*tert*-butyl)-1*H*-pyrazol-4-yl)methyl)piperidin-4-amine (0.2 g, 0.846 mmol, 1.0 equiv.) and 1*H*-imidazole-5-carbaldehyde (81 mg, 0.85 mmol, 1.0 equiv.) in MeOH (0.2 M) and stirred for 0.5 h prior to adding $NaBH_3CN$. The reaction was purified without workup by preparative HPLC (7-37 % $CH_3CN$ in water (+1% $NH_4HCO_3$)) followed by 1-20 % $CH_3CN$ in water (+0.1 % FA) using Waters Atlantis T3 150 * 30 mm, 5 µm column and finally 11-41 % $CH_3CN$ in water (+ 0.1% $NH_4OH$) to afford 1-((1*H*-imidazol-5-yl)methyl)-*N*-((3-(*tert*-butyl)-1*H*-pyrazol-4-yl)methyl)piperidin-4-amine (44.2 mg, 0.14 mmol, 16 %) as a light-yellow gum. **$^1$H NMR** (500 MHz, MeOD) δ 7.61 (s, 1H), 7.50 (br s, 1H), 6.98 (s, 1H), 3.78 (s, 2H), 3.54, (s, 2H), 2.97 – 2.91 (m, 2H), 2.59 – 2.52 (m, 1H), 2.15 – 2.07 (m, 2H), 1.98 – 1.91 (m, 2H), 1.52 – 1.41 (m, 2H), 1.36 (s, 9H). **$^{13}$C NMR** (126 MHz, MeOD) δ 134.8, 54.4, 51.6, 40.8, 31.2, 29.1. **HRMS** (ES+): calcd. for $C_{17}H_{28}N_6$ $[M+H]^+$ 317.2454, found 317.2456 (0.6 ppm).

**Preparation of tert-butyl 4-(((3-(tert-butyl)-1H-pyrazol-4-yl)methyl)amino)azepane-1-carboxylate (Intermediate 1, 32)**

Synthesized according to **general procedure 3** from *tert*-butyl 4-aminoazepane-1-carboxylate (6.97 g, 32.5 mmol, 1.1 equiv.*)* and *tert*-butyl-1*H*-pyrazole-4-carbaldehyde (4.50 g, 29.5 mmol, 1.0 equiv.*)* in MeOH (0.3 M) using AcOH (1.0 equiv.). The product was purified by reverse phase flash chromatography (5-95 % $CH_3CN$ in water (0.1% $NH_4OH$)) to afford *tert*-butyl 4-[(3-*tert*-butyl-1*H*-pyrazol-4-yl)methylamino]azepane-1-carboxylate (9.1 g, 26.0 mmol, 88 %) as a yellow solid. **$^1$H NMR** (400 MHz, $CDCl_3$) δ 7.49 (s, 1H), 3.85 - 3.67 (m, 2H), 3.60 - 3.41 (m, 2H), 3.41 - 3.17 (m, 2H), 2.81 - 2.65 (m, 1H), 2.02 - 1.82 (m, 3H), 1.65 - 1.44 (m, 12H), 1.38 (s, 9H). **MS** (ES+): *m/z* (%) 351.4 (100) $[M+H]^+$.

***N*-((3-(*tert*-butyl)-1*H*-pyrazol-4-yl)methyl)azepan-4-amine (Intermediate 2, 32)**

Synthesized according to **general procedure 4** from *tert*-butyl 4-[(3-*tert*-butyl-1*H*-pyrazol-4-yl)methylamino]azepane-1-carboxylate (9.0 g, 25.6 mmol, 1.0 equiv.) in $CH_3CN$ (0.28 M). The residue was dissolved with water and the mixture was worked up with an anion exchange resin. The resulting residue was washed with water and the filtrate was lyophilized to afford *N*-((3-(*tert*-butyl)-1*H*-pyrazol-4-yl)methyl)azepan-4-amine (6.5 g, 24.1 mmol, 94 %) as a yellow gum. **$^1$H NMR** (400 MHz, MeOD) δ 7.53 (s, 1H), 3.80 (s, 2H), 3.31-3.17 (m, 1H), 3.13 - 2.90 (m, 4H), 2.16 - 1.94 (m, 3H), 1.82 - 1.60 (m, 3H), 1.38 (s, 9H). **MS** (ES+): *m/z* (%) 251.2 (93) $[M+H]^+$.

***N*-((3-(*tert*-butyl)-1*H*-pyrazol-4-yl)methyl)-1-((3-methoxypyridin-2-yl)methyl)azepan-4-amine (32)**

Synthesized according to **general procedure 3** from *N*-((3-(*tert*-butyl)-1*H*-pyrazol-4-yl)methyl)piperidin-4-amine (200 mg, 0.798 mmol, 1.0 equiv.) and 3-methoxypyridine-2-carbaldehyde (109 mg, 0.79 mmol, 1.0 equiv.) in MeOH (0.16 M). The reaction was purified without workup by preparative HPLC (23-53 % $CH_3CN$ in water (+1 % $NH_4HCO_3$)) to afford *N*-((3-(*tert*-butyl)-1*H*-pyrazol-4-yl)methyl)-1-((3-methoxypyridin-2-yl)methyl)azepan-4-amine (80 mg, 0.21 mmol, 26 %) as a brown oil. **$^1$H NMR** (500 MHz, MeOD) δ 8.0 – 7.97 (m, 1H), 7.55 (b s, 1H), 7.45 – 7.41 (m, 1H), 7.33 – 7.29 (m, 1H), 3.87 (s, 4H), 3.85 (s, 3H), 3.06 – 2.99 (m, 1H), 2.99 – 2.93 (m, 1H), 2.84 – 2.79 (m, 2H), 2.77 – 2.70 (m, 1H), 2.01 – 1.89 (m 2H), 1.86 – 1.75 (m, 2H), 1.74 – 1.62 (m, 2H), 1.37 (2, 9H). **$^{13}$C NMR** (126 MHz, MeOD) δ 154.8, 147.2, 139.4, 123.6, 118.2, 57.1, 56.9, 55.2, 54.7, 50.9, 41.1, 32.4, 31.1, 29.1, 23.9. **HRMS** (ES+): calcd. for $C_{22}H_{36}N_5O_1$ $[M+H]^+$ 372.2763, found 372.2773 (2.7 ppm).

**[1]H NMR spectra of intermediate R1.HCl in DMSO-d6**

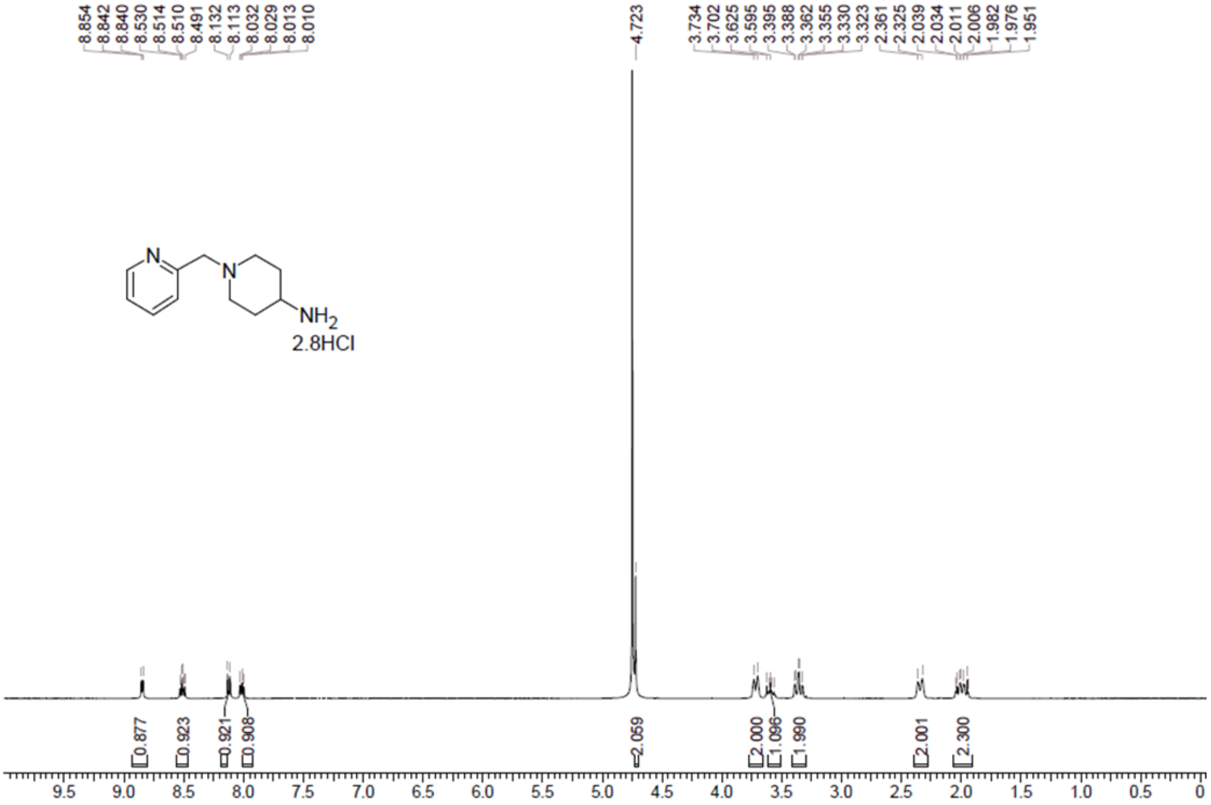

**[1]H NMR spectra of intermediate R1 in MeOD**

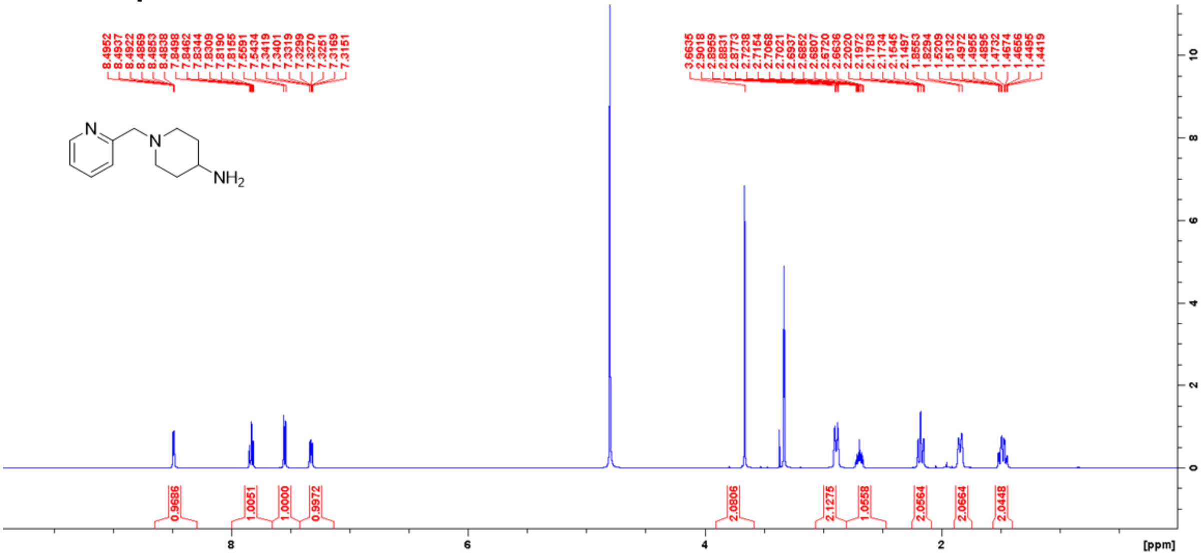

### [1]H NMR spectra of intermediate R2.HCl in $D_2O$

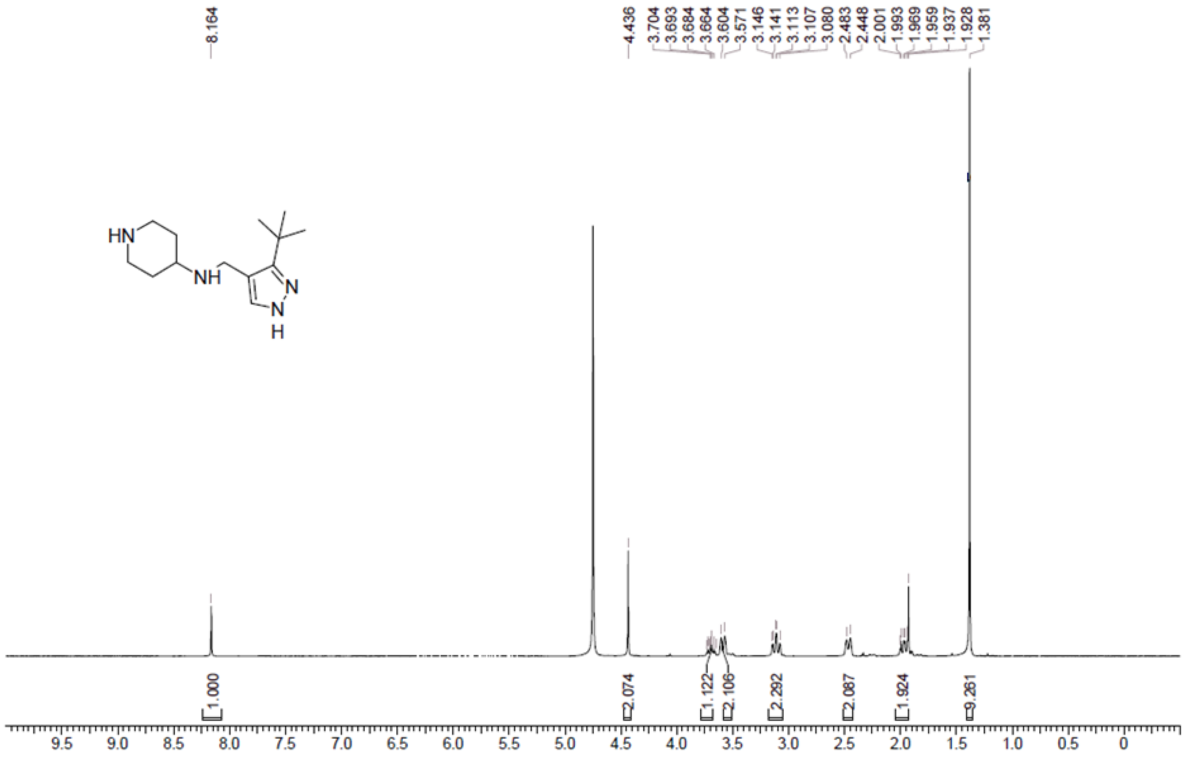

### [1]H NMR spectra of intermediate R2 in MeOD

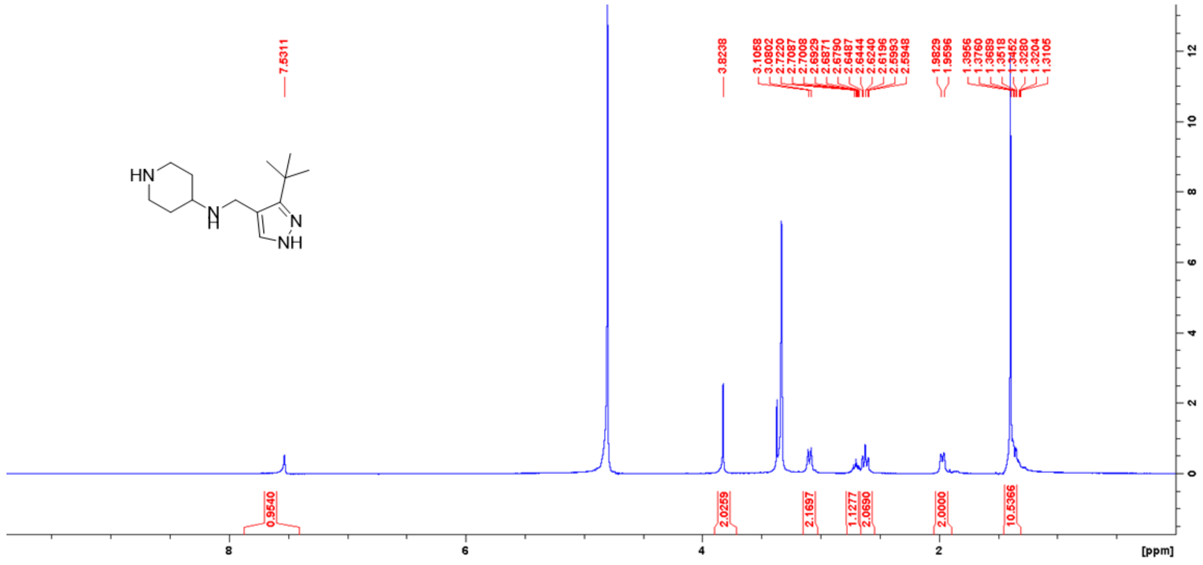

**[1]H and [13]C NMR spectra of compound 1 in MeOD**

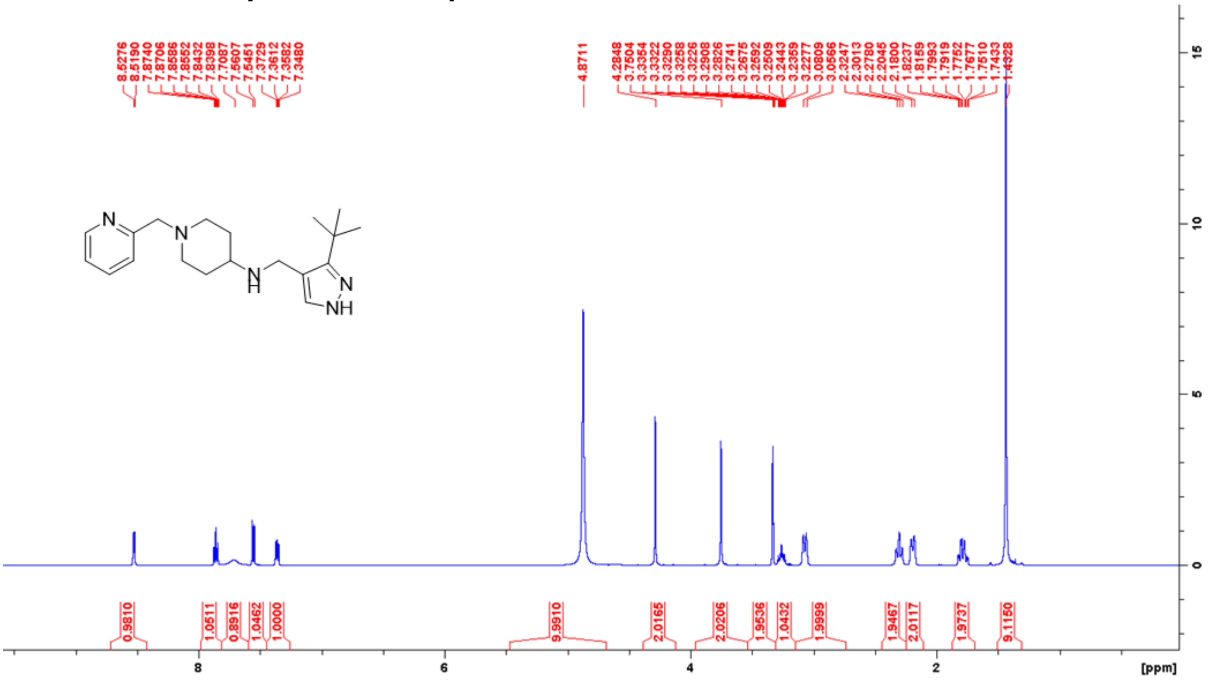

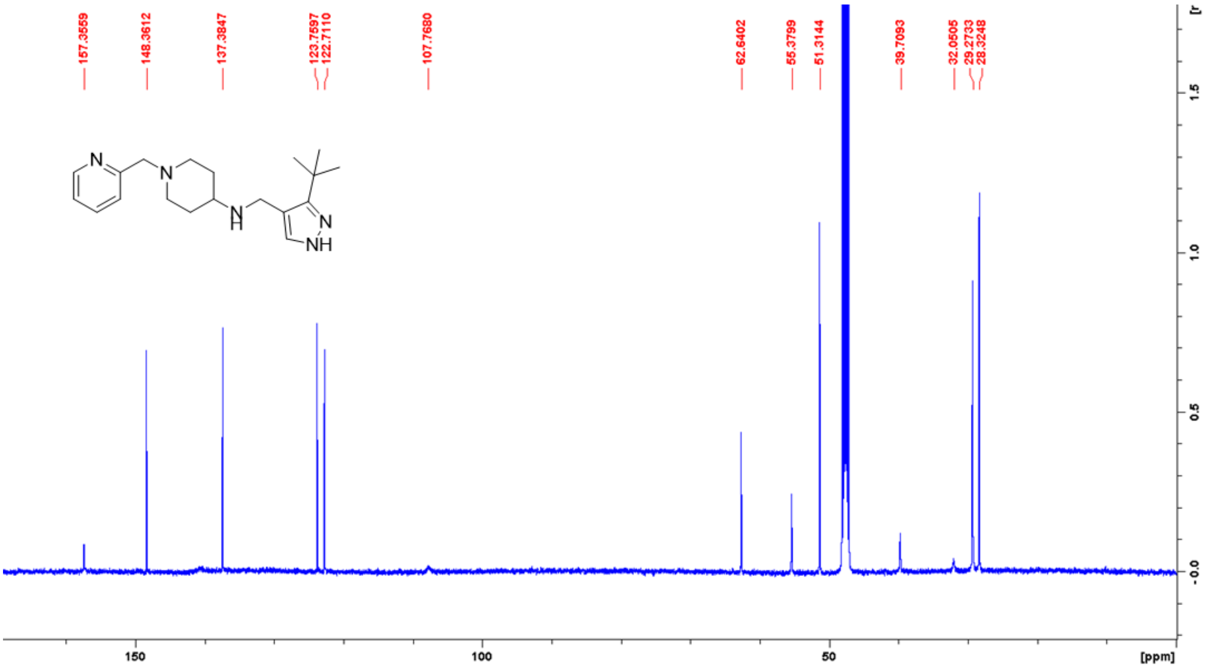

**$^1H$ and $^{13}C$ NMR spectra of compound 2 in MeOD**

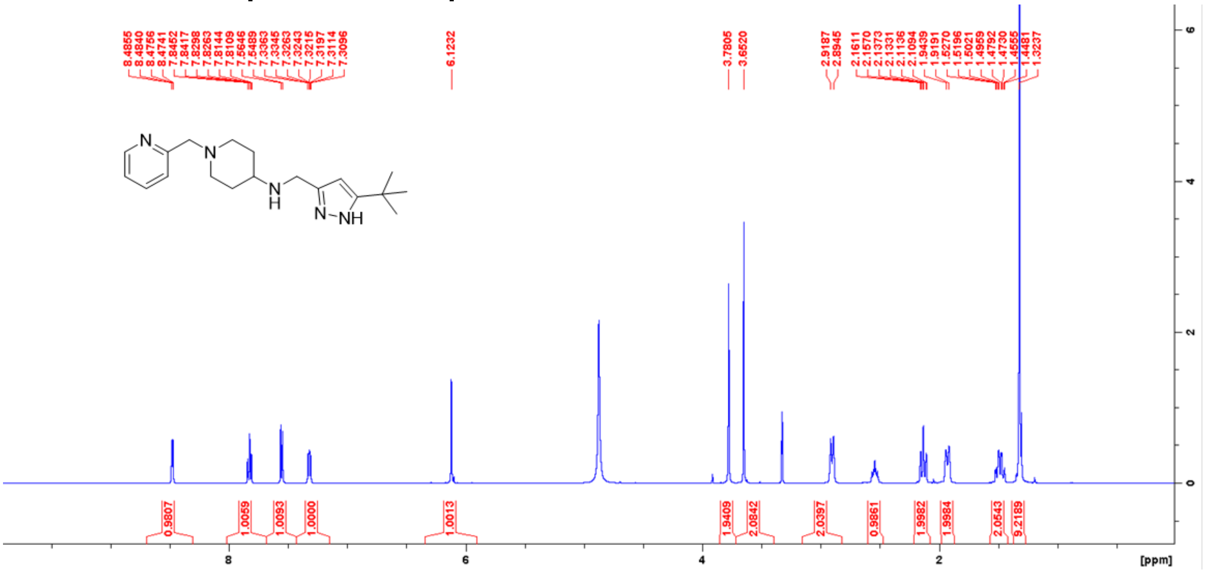

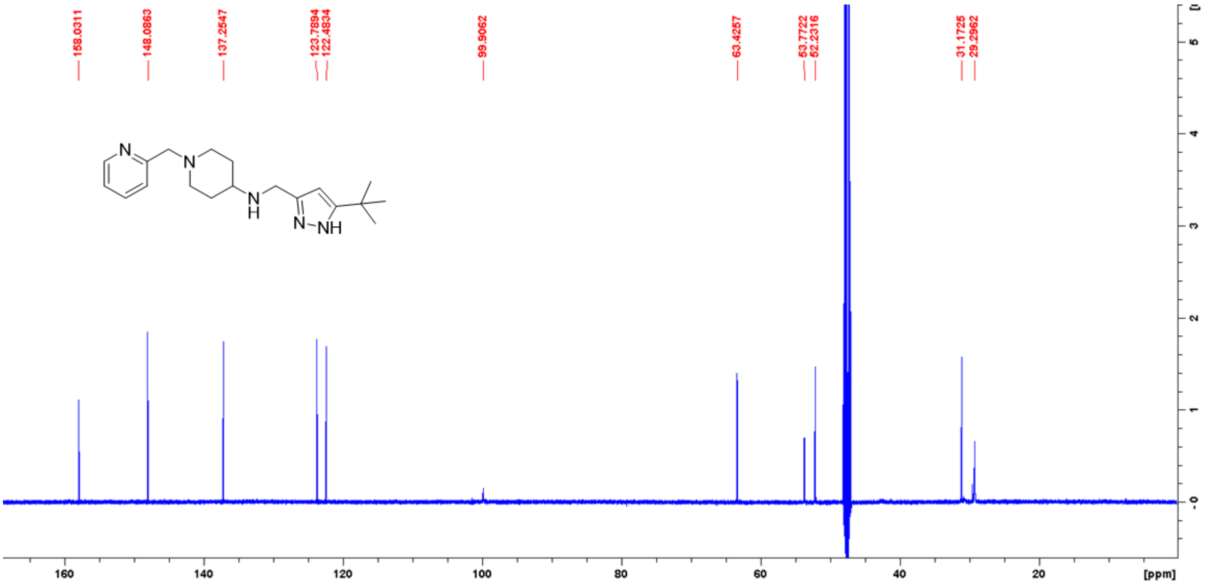

### $^{1}H$ and $^{13}C$ NMR spectra of compound 3 in MeOD

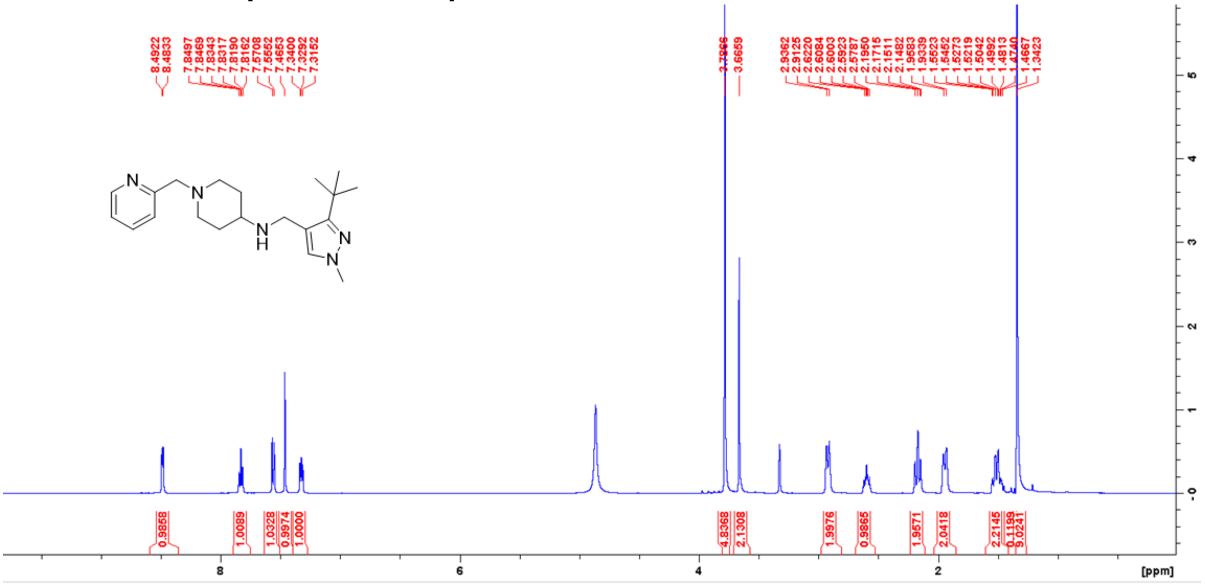


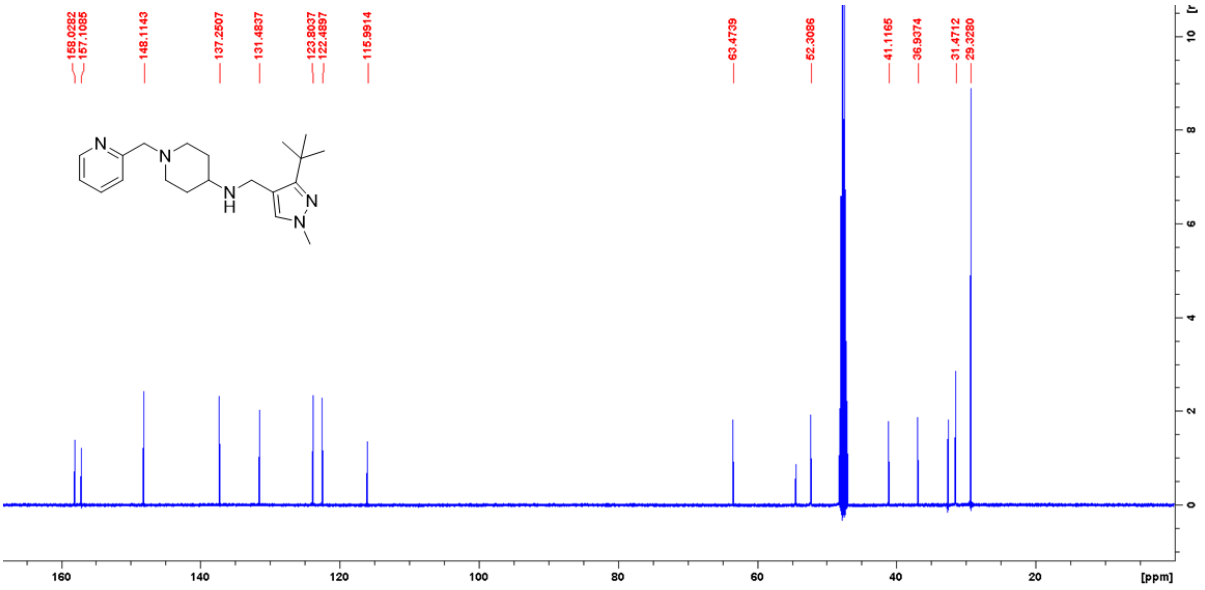

**$^{1}H$ and $^{13}C$ NMR spectra of compound 4 in MeOD**

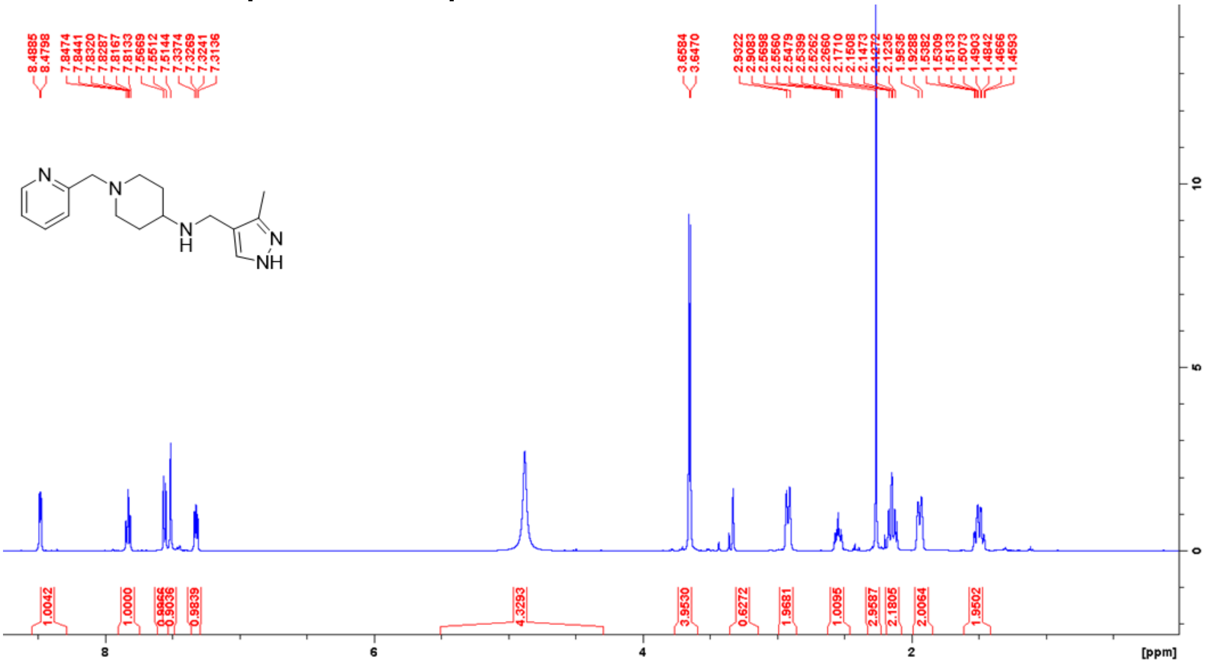


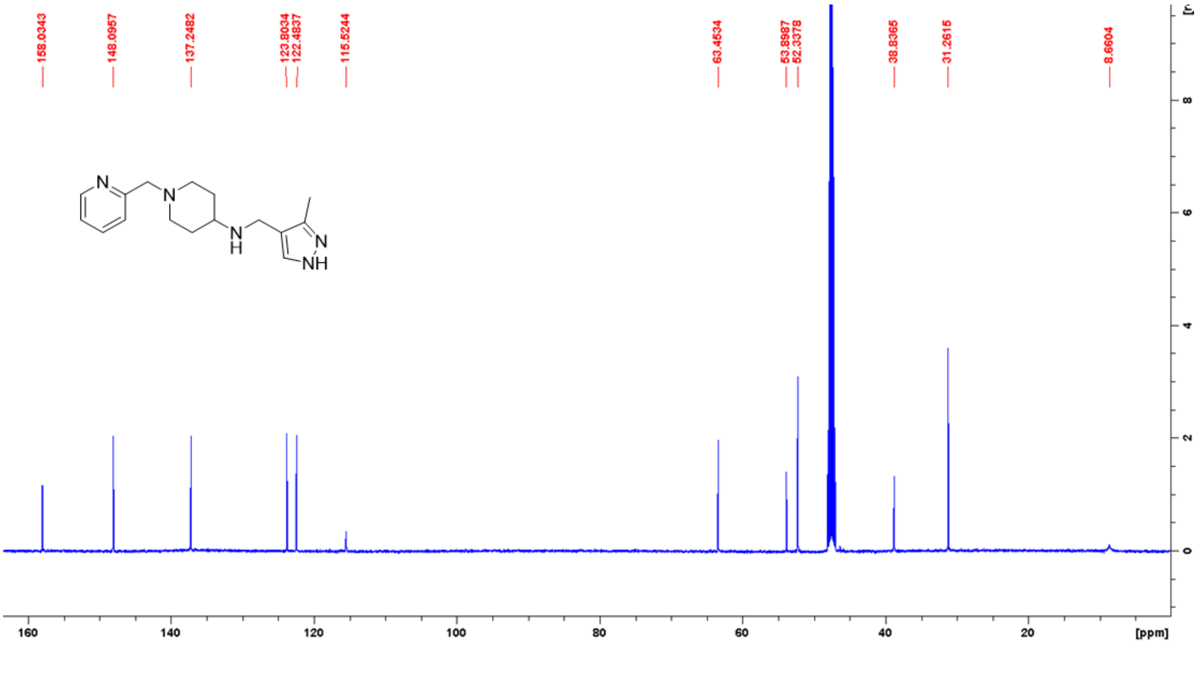

### $^{1}H$ and $^{13}C$ NMR spectra of compound 5 in MeOD

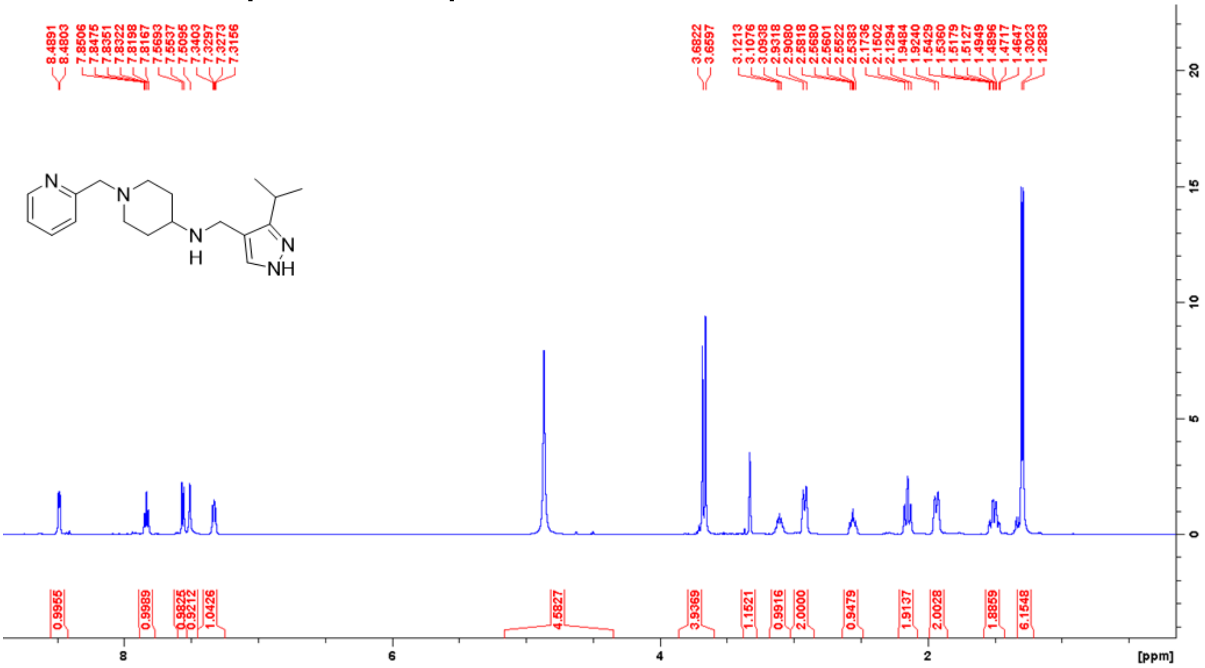


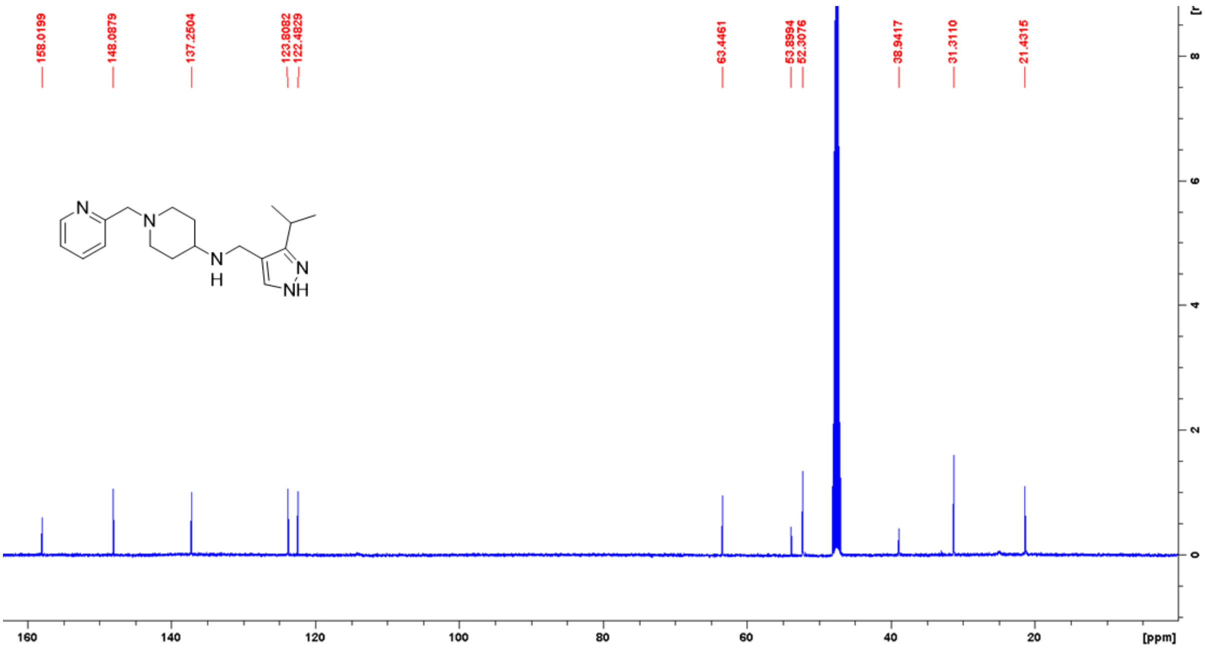

**[1]H and [13]C NMR spectra of compound 6 in MeOD**

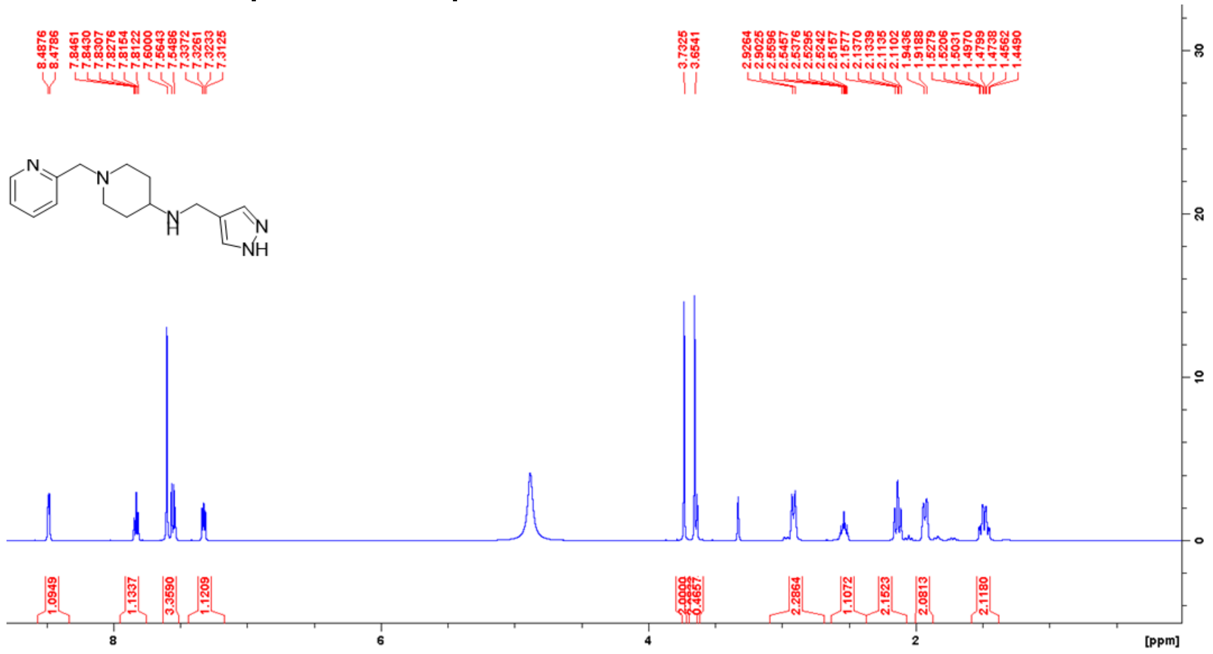

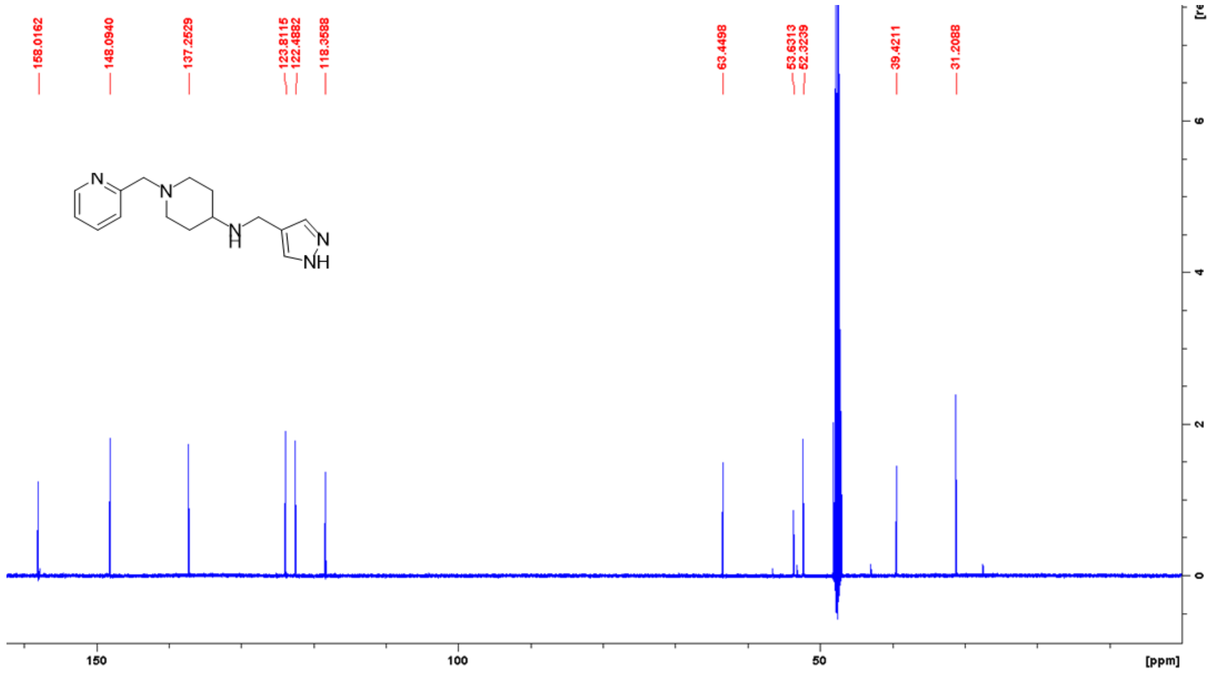

### [1]H and [13]C NMR spectra of compound 7 in MeOD

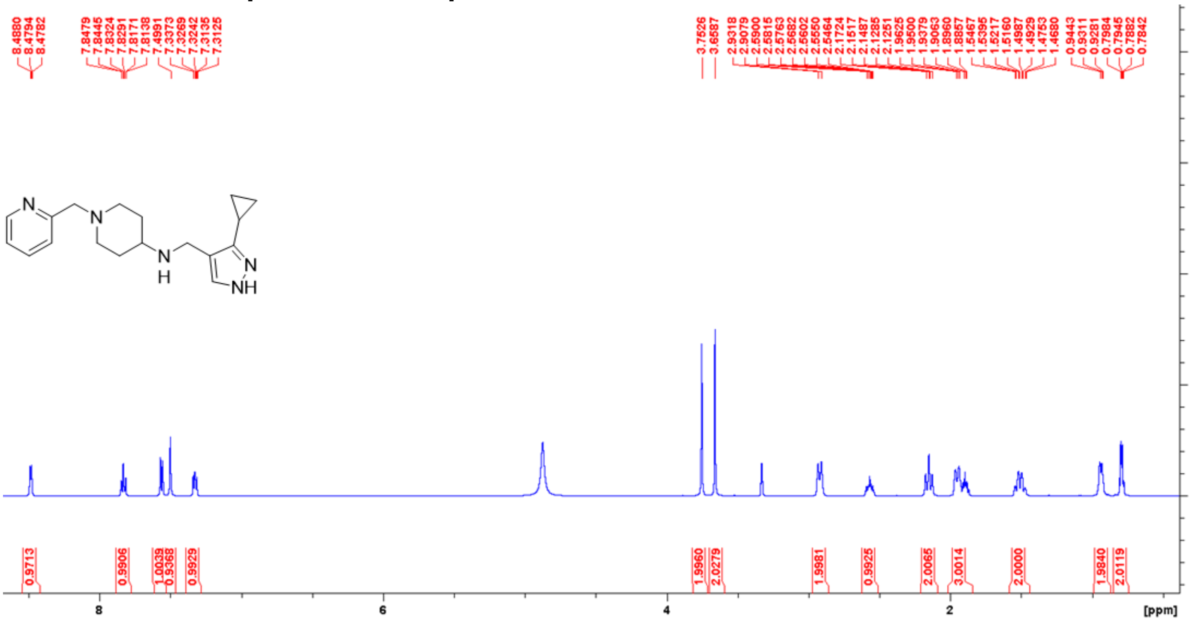

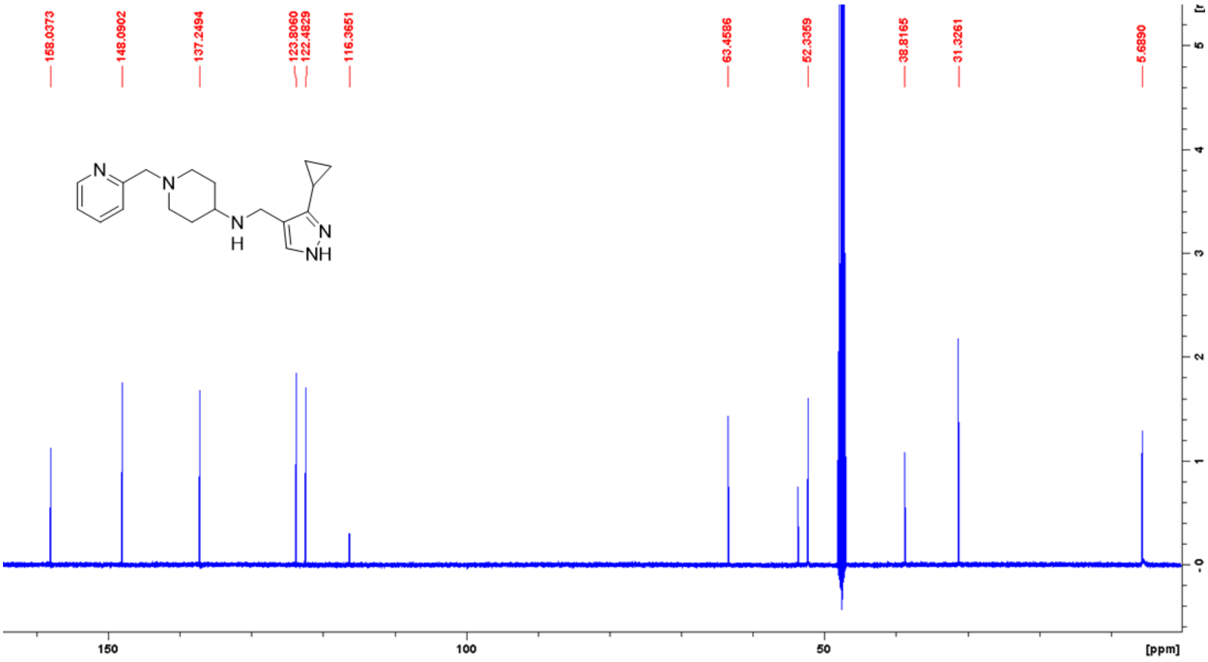

**[1]H and [13]C NMR spectra of compound 8 in MeOD**

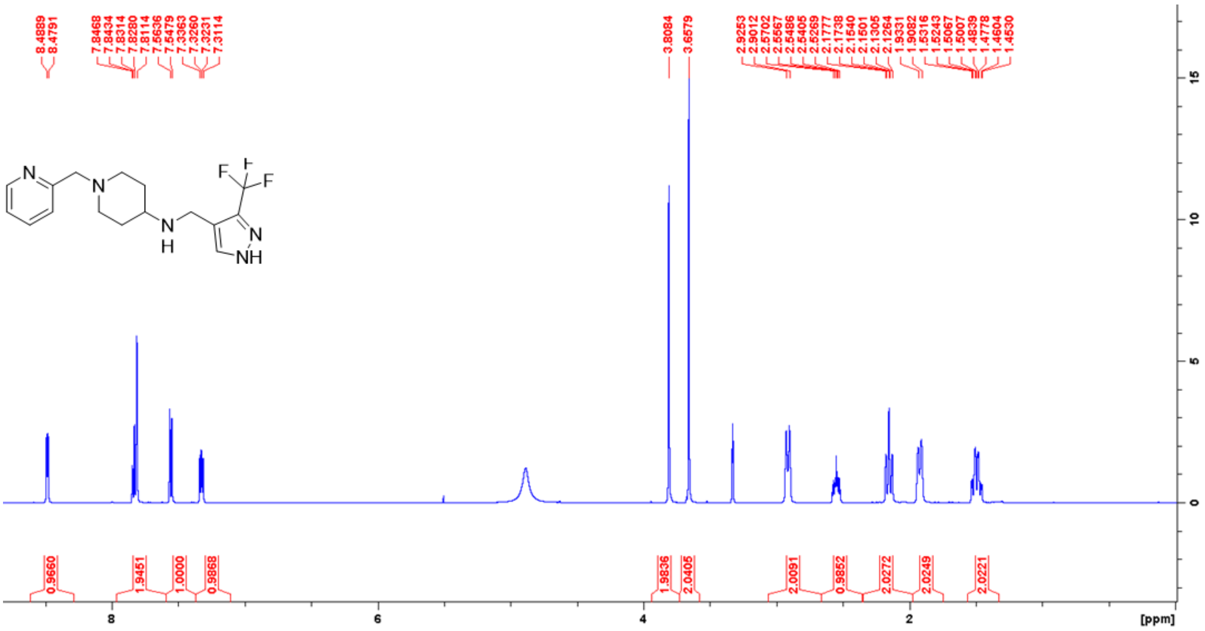


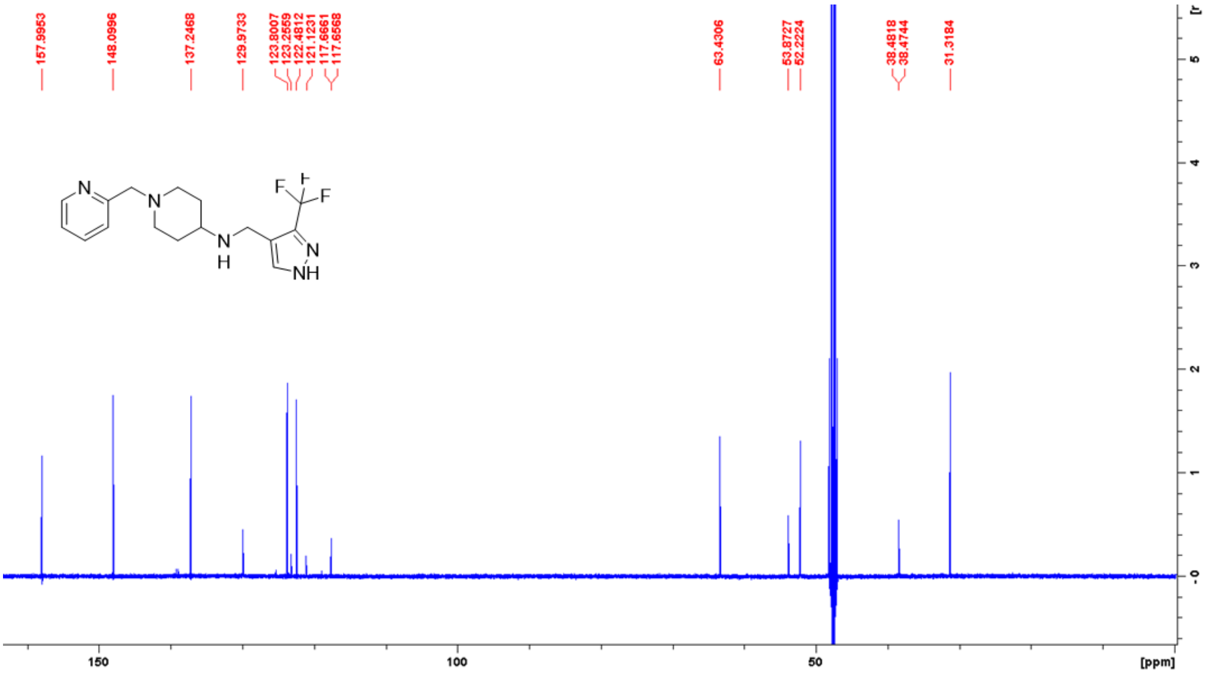

### $^{1}H$ and $^{13}C$ NMR spectra of compound 9 in MeOD

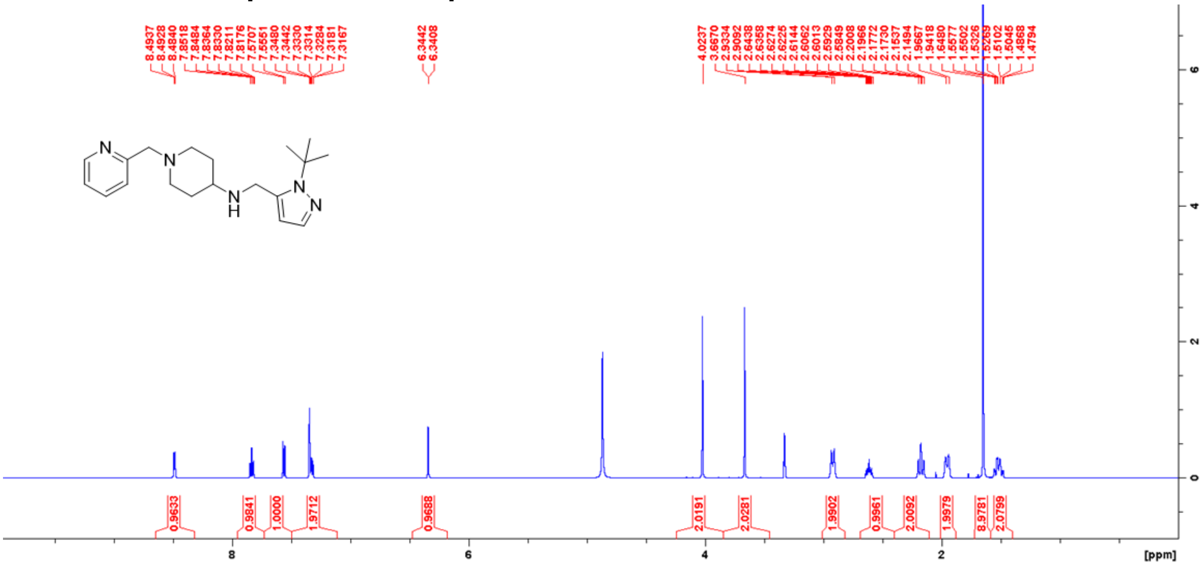

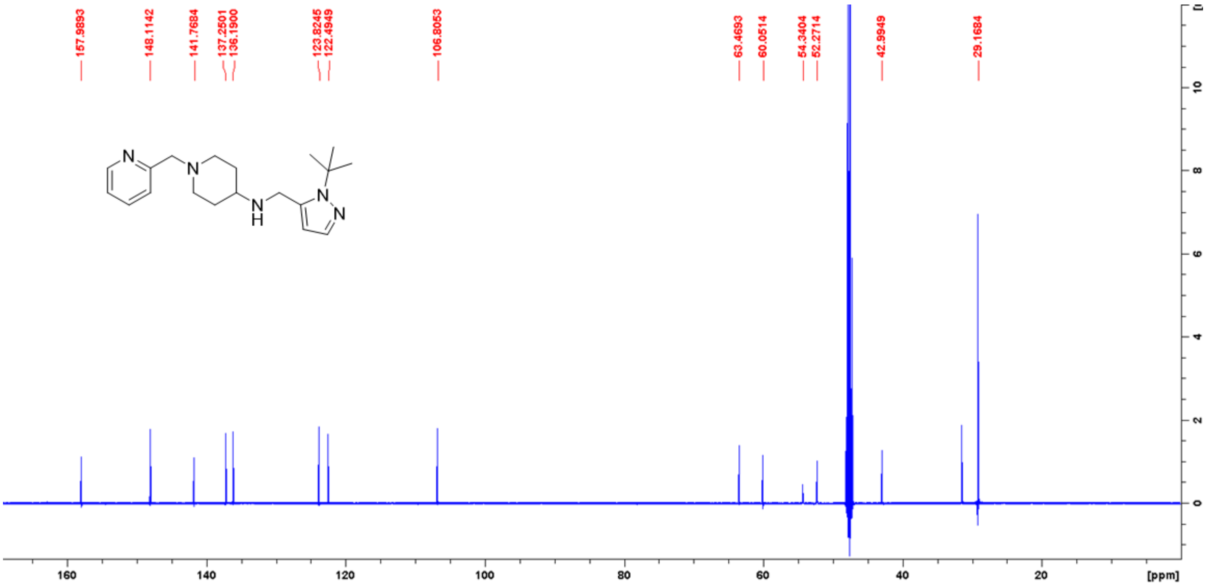

### [1]H and [13]C NMR spectra of compound 10 in MeOD

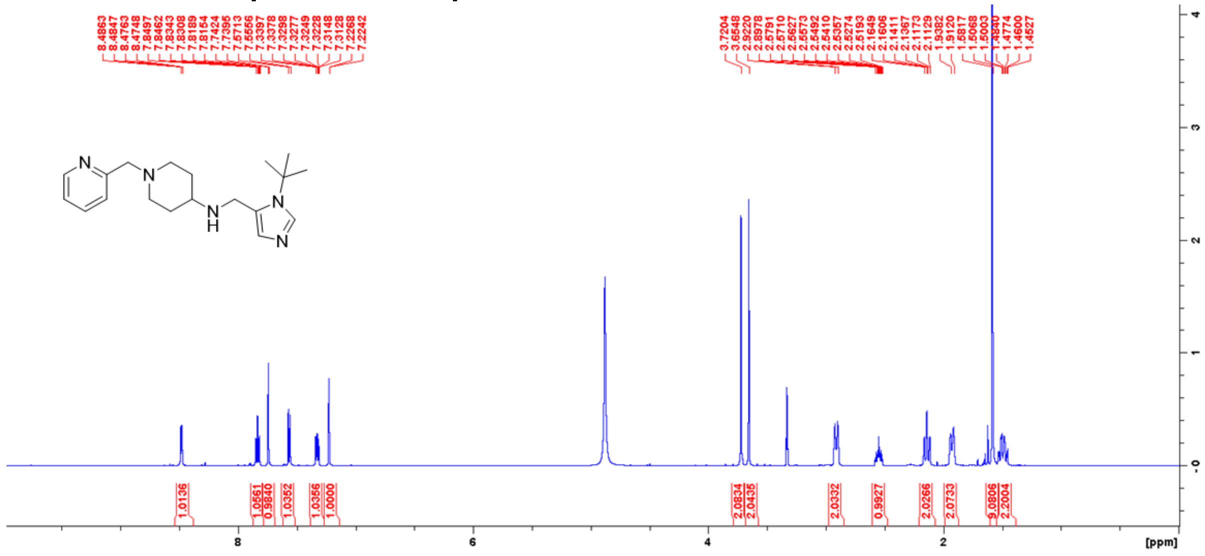


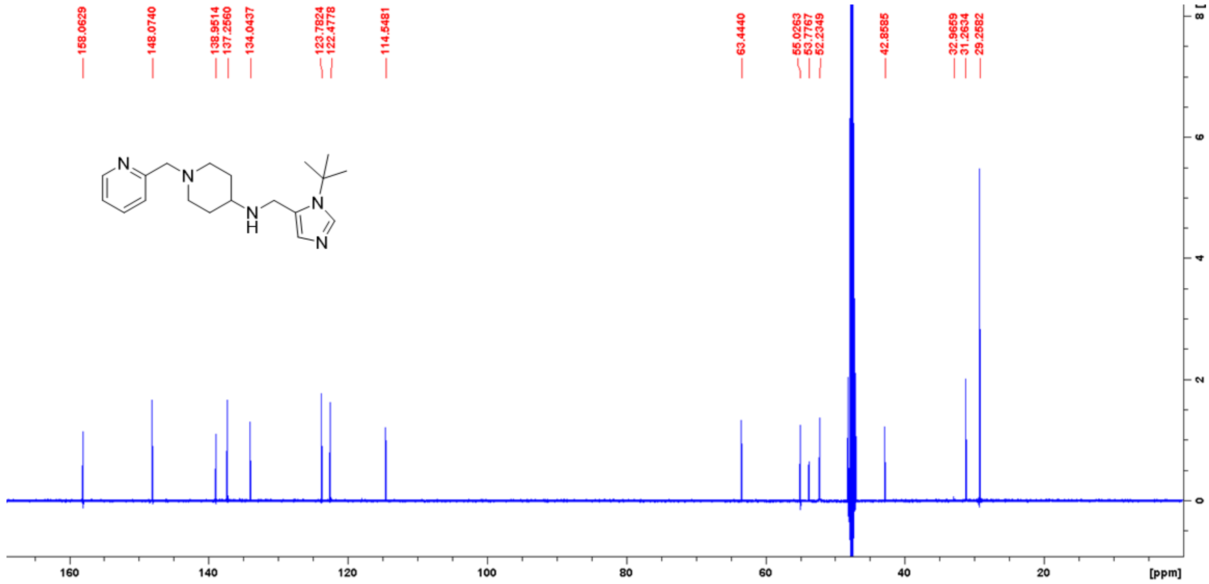

**[1]H and [13]C NMR spectra of compound 11 in MeOD**

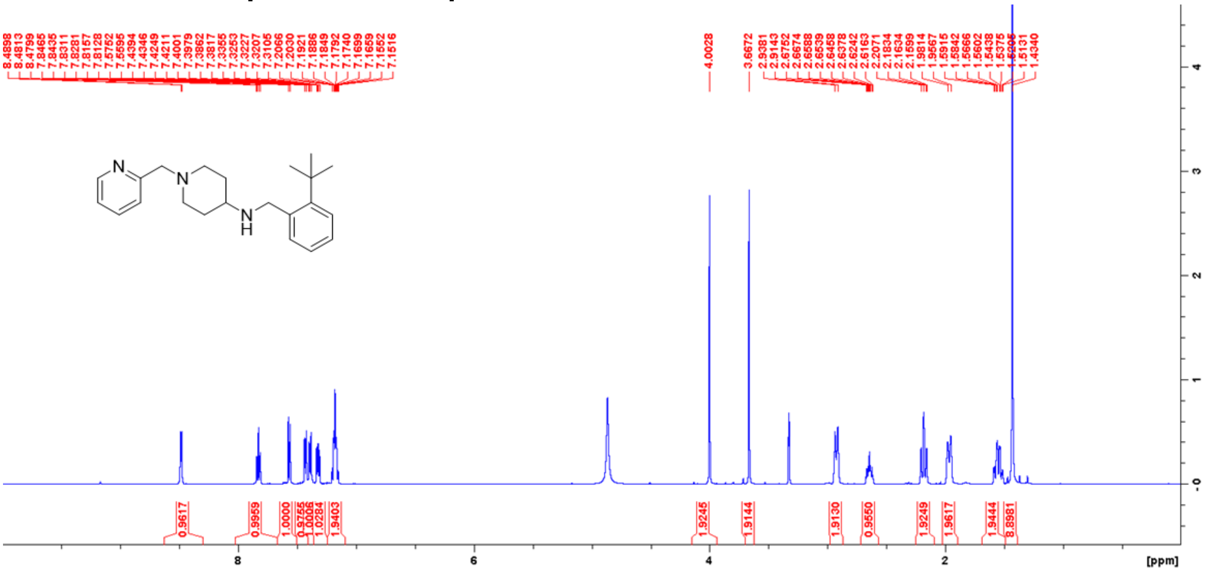

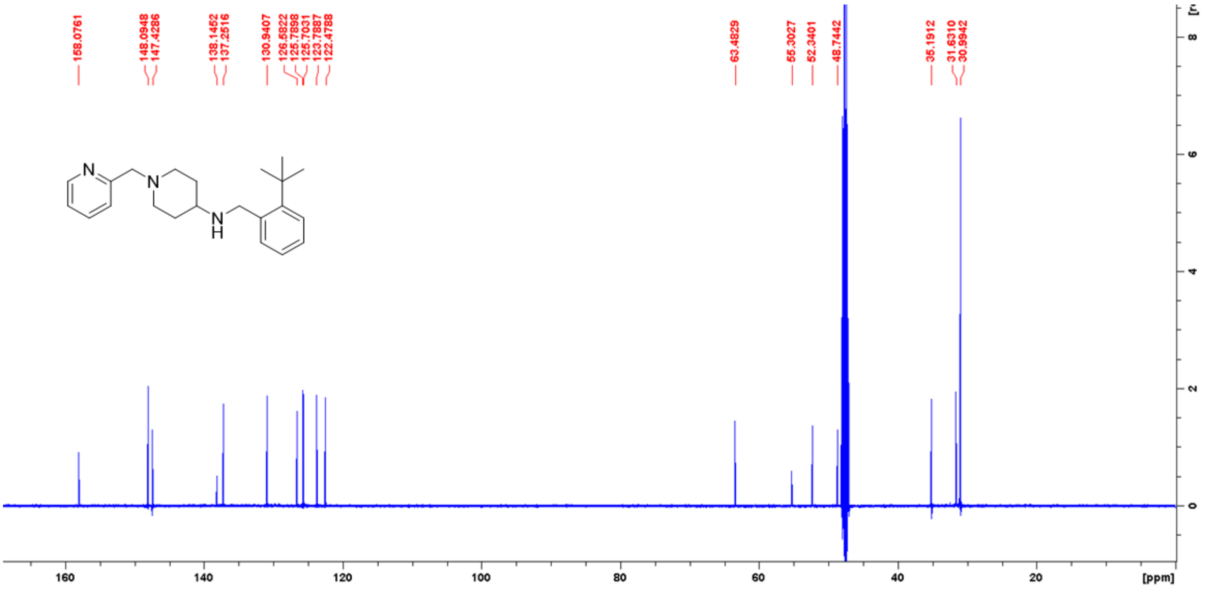

**$^{1}H$ and $^{13}C$ NMR spectra of compound 12 in MeOD**

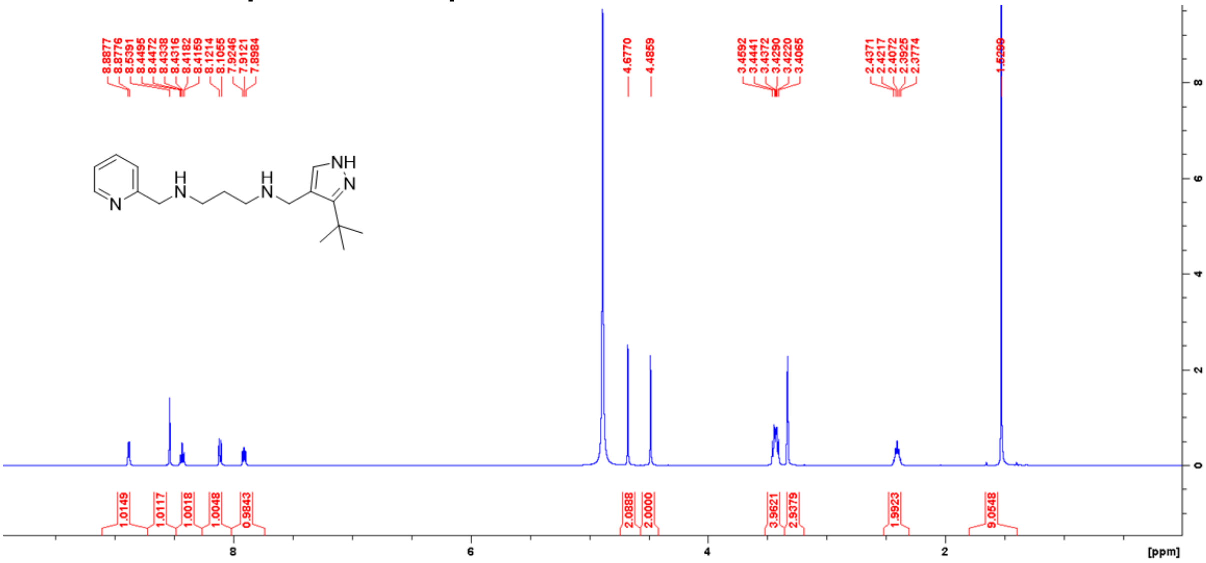

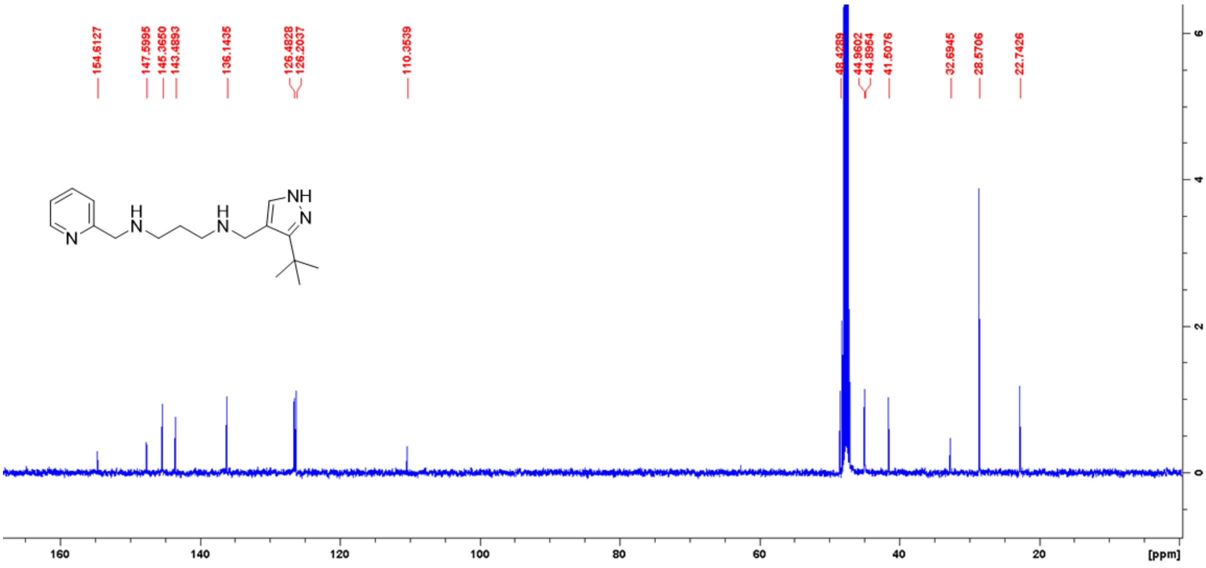

#### ¹H and ¹³C NMR spectra of compound 13 in MeOD

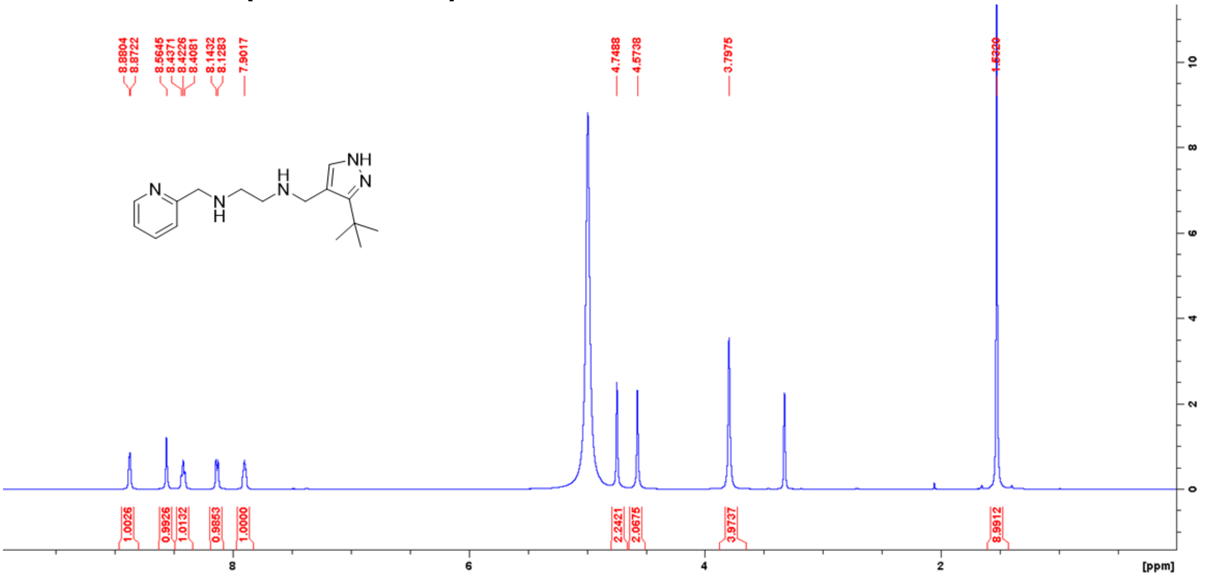

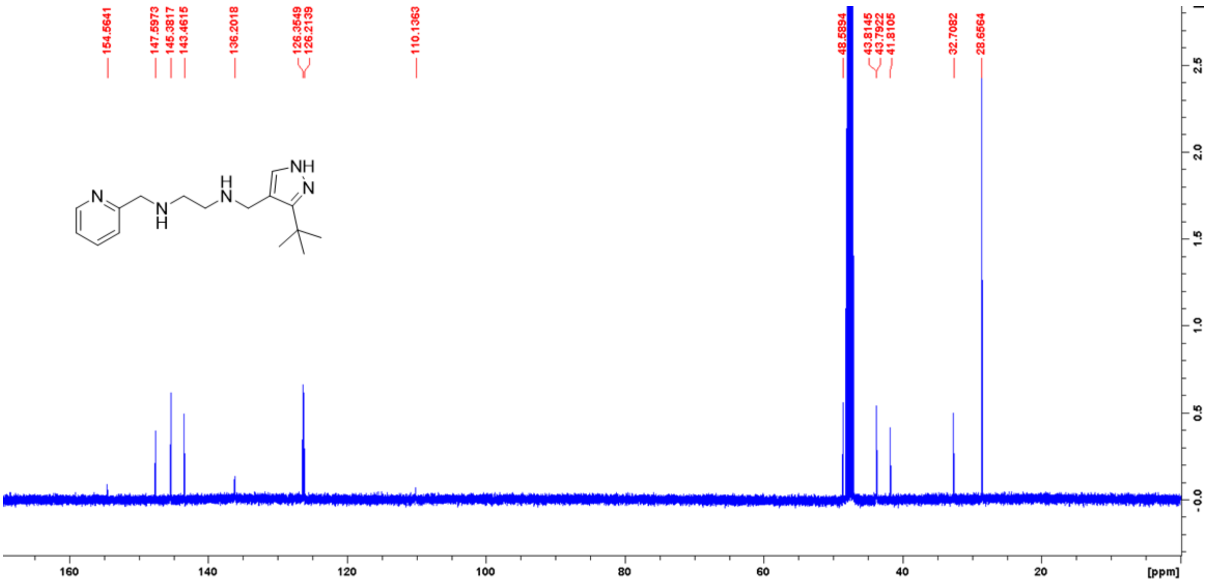

**$^{1}$H and $^{13}$C NMR spectra of compound 14 in MeOD**

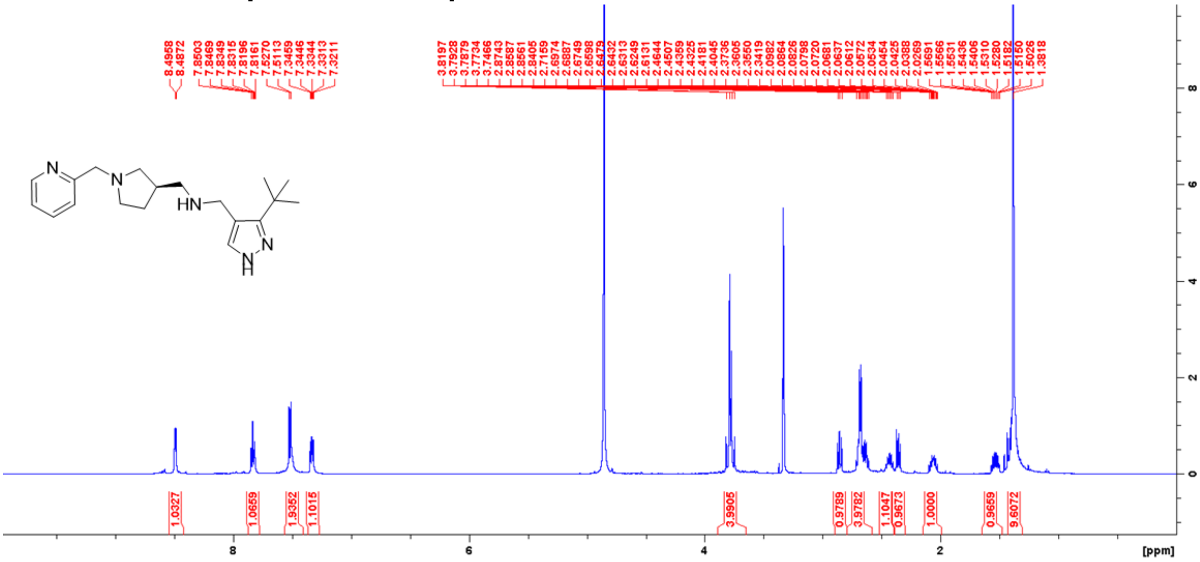

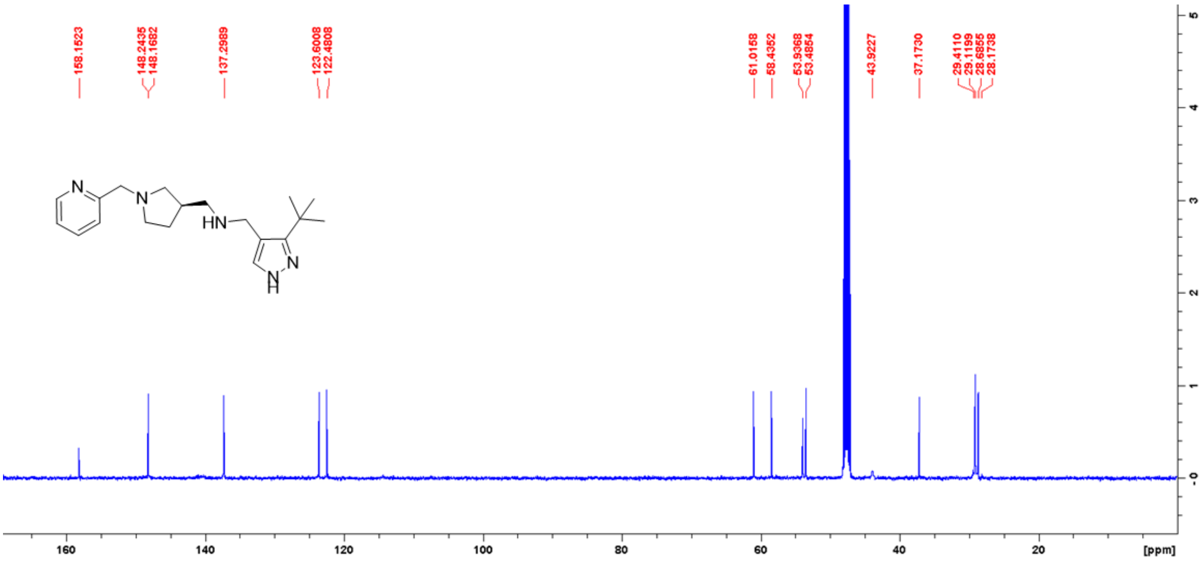

[1]H and [13]C NMR spectra of compound 15 in MeOD

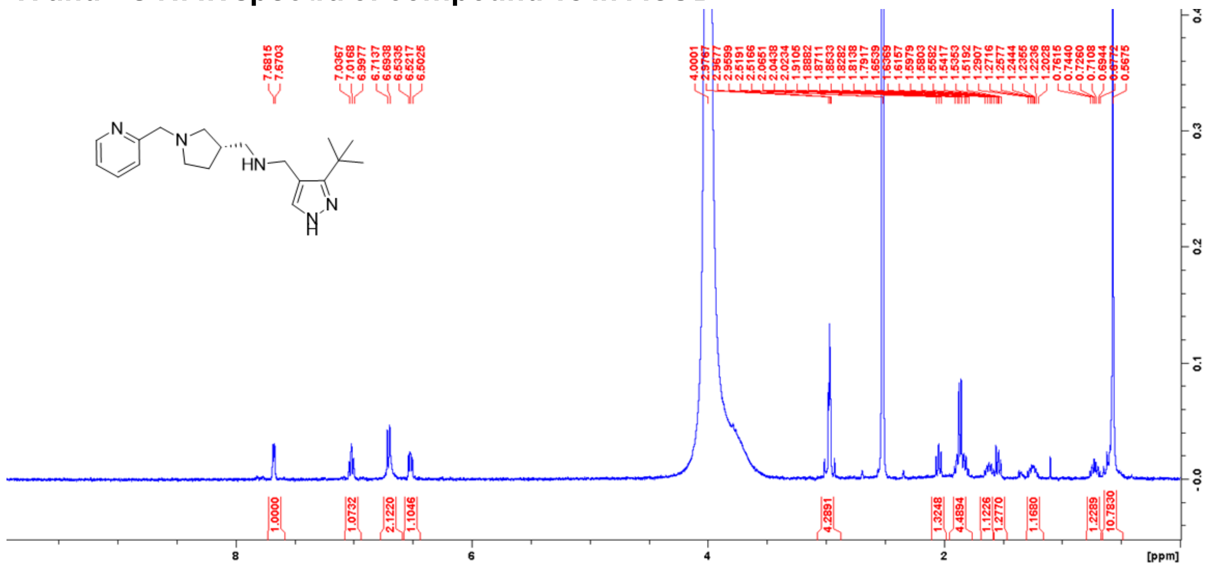

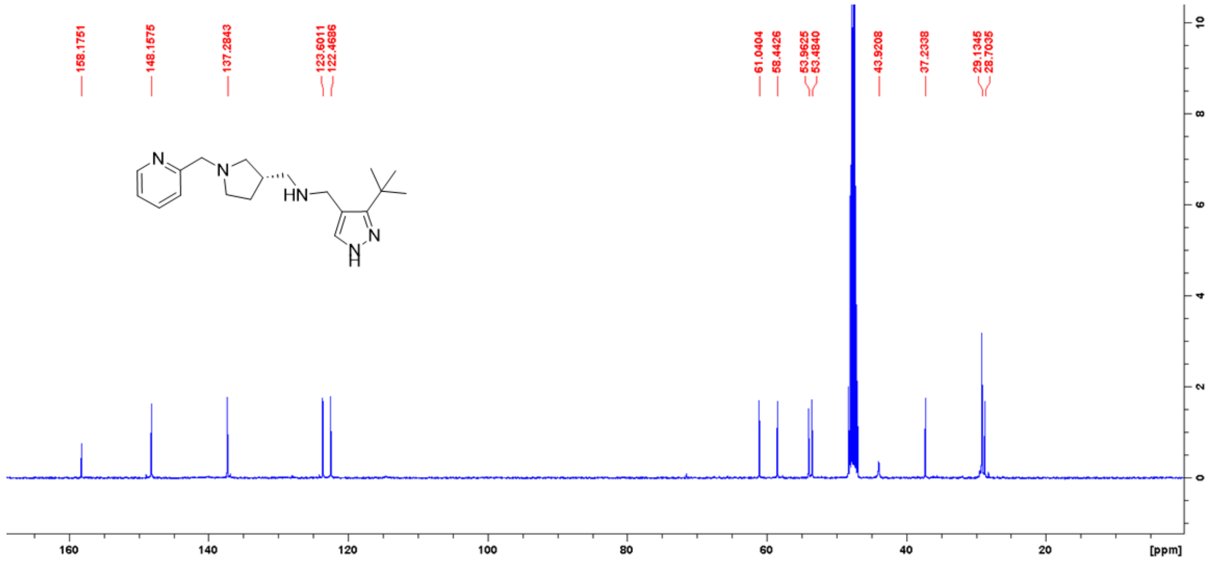

### $^{1}H$ and $^{13}C$ NMR spectra of compound 16 in MeOD

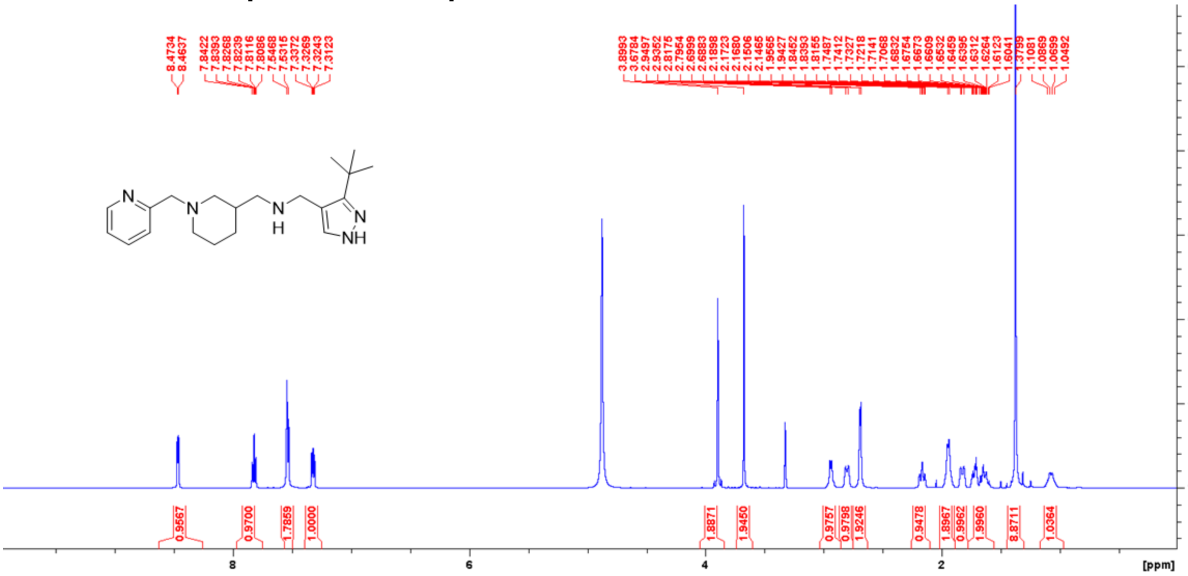


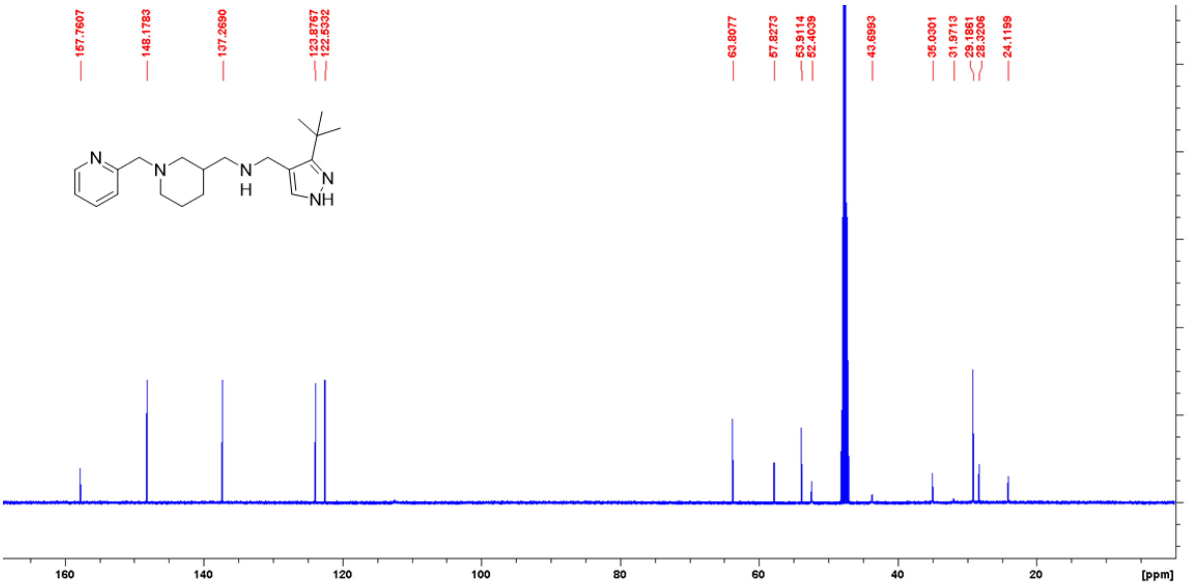

### $^{1}H$ and $^{13}C$ NMR spectra of compound 17 in MeOD

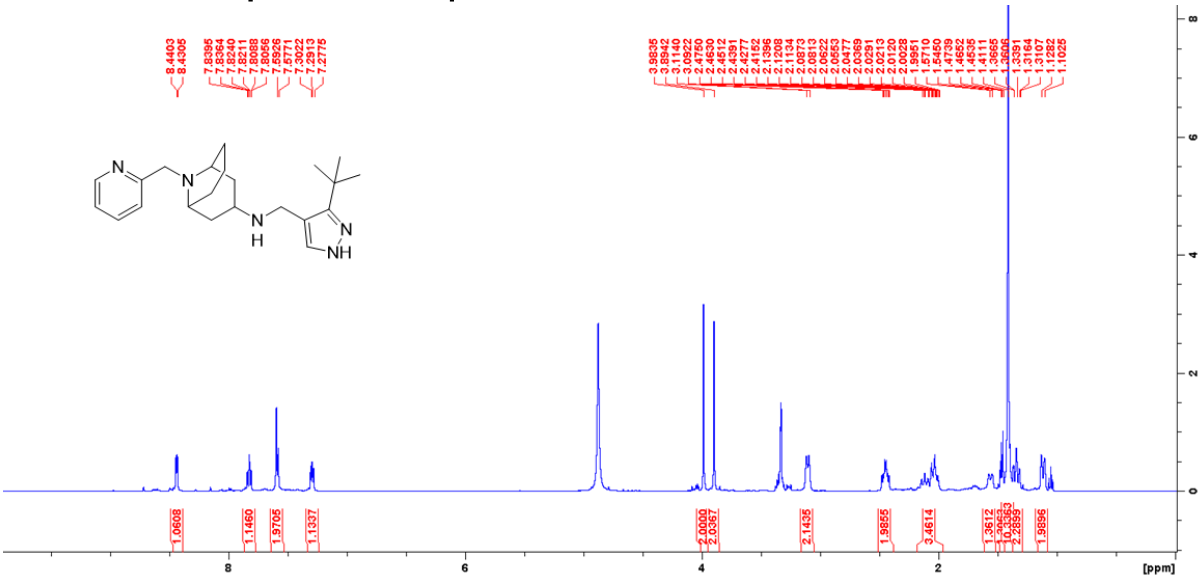

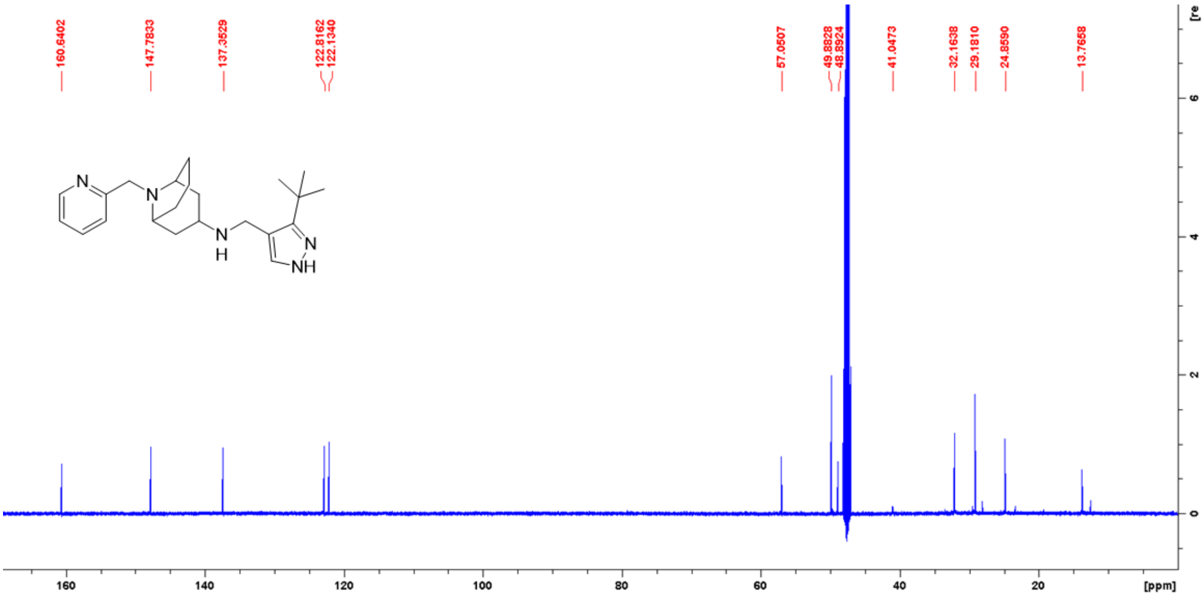

### $^{1}H$ and $^{13}C$ NMR spectra of compound 18 in MeOD

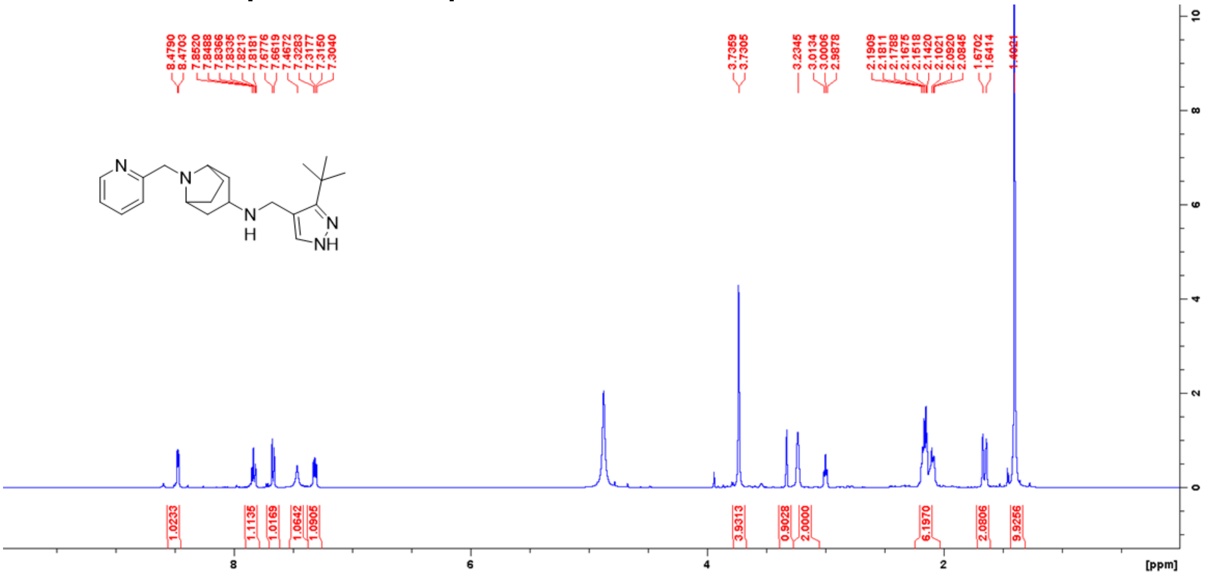

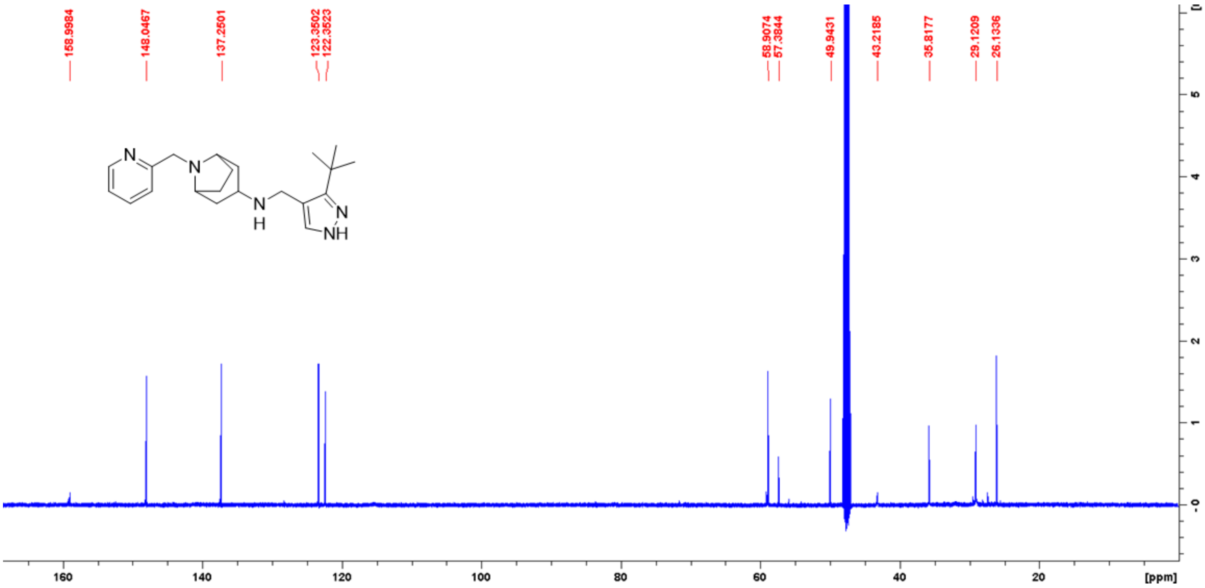

**$^1$H and $^{13}$C NMR spectra of compound 19 in MeOD**

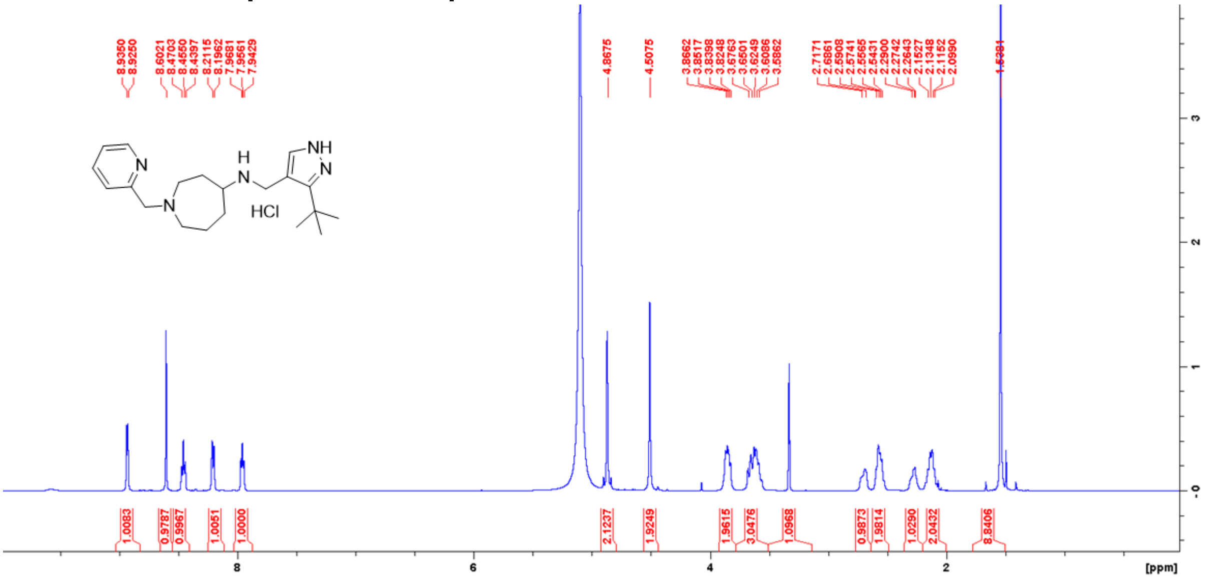


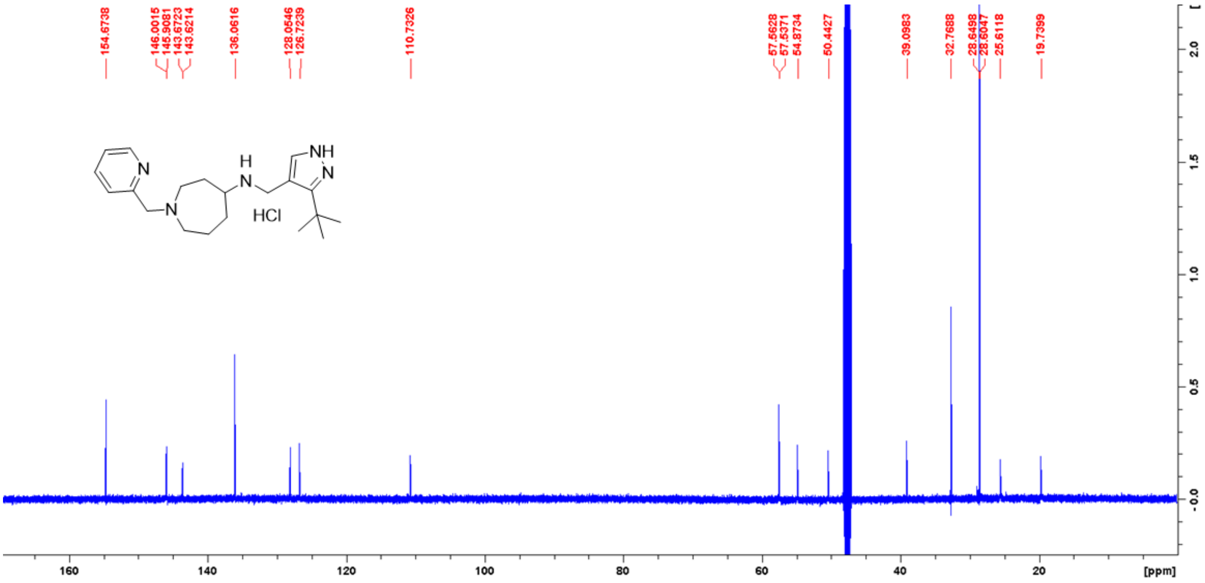

**[1]H NMR spectra of compound 20 in MeOD**

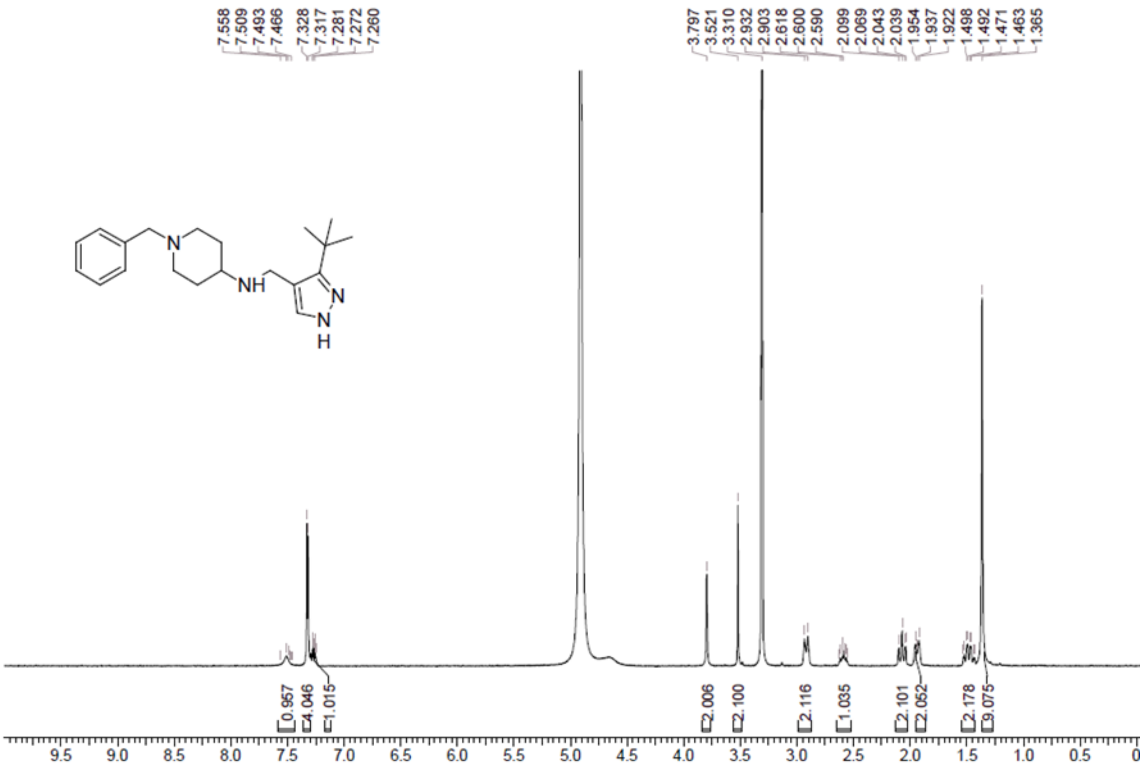

**[1]H and [13]C NMR spectra of compound 21 in MeOD**

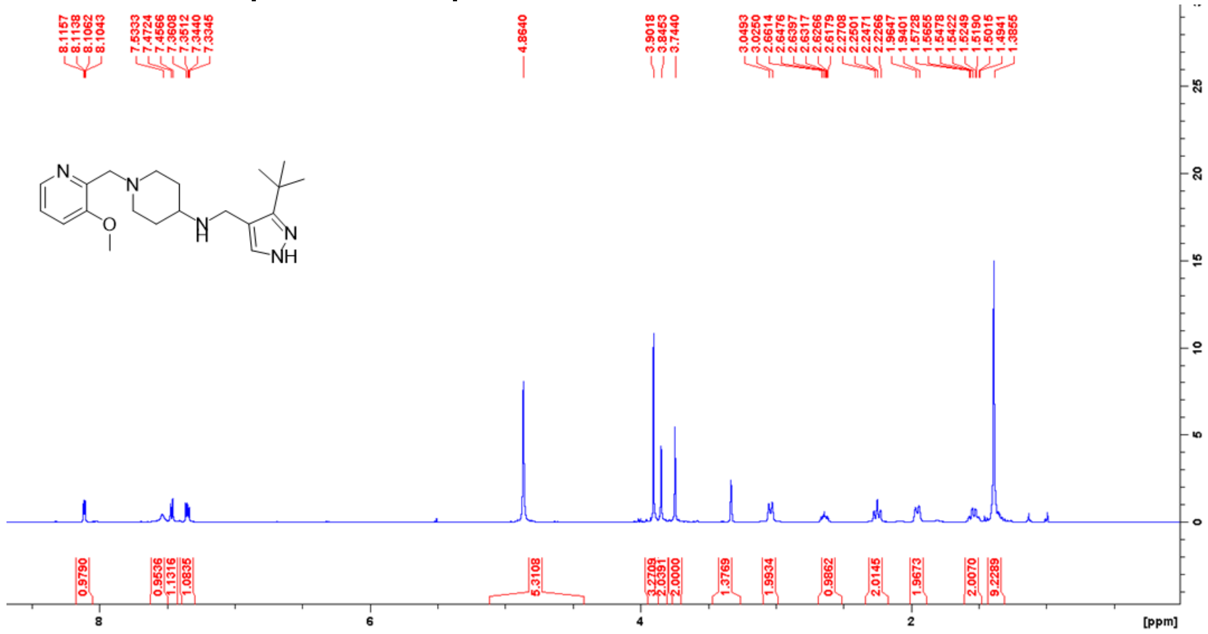

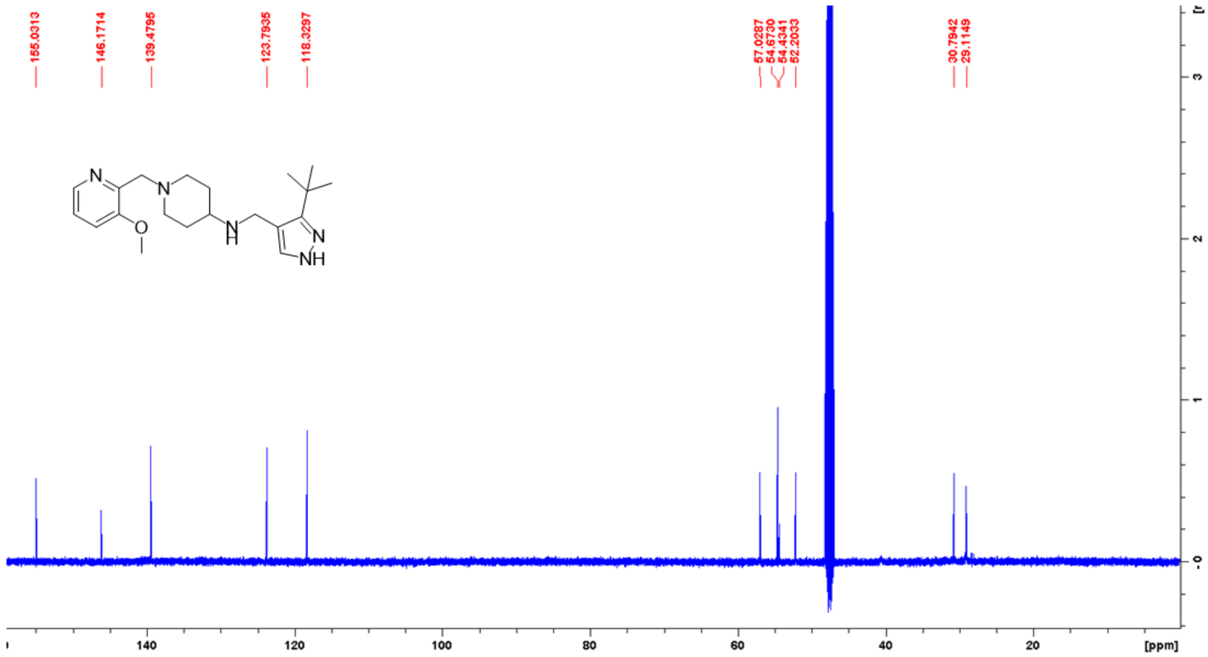

**[1]H NMR spectra of compound 22 in DMSO-d6**

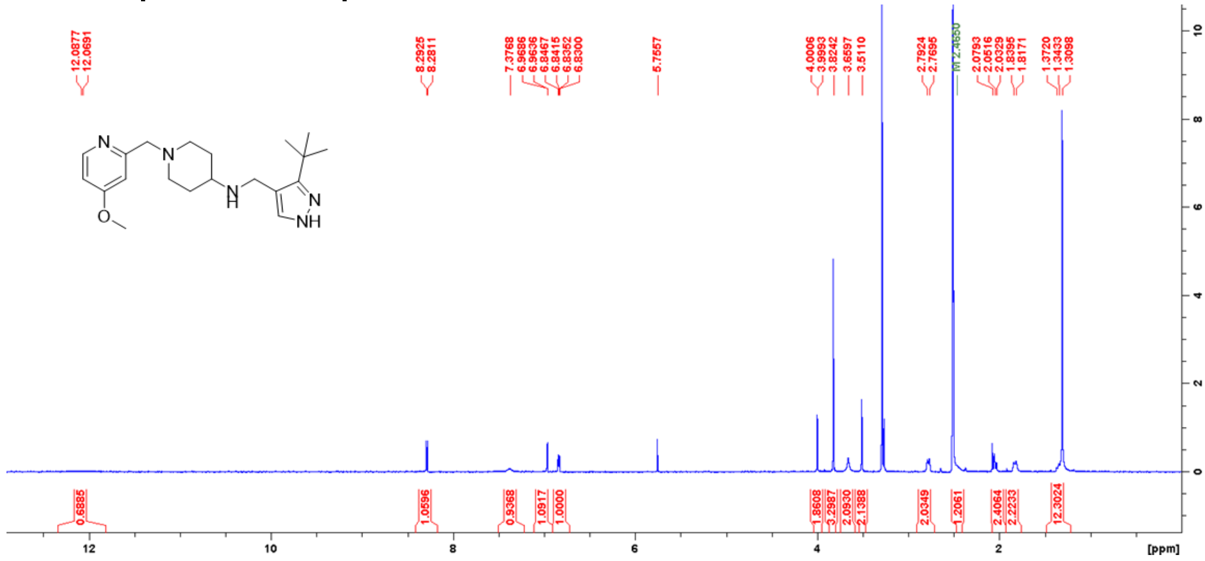

### [1]H and [13]C NMR spectra of compound 23 in MeOD

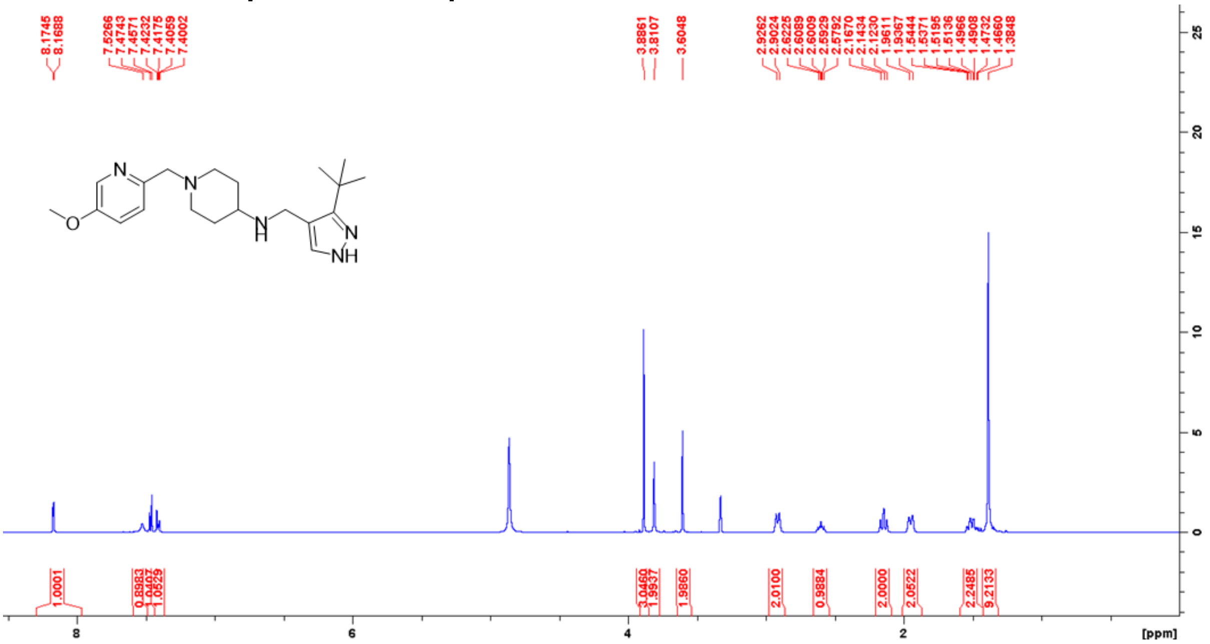

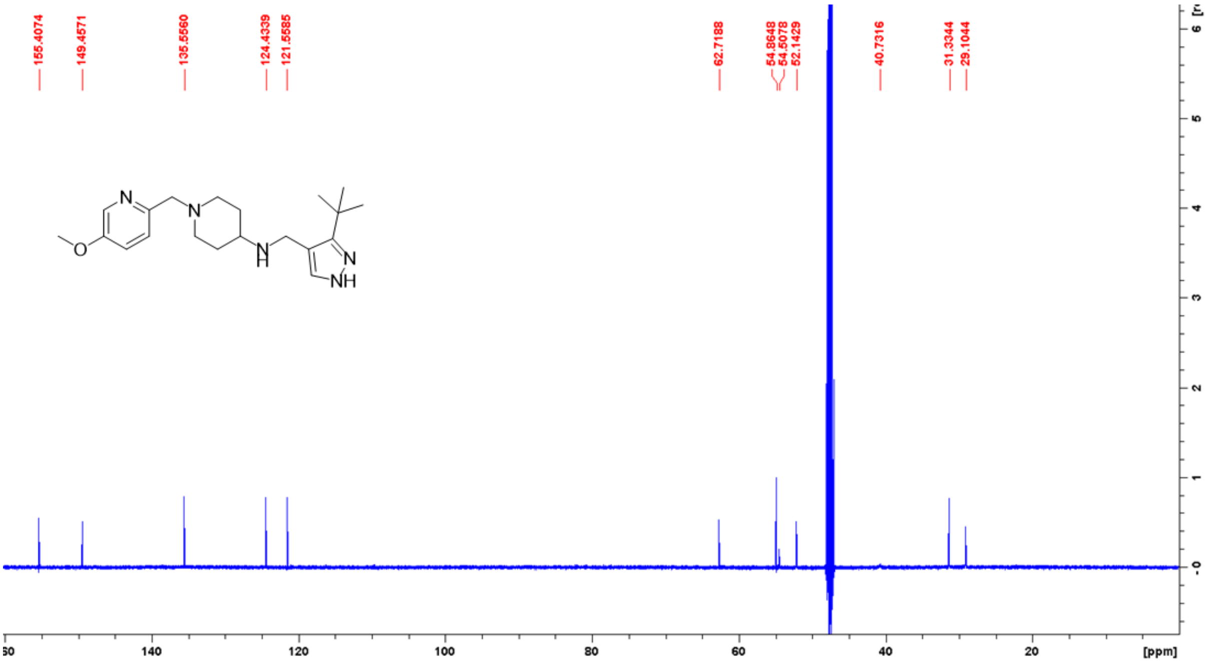

### [1]H and [13]C NMR spectra of compound 24 in MeOD

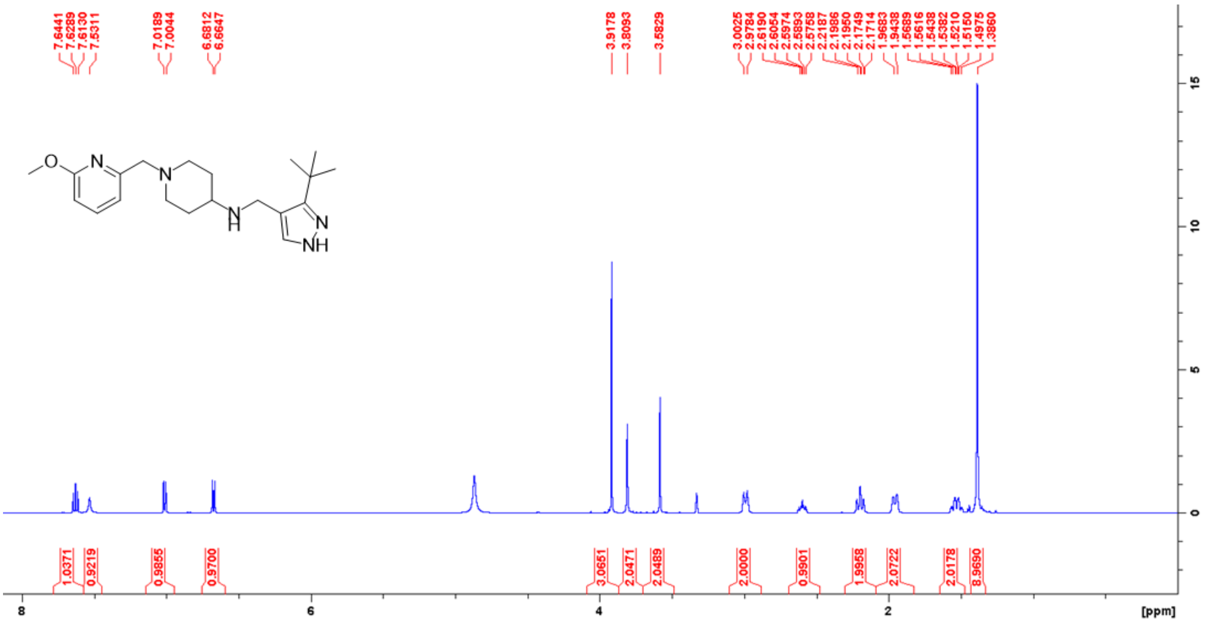


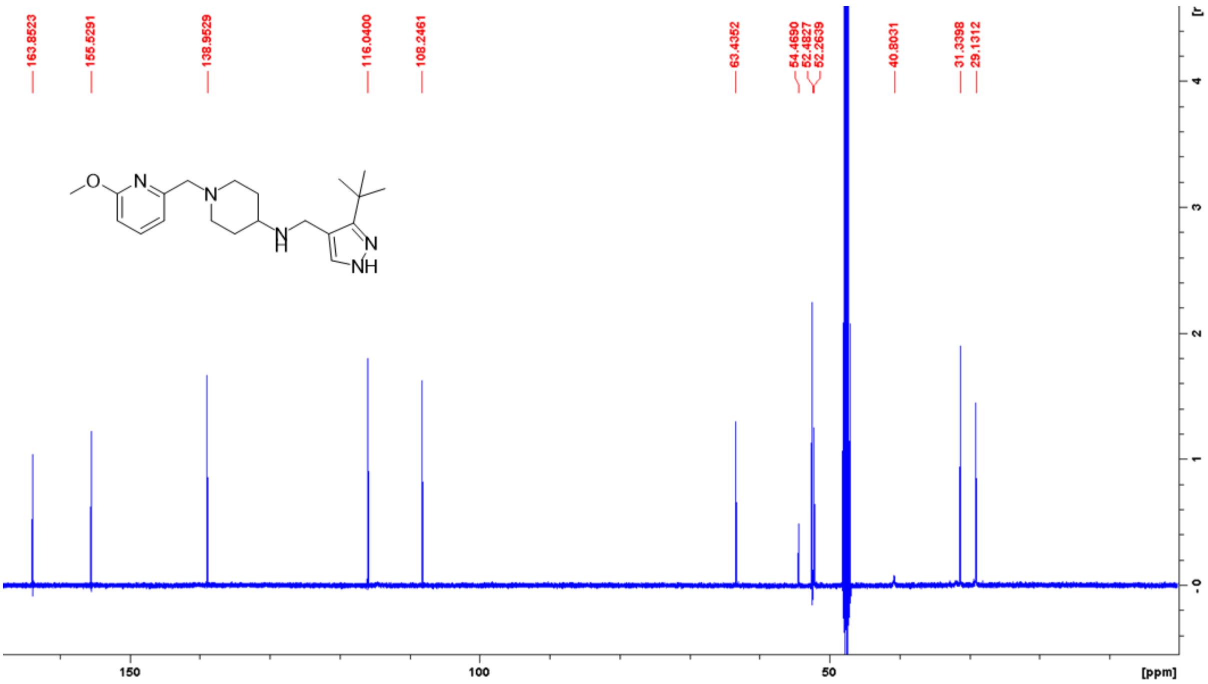

**[1]H and [13]C NMR spectra of compound 25 in MeOD**

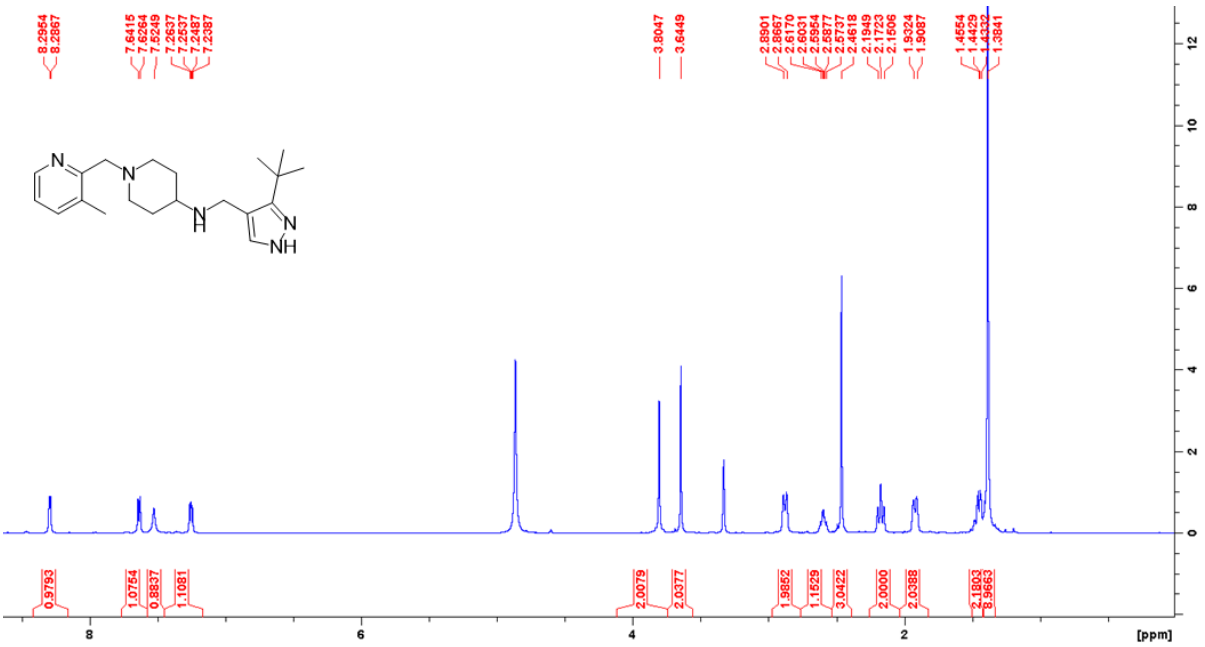

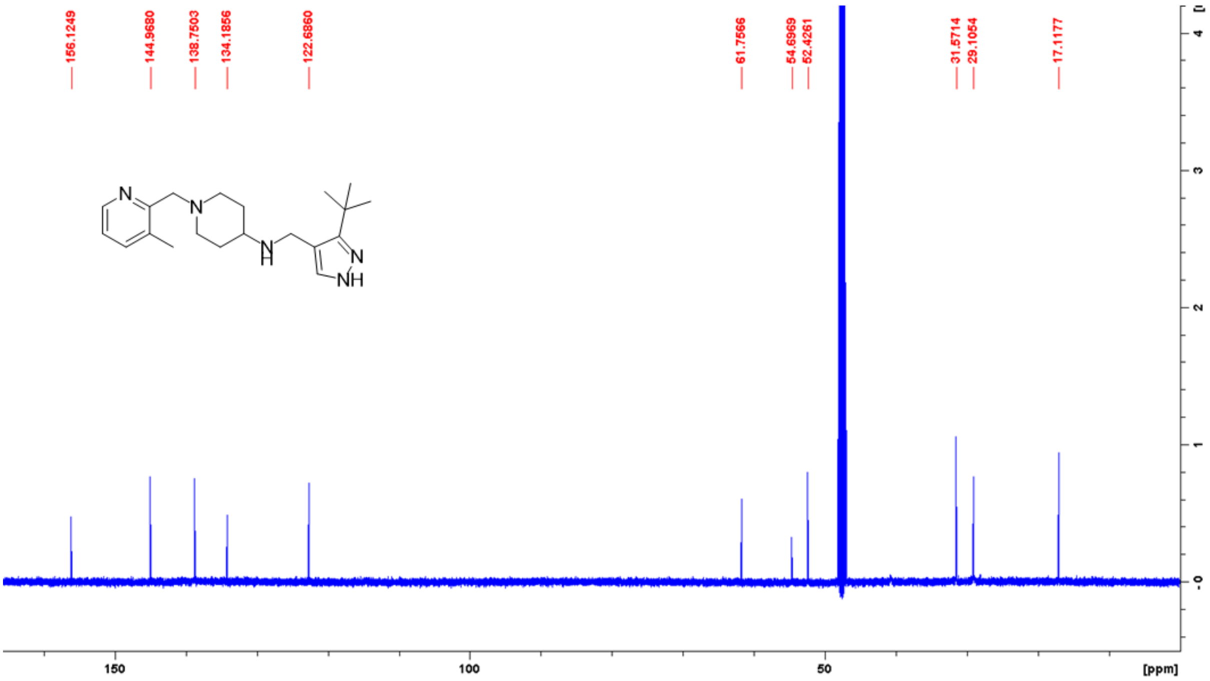

**¹H NMR spectra of compound 26 in DMSO-d6**

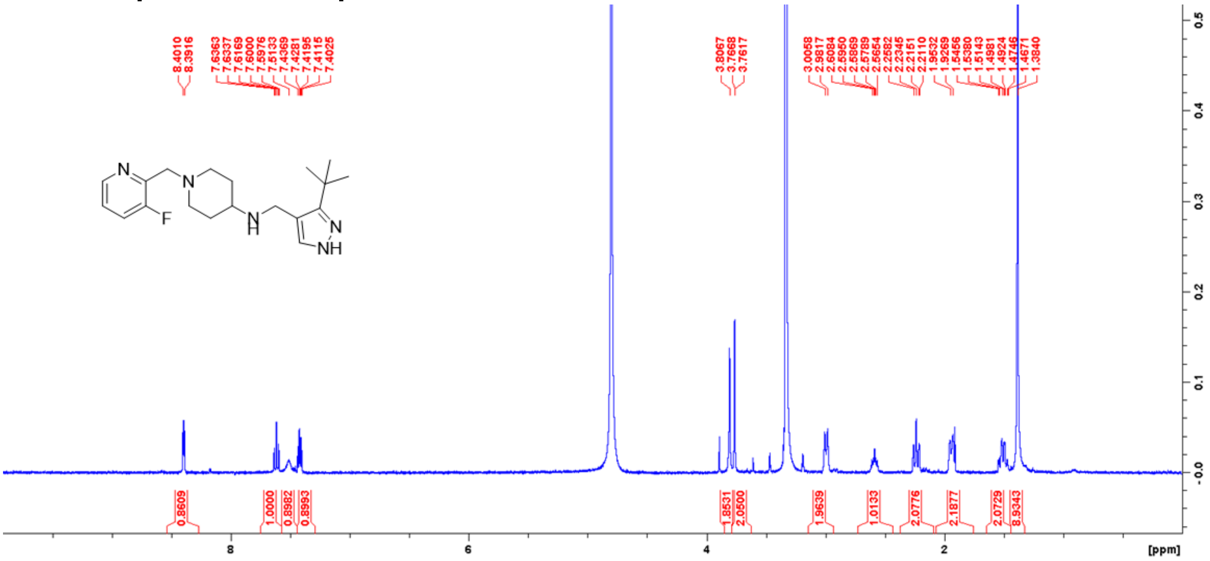

**[1]H and [13]C NMR spectra of compound 27 in MeOD**

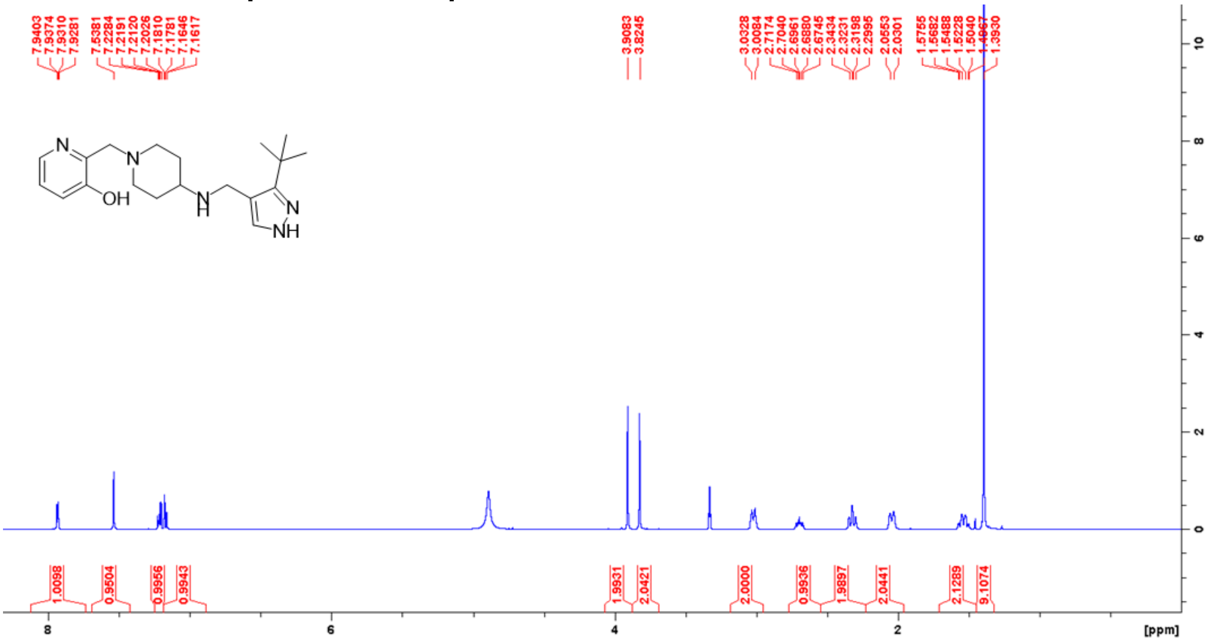


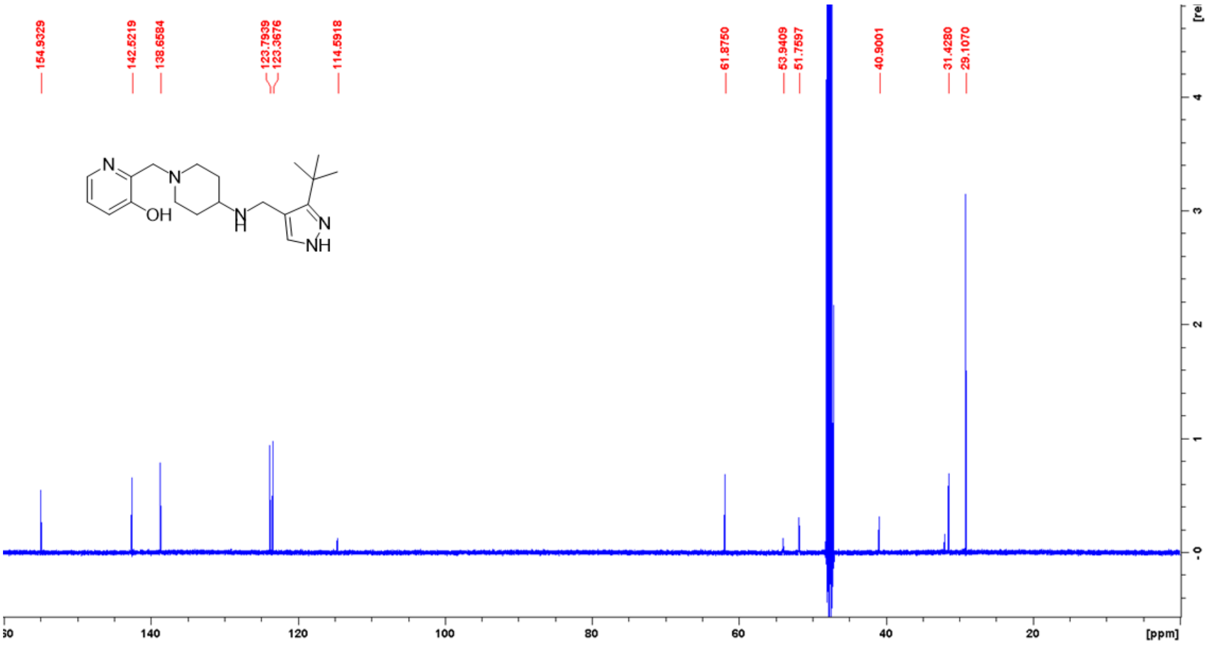

### [1]H and [13]C NMR spectra of compound 28 in MeOD

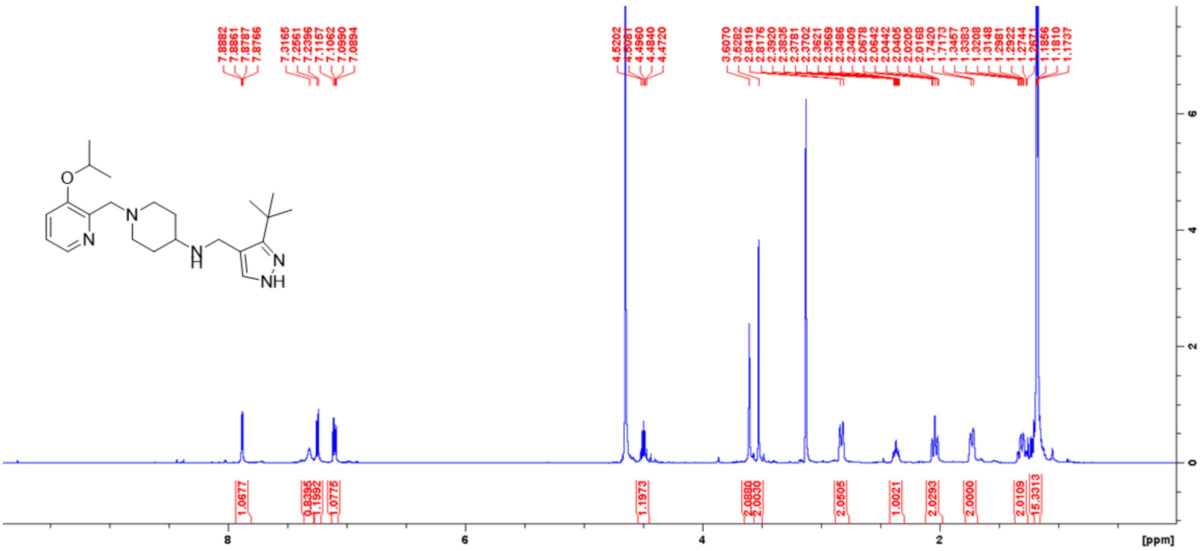


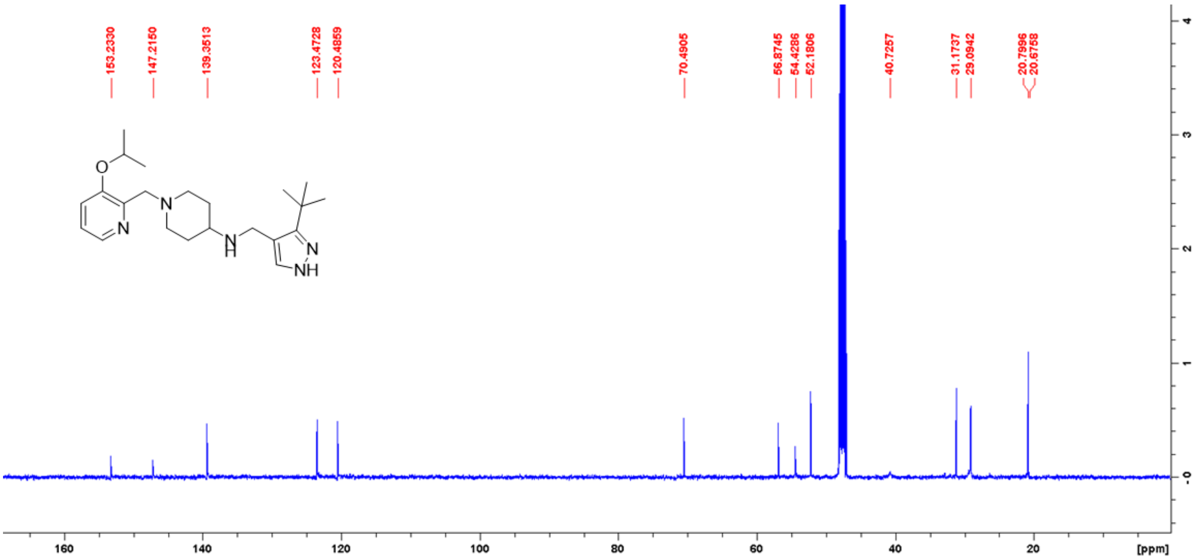

### $^{1}H$ and $^{13}C$ NMR spectra of compound 29 in MeOD

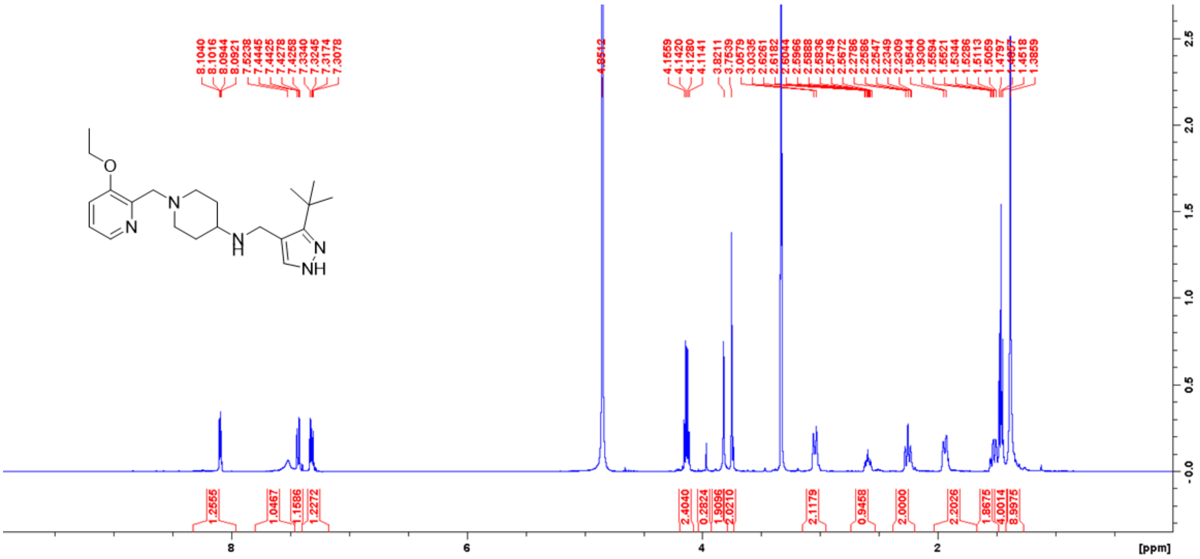


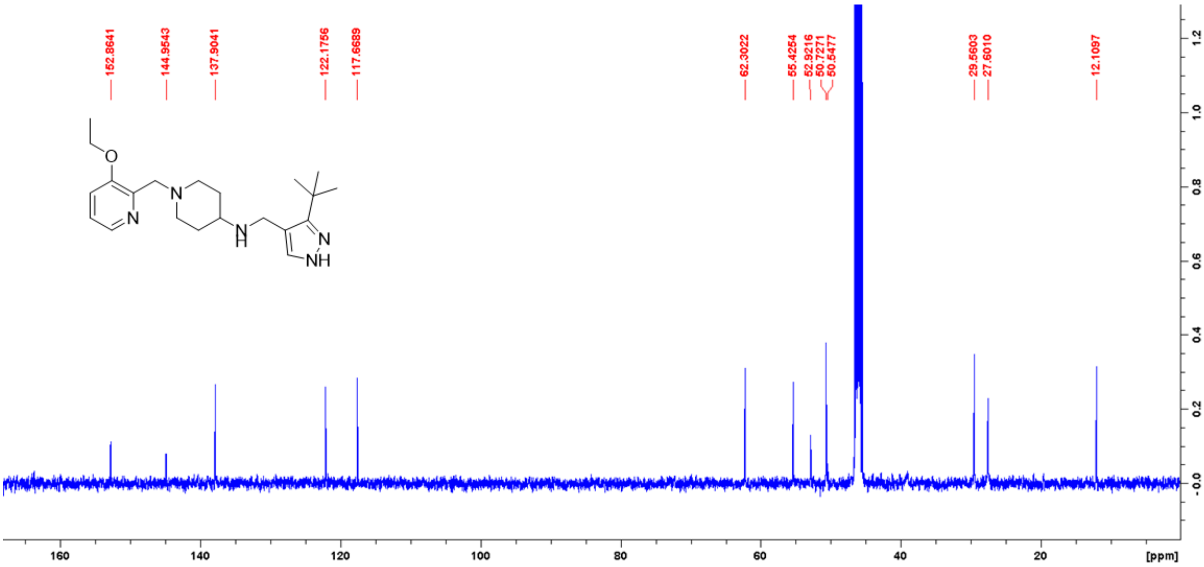

### [1]H and [13]C NMR spectra of compound 30 in MeOD

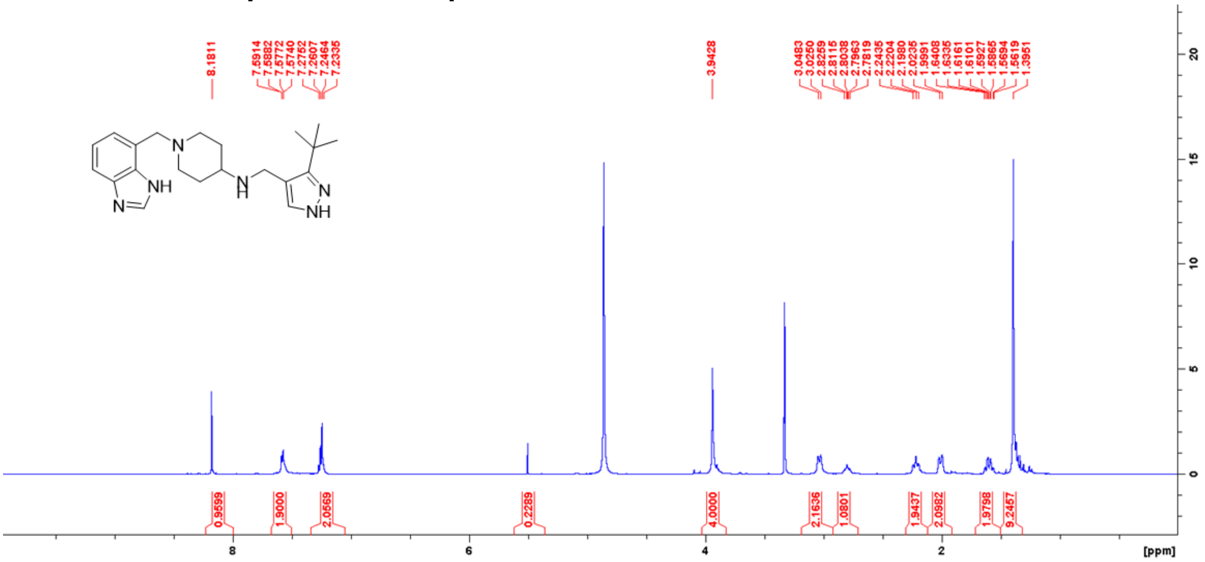

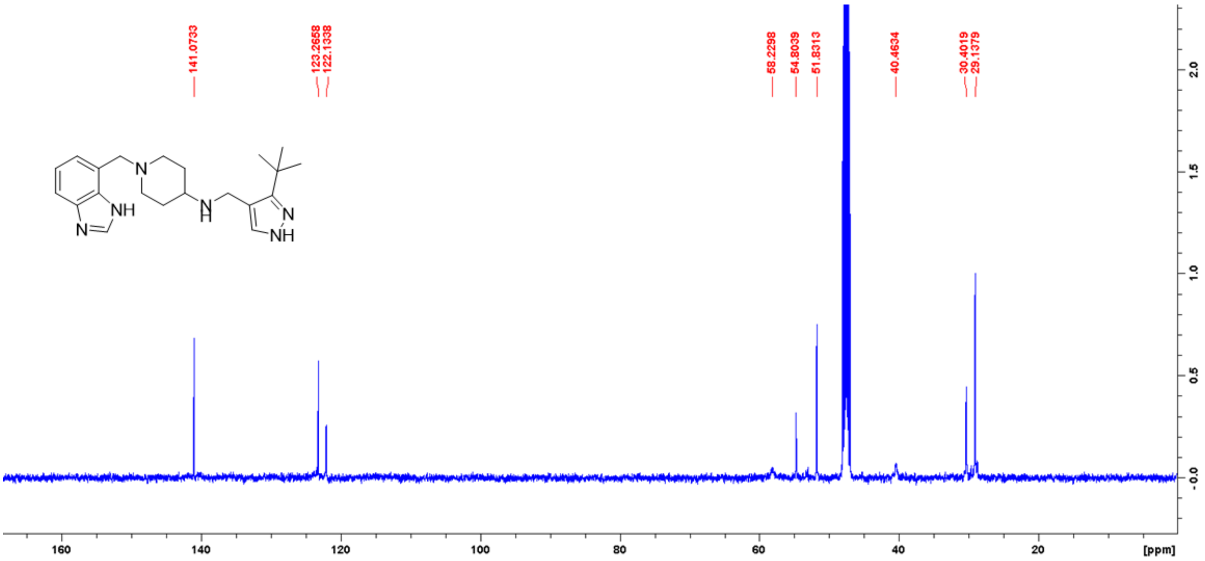

### $^{1}H$ and $^{13}C$ NMR spectra of compound 31 in MeOD

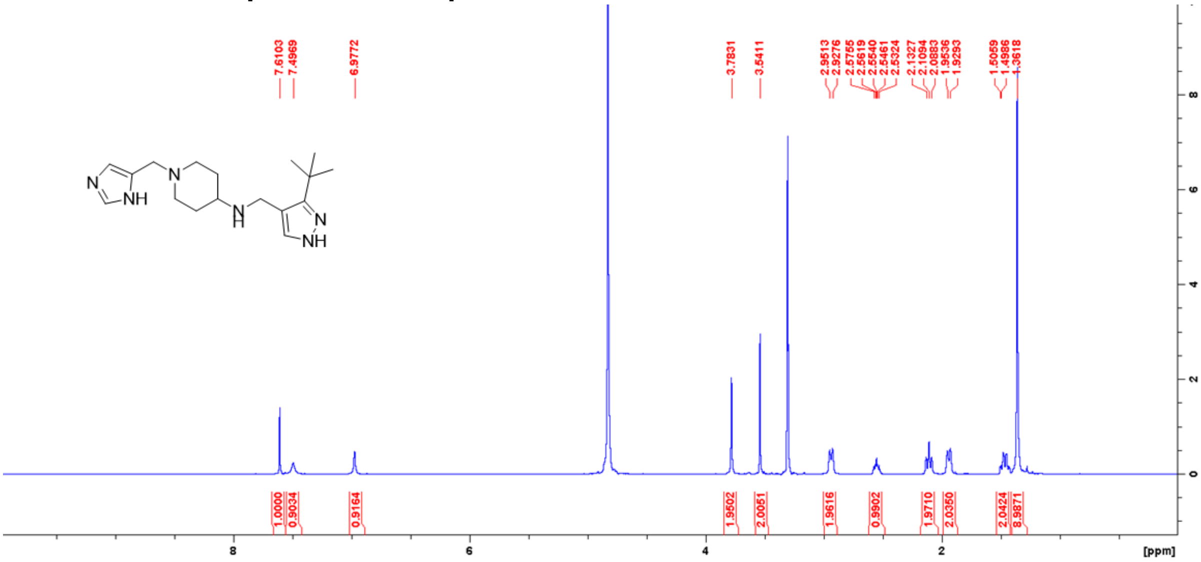


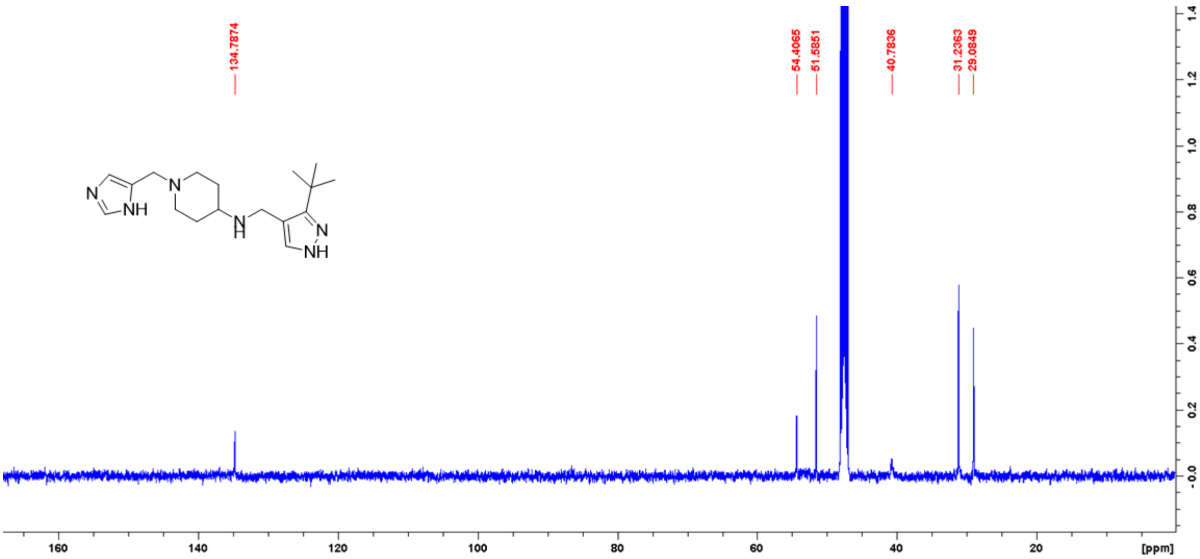

**$^{1}$H and $^{13}$C NMR spectra of compound 32 in MeOD**

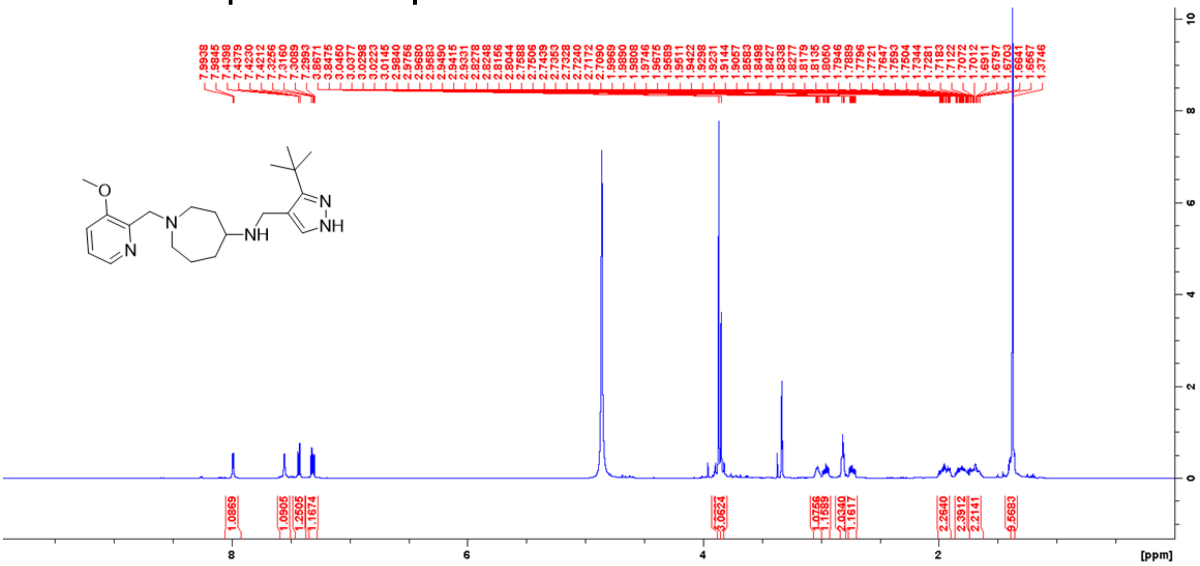


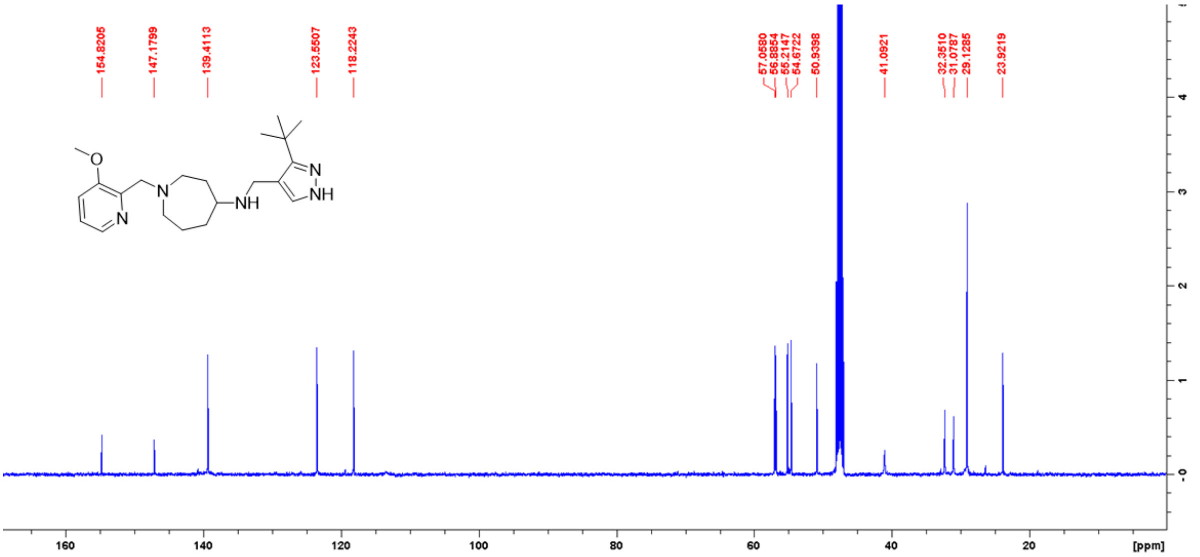
